# Most cancers carry a substantial deleterious load due to Hill-Robertson interference

**DOI:** 10.1101/764340

**Authors:** Susanne Tilk, Christina Curtis, Dmitri A Petrov, Christopher D McFarland

## Abstract

Cancer genomes exhibit surprisingly weak signatures of negative selection^1,2^. This may be because selective pressures are relaxed or because genome-wide linkage prevents deleterious mutations from being removed (Hill-Robertson interference)^3^. By stratifying tumors by their genome-wide mutational burden, we observe negative selection (dN/dS ~ 0.47) in low mutational burden tumors, while remaining cancers exhibit dN/dS ratios ~1. This suggests that most tumors do not remove deleterious passengers. To buffer against deleterious passengers, tumors upregulate heat shock pathways as their mutational burden increases. Finally, evolutionary modeling finds that Hill-Robertson interference alone can reproduce patterns of attenuated selection and estimates the total fitness cost of passengers to be 40% per cell on average. Collectively, our findings suggest that the lack of observed negative selection in most tumors is not due to relaxed selective pressures, but rather the inability of selection to remove deleterious mutations in the presence of genome-wide linkage.

## Introduction

Tumor progression is an evolutionary process acting on somatic cells within the body. These cells acquire mutations over time that can alter cellular fitness by either increasing or decreasing the rates of cell division and/or cell death. Mutations which increase cellular fitness (drivers) are observed in cancer genomes more frequently because natural selection enriches their prevalence within the tumor population^1,2^. This increased prevalence of mutations across patients within specific genes is used to identify driver genes. Conversely, mutations that decrease cellular fitness (deleterious passengers) are expected to be observed less frequently. This enrichment or depletion is often measured by comparing the expected number of nonsynonymous mutations (dN) within a region of the genome to the expected number of synonymous mutations (dS), which are presumed to be neutral. This ratio, dN/dS, is expected to be below 1 when the majority of nonsynonymous mutations are deleterious and removed by natural selection, be approximately 1 when all nonsynonymous mutations are neutral, and can be greater than 1 when a substantial proportion of nonsynonymous mutations are advantageous.

Two recent analyses of dN/dS patterns in cancer genomes found that for most non-driver genes dN/dS is ~1 and that only 0.1 - 0.4% of genes exhibit detectable negative selection (dN/dS < 1)^1,2^.This differs substantially from patterns in human germline evolution where most genes show signatures of negative selection (dN/dS ~ 0.4)^1^. Two explanations for this difference have been posited. First, the vast majority of nonsynonymous mutations may not be deleterious in somatic cellular evolution despite their deleterious effects on the organism. While most genes may be critical for proper organismal development and multicellular functioning, they may not be essential for clonal tumor growth. In this hypothesis, negative selection (dN/dS < 1) should be observed only within essential genes and absent elsewhere (dN/dS ~ 1). While appealing in principle, most germline selection against nonsynonymous variants appears to be driven by protein misfolding toxicity^4,5^, in addition to gene essentiality. These damaging folding effects ought to persist in somatic evolution.

A second hypothesis is that even though many nonsynonymous mutations are deleterious in somatic cells, natural selection fails to remove them. One possible reason for this inefficiency is the unique challenge of evolving without recombination. Unlike sexually-recombining germline evolution, tumors must evolve under genome-wide linkage that creates interference between mutations, known as-Hill-Robertson interference, which reduces the efficiency of natural selection^3^. Without recombination to link and unlink combinations of mutations, natural selection must act on entire genomes — not individual mutations — and select for clones with combinations of mutations of better aggregate fitness. Thus, advantageous drivers may not fix in the population, if they arise on an unfit background, and conversely, deleterious passengers can fix, if they arise on fit backgrounds.

The inability of asexuals to eliminate deleterious passengers is driven by two Hill-Robertson interference processes: *hitchhiking* and *Muller’s ratchet* (Fig. 1A). Hitchhiking occurs when a strong driver arises within a clone already harboring several passengers. Because these passengers cannot be unlinked from the driver under selection, they are carried with the driver to a greater frequency in the population. Muller’s ratchet is a process where deleterious mutations continually accrue within different clones in the population until natural selection is overwhelmed. Whenever the fittest clone in an asexual population is lost through genetic drift, the maximum fitness of the population declines to the next most fit clone (Fig. 1A). The rate of hitchhiking and Muller’s ratchet both increase with the genome-wide mutation rate^6,7^. Therefore, the second hypothesis predicts that selection against deleterious passengers should be more efficient (dN/dS < 1) in tumors with lower mutational burdens.

**Figure 1.**
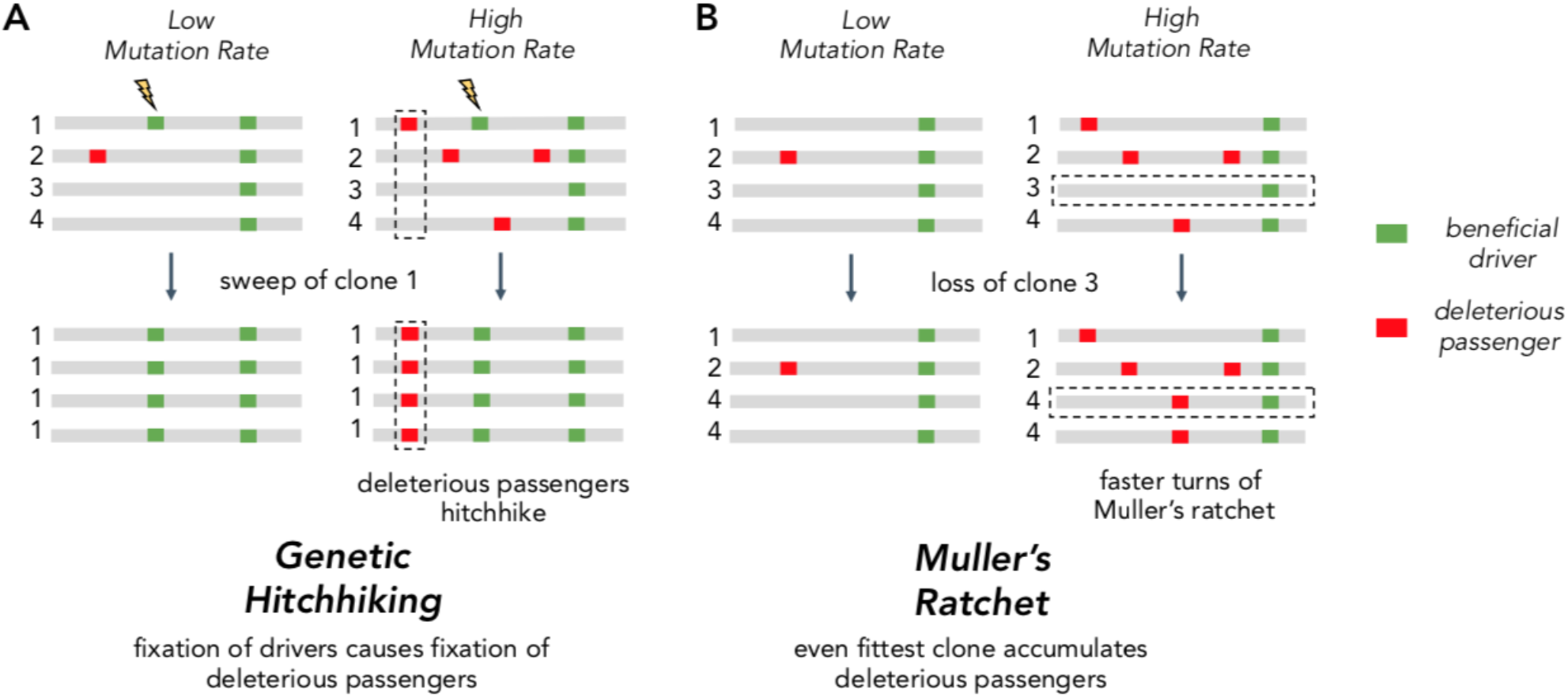
Two Hill-Robertson interference processes that accumulate deleterious mutations at high mutation rates. **(A) Genetic hitchhiking.** Each number identifies a different segment of a clone genome within a tumor. *De novo* beneficial driver mutations that arise in a clone can drive other mutations (passengers) in the clone to high frequencies (black dotted column). If the passenger is deleterious, both beneficial drivers and deleterious passengers can accumulate. **(B) Muller’s ratchet.** As the mutation rate within a tumor increases, deleterious passengers accumulate on more clones. If the fittest clone within the tumor is lost through genetic drift (black dotted row), the overall fitness of the population will decline.

Here, we leverage the 10,000-fold variation in tumor mutational burden across 50 cancer types to quantify the extent that selection attenuates, and thus becomes more inefficient, as the mutational burden increases. Using dN/dS, we find that selection against deleterious passengers and in favor of advantageous drivers is most efficient in low mutational burden cancers. Furthermore, low mutational burden cancers exhibit efficient selection across cancer subtypes, as well as within sub-clonal mutations, homozygous mutations, somatic copy-number alterations, and essential genes. Additionally, high-mutational burden tumors appear to mitigate this deleterious load by upregulating protein folding and degradation machinery. Finally, using evolutionary modeling, we find that Hill-Robertson interference alone can in principle explain these observed patterns of selection. Modeling predicts that most cancers carry a substantial deleterious burden (~40%) that necessitates the acquisition of multiple strong drivers (~5) in malignancies that together provide a benefit of ~130%. Collectively, these results explain why signatures of selection are largely absent in cancers with elevated mutational burdens and indicate that the vast majority of tumors harbor a large mutational load.

## Results

### Null models of mutagenesis in cancer

Mutational processes in cancer are heterogeneous, which can bias dN/dS estimates of selective pressures. dN/dS overcomes this issue by dividing observed mutation counts by what is expected under neutral evolution using null models. These null models must account for mutational biases that are often specific to cancer types and genomic regions.

To ensure our dN/dS calculations are robust and reproducible, we applied two different methods to account for mutational biases. The first approach uses a previously established parametric mutational model (dNdScv) that explicitly estimates the background mutational bias of each gene in its calculation of dN/dS^1^. The second approach uses a permutation-based, nonparametric (parameter-free) estimation of dN/dS. In this approach, every observed mutation is permuted while preserving the gene, patient samples, specific base change (e.g. A>T) and its tri-nucleotide context. Note that permutations do not preserve the codon position of a mutation and thus can change its protein coding effect (nonsynonymous vs synonymous). The permutations are then tallied for both nonsynonymous *d*_N_^(permuted)^ and synonymous *d*_S_^(permuted)^ substitutions (Fig. S1) and used as expected proportional values for the observed number of nonsynonymous *d*_N_^(observed)^ (or simply *d*_N_) and synonymous *d*_S_^(observed)^ (*d*_S_) mutations in the absence of selection. The unbiased effects of selection on a gene, dN/dS, is then:

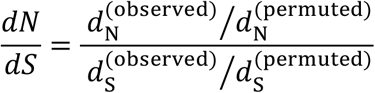

For all cancer types and patient samples, *P*-values and confidence intervals are determined by bootstrapping patient samples. Note that this permutation procedure will account for gene and tumor-level mutational biases (e.g. neighboring bases^9^, transcription-coupled repair, S phase timing^10^, mutator phenotypes) and their covariation. We confirmed that this approach accurately measures selection even in the presence of simulated mutational biases (Methods, Fig. S2A) as well as variation in gene length (Fig. S3). In addition, this approach also reliably measures the absence of selection (dN/dS = 1) in weakly expressed genes (Fig. S2C).

We find that both the parametric and nonparametric approaches identify similar patterns of selection (Fig. 2A and Fig. S3). Since parametric mutational models can become very complex in cancer (exceeding 5,000 parameters in some cases^1,8^), we elected to use the non-parametric approach, which makes fewer assumptions about underlying mutational processes, in subsequent calculations of dN/dS.

**Figure 2.**
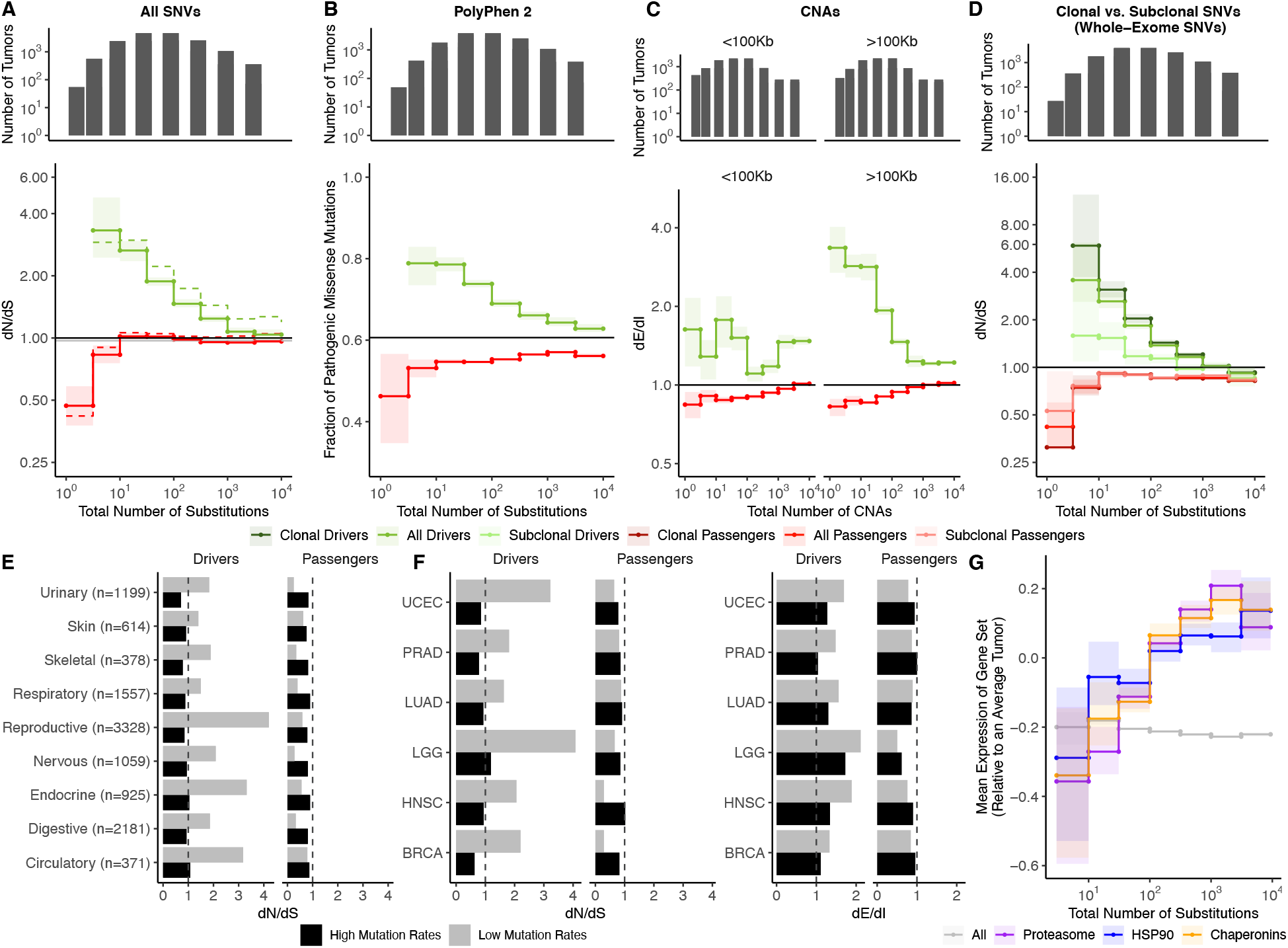
Attenuation of selection and increased protein folding stress in high mutation load tumors. **(A)** *dN/dS* of passenger (red) and driver (green) gene sets within 11,808 tumors (ICGC and TCGA) stratified by total number of substitutions present in the tumor (*d*_N_^(observed)^ + *d*_S_^(observed)^). *dN/dS* is calculated with error bars using a permutation based null model (solid line) and dNdScv (dashed). A *dN/dS* of 1 (solid black line) is expected under neutrality. Solid gray line denotes pan-cancer genomewide *dN/dS.* **(B)** Fraction of pathogenic missense mutations, annotated by PolyPhen2, in the same driver and passenger gene sets also stratified by total number of substitutions. Black line denotes the pathogenic fraction of missense mutations across the entire human genome. **(C)** Breakpoint frequency of CNAs that reside within exonic (dE) to intergenic (dI) regions within putative driver and passenger gene sets (identified by GISTIC 2.0, Methods) in tumors stratified by the total number of CNAs present in each tumor and separated by CNA length. Solid black line of 1 denotes values expected under neutrality. **(D)** *dN/dS* of clonal (VAF > 0.2; darker colors) and subclonal (VAF < 0.2; lighter colors) passenger and driver gene sets in tumors stratified by the total number of substitutions. A *dN/dS* of 1 (solid black line) is expected under neutrality. **(A-D)** Histogram counts of tumors within mutational burden bins are shown in the top panels. **(E)** Driver and passenger *dN/dS* values of the highest and lowest defined mutational burden bin in broad anatomical sub-categories. **(F)** Same as **(E)**, except for all specific cancer subtypes with ≥500 samples. **(G)** Z-scores of median gene expression within all genes, HSP90, Chaperonin and Proteasome gene sets averaged across patients (relative to an average tumor) stratified by the total number of substitutions. All shaded error bars are 95% confidence intervals determined by bootstrap sampling.

### Attenuation of selection in drivers and passengers for elevated mutational burden tumors

We estimated dN/dS patterns in both driver and passenger gene sets across 11,808 tumors from TCGA (whole-exome) and ICGC (whole-genome) aggregated over 50 cancer types (Methods). We used the following four mutational tallies as a proxy for the genome-wide mutation rate: (1) the total number of mutations or tumor mutational burden (TMB) (2) the total number of observed substitutions in both synonymous and nonsynonymous sites (*d*_N_ + *d*_S_) (Fig. 1), and (3) the total number of mutations in intergenic, and (4) intronic regions. All estimates are strongly correlated (*R^2^ > 0.97,* Fig. S4).

In principle, only the last two tallies — the number of substitutions in intergenic or intronic regions — are orthogonal to dN/dS, and least likely to be biased by selection. However, these measures can only be applied to whole-genome datasets, which constitute only 15% of sequenced samples. Therefore, for most of the analyses, we used the second measure (*d*_N_ + *d*_S_) to define mutational burden, while being cognizant that the analysis could be complicated by the fact that the same mutation tallies are used for both the x-axis (*d*_N_ + *d*_S_) and y-axis (*dN/dS*). We note that this interdependence leads to a slight underestimation of the degree of purifying selection, rendering our analysis conservative (Fig. S5, Methods).

To quantify the extent that selection attenuates as the mutational burden increases, we stratified tumors into bins based on their total number of substitutions on a log scale. For each bin of tumors, we pooled all of the variants together and estimated dN/dS jointly. Consistent with the inefficient selection model, whereby selection fails to eliminate deleterious mutations in high mutational burden tumors, we observe pervasive selection against passengers exclusively in tumors with low mutational burdens (dN/dS ~ 0.47 in tumors with mutational burden ≤ 3, while dN/dS ~ 0.95 in tumors with mutational burden > 10, Fig. 2A). We observed little negative selection in passengers when aggregating tumors across all mutational burdens (dN/dS ~ 0.97), which is broadly similar to previous estimates^1,2,8,11^.

We confirmed that negative selection on passengers is specific to low mutational burden tumors and not biased by small sample sizes (Fig. S2B). We randomly sampled passengers from high mutational burden tumors (>10 substitutions) 1000 times using the same bin sizes in Fig. 2A and calculated dN/dS. Within the smallest bin size (N=130 SNVs), negative selection on passengers sampled from high mutational burden tumors was absent (average dN/dS ~ 1.02) compared to observed dN/dS in low mutational burden tumors (dN/dS ~ 0.47; *p* < 2.2^−16^). In fact, only 0.6% of randomly sampled sets of sites had similar signals of negative selection (dN/dS < 0.47).

Also consistent with the inefficient selection model, drivers exhibit a similar but opposing trend of attenuated selection at elevated mutational burdens (dN/dS ~ 3.3 when mutational burden ≤ 3 and gradually declines to ~1.5 when mutational burden > 100). This pattern is not specific to drivers that are oncogenes or tumor suppressors (Fig. S6). While the attenuation of selection against passengers in higher mutational burden tumors is a novel discovery, this pattern among drivers has been reported previously^1^.

Furthermore, we confirmed that these patterns are robust to the choices that we made in our analysis pipeline. These include the: (1) somatic mutation calling algorithm (Mutect2 and MC3 SNP calls^12^, Fig. S3B), (2) dataset (TCGA^13^, ICGC^14^, COSMIC^15^ and an additional independent validation cohort; Fig. S3B and Fig. S3D), (3) effects of germline SNP contamination (Fig. S7), (4) choice of driver gene set (Bailey et al^16^, IntOGen^17^, and COSMIC^15^, Fig. S3B and Fig. S8), (5) mutational burden metric (Fig. S3A), (6) differences in tumor purity and thresholding (Fig. S9), and (7) null model of mutagenesis (dNdScv, Fig. S3C & S10)^1^ (Methods).

If negative selection is more pronounced in low mutational burden tumors, then the nonsynonymous mutations observed should also be less functionally consequential. By annotating the functional effect of all missense mutations using PolyPhen2^18^ (Fig 2B), we indeed find that observed nonsynonymous passengers are less damaging in low mutational burden cancers. Similarly, driver mutations become less functionally consequential as mutational burden increases, as expected for mutations experiencing inefficient positive selection (Fig 2B). Together these two trends provide additional and orthogonal evidence that selective forces on nonsynonymous mutations are more efficient in low mutational burden cancers.

Since all mutational types experience Hill-Robertson interference, attenuated selection should also persist in Copy Number Alterations (CNAs). We used two previously-published statistics to quantify selection in CNAs: Breakpoint Frequency^19^ and Fractional Overlap^20^. For both measures, we compare the number of CNAs that either terminate (Breakpoint Frequency) within or partially overlap (Fractional Overlap) **E**xonic regions of the genome relative to non-coding (**I**ntergenic and **I**ntronic) regions (*dE/dI*, See Methods). Like dN/dS, dE/dI is expected to be <1 in genomic regions experiencing negative selection, >1 in regions experiencing positive selection (e.g. driver genes), and approximately 1 when selection is absent or inefficient (Fig. S23). Using dE/dI, we observe attenuating selection in both driver and passenger CNAs as the total number of CNAs increases for both Breakpoint Frequency (Fig. 2C) and Fractional Overlap (Fig. S11). While CNAs of all lengths experience attenuated selection, CNAs longer than the average gene length (>100 KB) experience greater selective pressures in drivers (*p* < 10^-4^).

Collectively, these results suggest that tumors with elevated mutational burdens carry a substantial deleterious load. Since nonsynonymous mutations are thought to be primarily deleterious by inducing protein misfolding^4,5^, we tested whether an increase in the number of passenger mutations in tumors would lead to elevated protein folding stress, and, in turn, drive the upregulation of heat shock and protein degradation^21^ pathways in cancer^22^. Indeed, gene expression of HSP90, Chaperonins, and the Proteasome does increase across the whole range of SNV (weighted R^2^ of 0.83, 0.77, and 0.75 respectively) and CNA burdens (weighted R^2^ of 0.78, 0.87 and 0.84, respectively) (Fig. 2G and S22). This trend persists across cancer types for SNVs and CNAs (Fig. S22). Importantly, expression of these gene sets increases across the whole range of mutational burdens, even after the dN/dS of passengers approaches 1. This result presents additional evidence that passengers continue to impart a substantial cost to cancer cells, even in high mutational burden tumors.

### Strong selection in low mutational burden tumors cannot be explained by mutational timing, gene function, or tumor type

We next tested alternative hypotheses to the inefficient selection model. We considered the possibility that selection is strong only during normal tissue development, but absent after cells have transformed to malignancy. This would disproportionately affect low mutational burden tumors, as a greater proportion of their mutations arise prior to tumor transformation. If true, then attenuated selection should be absent in sub-clonal mutations, which must arise during tumor growth. However, selection clearly attenuates with increasing mutational burden for the subset of likely sub-clonal mutations with Variant Allele Frequency (VAF) below 20% (Fig. 2D & S12). Although selection attenuates in drivers and passengers in both sub-clonal and clonal mutations, selection is weaker in both drivers and passengers with lower VAFs. Weaker efficiency of selection among less frequent variants is expected under a range of population genetic models^23^ and especially so in rapidly-expanding, spatially-constrained cancers^24^. In addition, heterozygous mutations, to the extent they are only partially-dominant^25^, are also expected to exhibit lower VAFs and experience weaker selection.

Next, we considered and rejected the possibility that attenuated selection is limited to particular types of genes. We first annotated our observed mutations by different functional categories and Gene Ontology (GO) terms^26^ and find that negative selection is not specific to any particular gene functional category, and specifically not limited to essential or housekeeping genes — a key prediction of the ‘weak selection’ model^1^ (Fig. S13, *p* < 0.05, Wilcoxon signed-rank test).

Finally, we found that these patterns of attenuated selection persist across cancer subtypes for both SNVs and CNAs. We calculated dN/dS in tumors grouped by nine broad anatomical sub-categories (e.g. neuronal) and 50 subtype classifications ^27^(Fig. 2E-F). We find that patterns of attenuated selection in SNVs persists in the broad and specific (drivers *p* = 1.4 × 10^-5^, passengers *p* = 1.3 × 10^-2^, Wilcoxon signed-rank test; Fig. S14) classification schemes. Furthermore, dE/dI measurements of CNAs exhibit these same patterns of selection in broad (Fig. S15) and specific subtypes (Fig. 2F; drivers *p* < 10^-6^ and passengers *p* = 7.3 × 10^-4^).

Collectively, these results strongly support the inefficient selection model and argue that the observed patterns must be due to a universal force in tumor evolution. We find that selection consistently attenuates in both drivers and passengers across all cancers as mutational burden increases.

### Evolutionary modeling estimates the fitness effects of drivers and passengers, and rate of Hill-Robertson interference processes

We next tested whether Hill-Robertson interference – a process where selection becomes inefficient due to interference between linked mutations with competing fitness effects – alone can generate these patterns of attenuated selection. Specifically, we modeled tumor progression as a simple evolutionary process with advantageous drivers and deleterious passengers. We then used Approximate Bayesian Computation (ABC) to compare these simulations to observed data and infer the mean fitness effects of drivers and passengers.

Our previously-developed evolutionary simulations model a well-mixed population of tumor cells that can randomly acquire advantageous drivers and deleterious passengers during cell division^28^. The product of the individual fitness effects of these mutations determines the relative birth and death rate of each cell, which in turn dictates the population size *N* of the tumor. If the population size of a tumor progresses to malignancy (*N* > 1,000,000) within a human lifetime (≤100 years), the accrued mutations and patient age are recorded. The mutation rate of each simulated tumor is randomly sampled from a broad range (10^−12^ to 10^−7^ mutations • nucleotide^-1^ • generation^-1^, Methods). Although this model ignores a great deal of known tumor biology, we believe it constitutes the simplest evolutionary model that could possibly recapitulate observed selection for drivers and against passengers. Our question is not whether this model is correct in all details but rather whether even such a simple model can generate quantitatively similar patterns as observed in the data with sensible values of mutation rates and selection coefficients.

Figure 3A illustrates the ABC procedure. To compare our model to observed data, we simulated an exponential distribution of fitness effects with mean fitness values that spanned a broad range (10^−2^ - 10^0^ for driver and 10^−4^ - 10^−2^ for passengers, Methods). We summarized observed and simulated data using statistics that capture three relationships: (i) the dependence of driver and passenger dN/dS rates on mutational burden, (ii) the rate of cancer age-incidence (SEERs database^29^), and (iii) the distribution of mutational burdens (summary statistics of (ii) and (iii) were based on theoretical parametric models^30^, Methods, Fig. S16 & S17). We then inferred the posterior probability distribution of mean driver fitness benefit and mean passenger fitness cost using a rejection algorithm that we validated using leave-one-out Cross Validation (Methods, Fig. S18).

**Figure 3.**
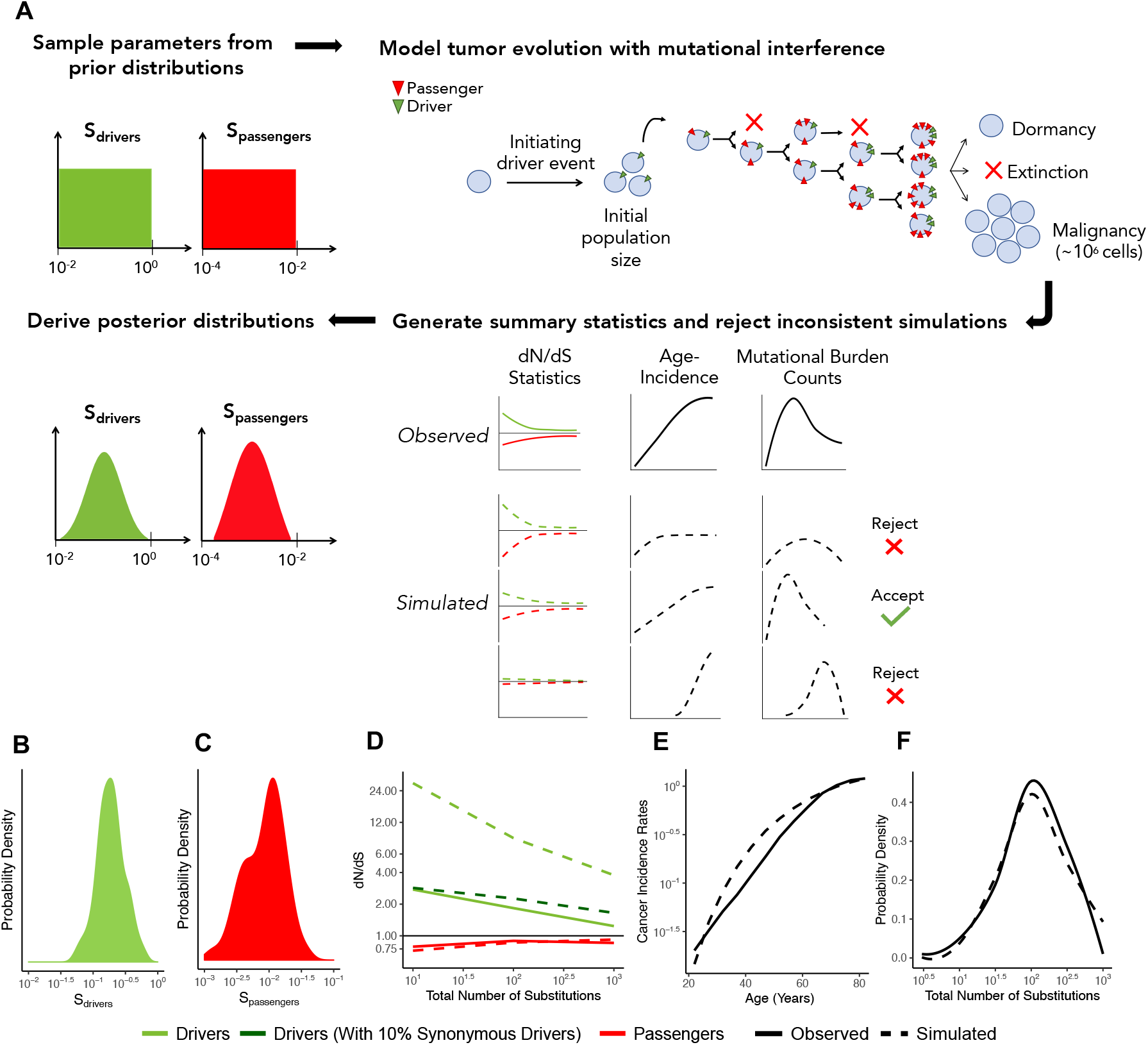
ABC procedure estimates the strength of selection in passengers and drivers. **(A)**Schematic overview of the ABC procedure used. A model of tumor evolution with genome-wide linkage contains two parameters — *s*_drivers_ (mean fitness benefit of drivers) and *s*_passengers_ (mean fitness cost of passengers) — sampled over broad prior distributions of values. Simulations begin with an initiating driver event that establishes the initial population size of the tumor. The birth rate of each individual cell within the tumor is determined by the total accumulated fitness effects of drivers and passengers. If the final population size of the tumor exceeds one million cells within a human lifetime (100 years), patient age and accrued mutations are recorded. Summary statistics of four relationships are used to compare simulations to observed data: (i) *dN/dS* rates of drivers and (ii) passengers across mutational burden, (iii) rates of cancer incidence versus age, and (iv) the distribution of mutational burdens. Simulations that excessively deviate from observed data are rejected (Methods). **(B-C)** Inferred posterior probability distributions of *s*_drivers_ and *s*_Passengers_. The Maximum Likelihood Estimate (MLE) of *s*_drivers_ is 18.8% (green, 95% CI [13.3, 32.7]), and the MLE of *s*_passengers_ is 0.96% (green, 95% CI [0.28, 3.6%]). **(D-F)** Comparison of the summary statistics of the best-fitting simulations (MLE parameters, dashed lines) to observed data (solid lines). **(D)** dN/dS rates of passengers (red) and drivers (light green) for simulated and observed data versus mutational burden. A model where 10% of synonymous mutations within drivers experience positive selection (dark green) was also considered. **(E)** Cancer incidence rates for patients above 20 years of age. **(F)** Distribution of the mutational burdens of tumors. **(G)** Fine scale comparison of *dN/dS* within drivers and passengers in observed data compared to one million simulated tumors using the MLE estimates of *s*_drivers_ and *s*_passengers_.

Using this approach, the Maximum Likelihood Estimate (MLE) of mean driver fitness benefit is 18.8% (Fig. 3B), while the MLE of passenger mean fitness cost is 0.96% (Fig. 3C). Simulations with these MLE values agree well with all observed data (Fig. 3D-F, Pearson’s *R* = 0.95, 0.80, 0.99, 0.97 for driver dN/dS, passenger dN/dS, Age-Incidence, and Mutational Burden respectively).

While Hill-Robertson interference alone explains dN/dS rates in the passengers well, the simulations most consistent with observed data still exhibited consistently higher dN/dS rates in drivers (Fig. 3D). We tested whether positive selection on synonymous mutations within driver genes could explain this discrepancy. Indeed, we find that a model incorporating synonymous drivers agrees modestly better with observed statistics (*p* = 0.043, ABC posterior probability). The best-fitting model predicts that ~10% of synonymous mutations within driver genes experience positive selection, which is consistent with previous estimates for human oncogenes^31^ (Methods, Fig. 3D, S19). Furthermore, we observe additional evidence of selection and codon bias in synonymous drivers exclusive to low mutational burdens (TCGA samples, Methods, Fig. S19).

Finally, our simulations demonstrate that deleterious passengers are necessary and sufficient to explain the quantitative shape of dN/dS curves – both the steep attenuation in passengers and the gradual attenuation in drivers (Fig. 3G). In the Supplementary Note, we discuss several simple models of progression with neutral passengers that cannot explain the dN/dS curves of drivers (e.g. such as a 5-hit model or selection bias of tumors with a low mutational burden).

Our results indicate that rapid adaptation through natural selection – acting on entire genomes, rather than individual mutations – is pervasive in all tumors, including those with elevated mutational burdens. Given the quantity of drivers and passengers observed in a typical cancer (TCGA), our model implies that cancer cells are in total ~90% fitter than normal tissues (130% total benefit of drivers, 40% total cost of passengers). A median of five drivers each of which has a mean benefit of ~19% accumulate per tumor in these simulations – also consistent with estimates from age-incidence curves^29^, known hallmarks of cancer^37^, and estimates of the selective benefit of individual drivers^34^. Lastly, the mutation rates of tumors that could progress to cancer in our model also recapitulate observed mutation rates in human cancer^38^ (median 3.7 x 10^-9^, 95% Interval 1.1 x 10^-10^ - 8.2 x 10^-8^, Fig. S20).

Most notably, under our modeling assumptions, all passengers together confer a fitness cost of ~40% per tumor. While this collective burden appears large, the individual fitness effects of accumulated passengers in these simulations (mean 0.8%) are similar to observed fitness costs in cancer cell lines (1 - 3%)^39^ and the human germline (0.5%)^40^. Note that in our model, these passengers accumulated primarily via Muller’s Ratchet, while only ~14% accumulated via hitchhiking (inferred using population genetics theory^28^ and MLE fitness effects, Methods, Fig. S21). These results suggest that Hill-Robertson interference is a plausible model for the empirical patterns of attenuated selection with mutational burden observed in the data.

## Discussion

Here we argue that signals of selection are largely absent in cancer because of the inefficiency of selection and not because of weakened selective pressures. In low mutational burden tumors (≤ 10 total substitutions per tumor), increased selection for drivers and against passengers is observed and ubiquitous: in SNVs and CNAs; in heterozygous, homozygous, clonal, and sub-clonal mutations; and in mutations predicted to be functionally consequential. These trends are not specific to essential or housekeeping genes. Importantly, these patterns persist across broad and specific tumor subtypes. Collectively, these results suggest that inefficient selection is generic to tumor evolution and that deleterious load is a nearly-universal hallmark of cancer.

Importantly, these patterns of selection are missed when dN/dS rates are not stratified by mutational burden. Since only 0.1% of mutations in TCGA and ICGC reside within low mutational burden tumors (4% of all tumors, *N*=563), the dN/dS of passengers at low mutational burdens (~0.47 - 0.8) do not appreciably alter the pan-cancer dN/dS of passengers (0.97 in our study, 0.82 — 0.98 in^1,2,8,11^). In fact, the power to detect negative selection on passengers at low mutational burdens is only possible by aggregating all mutations within these tumors and estimating dN/dS jointly. Thus, we believe that low mutational burden tumors are uniquely valuable for identifying genes and pathways under positive and negative selection. While only 4% of tumors exhibit substantial negative selection, selection in drivers, selection on CNAs, and expression patterns of chaperones and proteasome components all show a continuous response to deleterious passenger load across a broad range of mutational burdens. Collectively, this suggests that passengers continue to be deleterious even in high mutational burden tumors.

Using a simple evolutionary model, we show that Hill-Robertson Interference alone can explain this ubiquitous trend of attenuated selection in both drivers and passengers. dN/dS rates attenuate in drivers because the background fitness of a clone becomes more important than the fitness effects of an additional driver at elevated mutation rates. Furthermore, these simulations indicate that, despite dN/dS patterns approaching 1 in tumors with elevated mutational burdens, passengers are not effectively neutral (*Ns* > 1). Instead, passengers confer an individually-weak, but collectively-substantial fitness cost of ~40% that measurably impacts tumor progression. Because this simple evolutionary model does not explicitly incorporate many known aspects of tumor biology (e.g. haploinsufficiency, see Table S2), these fitness estimates are highly provisional. Nonetheless, we note that selection’s efficiency in cancer is further reduced when spatial constraints are considered^24^.

The functional explanation for why passengers in cancer are deleterious is unknown. In germline evolution, mutations are believed to be primarily deleterious because of protein misfolding^4,5^. Deleterious passengers in somatic cells should confer similar effects. Indeed, we find that elevated mutational burden tumors may buffer the cost of deleterious mutations by upregulating multiple heat-shock pathways. However, deleterious passengers may carry other costs to cancers or be buffered by additional mechanisms. Understanding and identifying how tumors manage this deleterious burden should identify new cancer vulnerabilities that enable new therapies and better target existing ones^41–43^.

## Acknowledgements

We thank Judith Frydman for her contribution on the heat shock response analysis, Monte Winslow for his contribution on cancer subtype analysis, Donate Weghorn for her contribution on the interdependence of *dN/dS* and mutational burden, Leonid Mirny, Grant Kinsler, Gabor Boross, Chuan Li, Alison Feder, Eliot Cowan and other members of the Petrov and Curtis labs for helpful comments and discussions. This work is supported by NIH grants T32-HG000044-21, E25-CA180993; the Director’s Pioneer Award DP1-CA238296 to C.C.; R01-CA207133, R35-GM118165, and R01-CA231253 to D.A.P.; and K99-CA226506 to C.D.M.

## References

### Methods & Supplementary Materials

#### Data Availability

Exonic, open-access SNV calls (WES) of 10,486 cancer patients in (The Cancer Genome Atlas) TCGA were downloaded from the Multi-Center Mutation Calling in Multiple Cancers (MC3) project^12^. This repository uses a consensus of seven mutation-calling algorithms. Whole-Genome Sequencing SNV calls (WGS) of 1,830 patients were downloaded from the ICGC data portal in November 2018^44^. Supplemental analyses on the effect of variant callers, SNVs from exome and whole genome wide screens were downloaded on October 2016 from the Catalog of Somatic Mutations in Cancer’s (COSMIC) Mutant Export Census^15^. Expression data of SNVs were downloaded from the Genotype-Tissue Expression (GTEx) project (v7 release)^45^. All CNAs were downloaded from the COSMIC database on June 2015^15^. Gene expression data compared to CNAs was downloaded from the COSMIC database on September 2019. To validate our findings, additional WES and WGS SNV calls were downloaded from cBioPortal from 1,786 treatment-naive, tumor-normal sample pairs across 17 studies of varying cancer types in February 2019. Formalin-Fixed Paraffin Embedded (FFPE) samples were removed. ^46,47,56^-^62,48–55^

#### Code Availability

All code for the simulations, associated theoretical analysis, and generation of summary statistics will be made publicly available under the open-source MIT License upon publication. Code for simulations of tumor growth with advantageous drivers and deleterious passengers is currently available at https://github.com/mirnylab/pdSim.

#### Mutation calling and quality controls

Mutations were downloaded from online repositories that have already invested heavily in quality control. Multiple data repositories were used to ensure reproducibility. Post-processing was minimal to avoid engendering a particular result, and only excluded sequencing samples obtained from cell lines, or studies that did not report synonymous variants. These exclusions are described in greater detail below.

#### Somatic Nucleotide Variants (SNVs)

Only consensus mutation calls from the PCAWG Consensus SNV-MNV caller were considered. Both missense and nonsense mutations are defined as nonsynonymous mutations. Frameshift, indels, and splice-site variants were not included in analyses. Samples without any synonymous or nonsynonymous mutations in either dataset were excluded. Note that there is no evidence of germline contamination by common SNPs (MAF > 5%) from 1,000 Genomes Project^63^ (v 2015 Aug) using ANNOVAR^64^ to annotate mutations in either datasets (Fig. S7). A final of 1,703 whole-genome and 10,152 whole-exome sequencing samples were used for the analyses in this paper. In SNV data collected from COSMIC, studies before 2010 that didn’t report silent mutations, and cell lines were removed from analysis. Whole-exome SNVs in TCGA were also called using Mutect2^65^ (Fig. S3B).

#### Defining tumor burden

We tested four different mutation burden metrics as a proxy for the genome-wide mutation rate: (1) the total number of observed mutations, (2) total number of substitutions in both synonymous and nonsynonymous sites (*d*_N_^(observed)^ + *d*_S_^(observed)^), (3) the total number of mutations in intergenic, and (4) intronic regions. Although only the last two definitions of mutational burden are completely independent to *dN/dS,* the vast majority of samples (10,152 vs 1,703) are derived from whole-exome data. We note that all mutation rates are strongly correlated to each other (R^2^ > 0.97). Because only d_*N*_ + d_*S*_ could be applied to WES data — the majority of samples — and all metrics worked equally-well, we primarily used d_*N*_ + d_*S*_ to measure mutational burden. Lastly, because dN/dS is undefined for tumors with no synonymous mutations, we necessarily excluded these samples. We also excluded samples with no nonsynonymous mutations so as to apply a symmetric filter on the data and because data quality may be compromised in these samples. Inclusion of samples with zero synonymous mutations or zero nonsynonymous mutations did not appreciably alter observed trends in the TCGA and ICGC datasets (Fig. S5D).

#### A Nonparametric Null Model of Mutagenesis to calculate dN/dS

We assume that for any particular tumor, mutation rates are constant across a gene for a particular tri-nucleotide context and base change (e.g. C > G). Our procedure is inspired by Constrained Marginal Models (or ‘edge switching’ in network analysis), whereby the marginal distributions of observations aggregated over known confounding variables are preserved under permutation to create a null distribution. In our application of this strategy, the marginal distributions of mutations (across tri-nucleotide context, base change, gene, and tumor) remain preserved – as they would be in a Constrained Marginal Model; however, we exhaustively consider every acceptable permutation of the data. Because our approach is highly-constrained, these permutations are exhaustively computable (median 36 alternatives per mutation). Thus, resampling is unnecessary.

Our null model presumes that all mutations of type *i,* defined by a tri-nucleotide context and base change, arise with probability *M*_igt_ within each gene *g* and tumor *t.* For each gene, we tally the total quantity of nonsynonymous mutations *N*_ig_ and synonymous mutations *S*_ig_. Suppose selection enriches or depletes nonsynonymous mutations within a gene and tumor by a rate *ω*_gt_. The expected number of nonsynonymous and synonymous mutations within a particular tumor and gene are 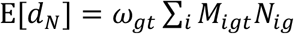 and 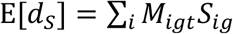 in the absence of selective pressures on synonymous mutations. As with the main text, *d*_N_ and *d*_N_^(observed)^ are used interchangeably. Although *M*_igt_ is unknown, *dN/dS* statistics attempt to infer selection nonetheless by noting that:

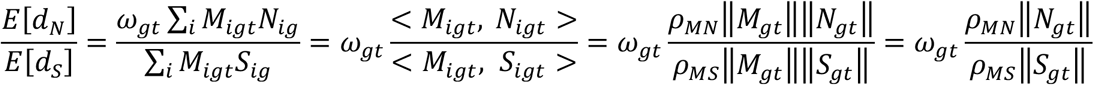

Note that *ρ*_AB_ =< *A,B* >/(||*A*||||*B*||) where 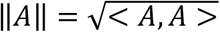 is the Pearson product-moment correlation coefficient. When *ρ*_MN_ ≈ *p*_ms_,

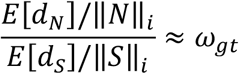

I.e. *dN/dS* is approximately equal to the selective pressures on nonsynonymous mutations when the accessible nonsynonymous and synonymous loci are properly accounted and when the correlation between mutational processes and nonsynonymous loci are roughly equivalent to the correlation between mutational processes and synonymous loci. Traditionally, this assumption was used to calculated *dN/dS.* To improve resolution of *dN/dS,* researchers have attempted to account for these correlations using sophisticated parametric models of *M*_igt_. An alternative statistical approach, however, is to treat these correlations as nuisance parameters.

Constrained Marginal Models permute observed data in all possible manners that preserve the underlying covariance structure of the data (e.g. *ρ*_MN_, *ρ*_MS_). In our particular case of this method, we note that by definition, 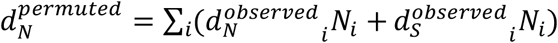. Thus:

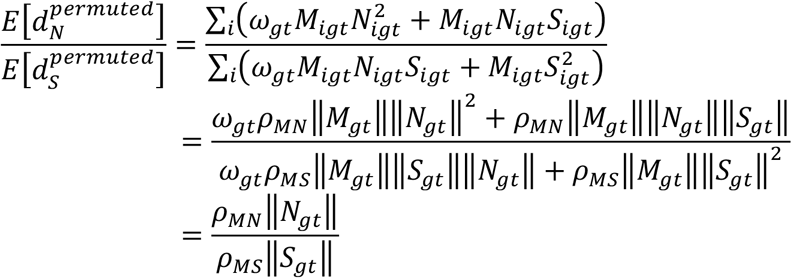

Hence, by dividing the observed mutations by all permutations, we eliminate the covariance of mutational processes with available loci and, thus, measure *ω*_gt_ directly for any particular gene-tumor combination without mutational bias.

Unfortunately, because of the log-sum inequality, mutational bias can arise once cohorts of genes and cohorts of tumor samples are binned. This problem is common to all *dN/dS* measures and is a consequence of the correlation of mutational biases with *selection* (i.e. < *M*_igt_, *ω_gt_* >) – not the correlation of mutational biases with one another, as these covariances are already accounted-for in a Constrained Marginal Model. For example, if tri-nucleotide biases covary linearly with gene-level biases, and are independent of tumor-level biases, then a parametric estimate of *M*_igt_ may deconstruct *M*_igt_ into *M_igt_ = f(i, g, t, p_ig_),* where *p_ig_* is the covariation of tri-nucleotide mutational biases with gene-level biases. Nonetheless, < *M*_igt_, ω_gt_ > ∝ < *p_ig_, ω_gt_ >* will still be ignored. Indeed, this covariation of mutational processes with selective forces is the focus of our current study: selection and genome-wide mutation rate are correlated (i.e. *∑_t_ *M*_igt_ω_gt_* ≠ 0) because of Hill-Robertson Interference. Hence, the level at which observed *d*_N_ values *d*_S_ are binned necessarily ignores covariation between mutational processes and selection (in addition to any variation of *ω*_gt_ within the bin). Another example of this binning challenge arises when positive and negative selection act on different regions of the same gene, which gene-level *dN/dS* binning can misinterpret as neutral evolution.

#### Validation of nonparametric null model

To confirm that our null model can accurately estimate *dN/dS* even in the presence of extreme tri-nucleotide mutational biases, we simulated artificial data where different COSMIC signatures^15^ (SBS Signatures 1-9, v3) contribute to all of the mutations. Permuted *d*_N_ and *d*_S_ tallies for each mutational context were simulated by randomly sampling 1,000 genes with the same mutational context. The fraction of permuted *d*_N_ and *d*_S_ tallies for each mutational context was used as weighted probabilities to derive observed *d*_N_ and *d*_S_ tallies. To simulate negative selection, *d*_N_ counts were randomly removed from each context at a rate 1 - *ω*_gt_ (e.g. a simulated ‘true’ *dN/dS* of 0.8 in a cohort of samples indicates a 20% chance of nonsynonymous mutations being removed in the samples). These simulated (true) rates were then compared to observed and permuted *d*_N_ and *d*_S_ tallies according to the *dN/dS* metric that we used throughout this study:

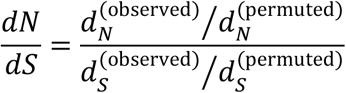

We confirmed that this approach accurately measures selection in the presence of simulated mutational biases (Fig. S2)

The number of permutations available for each gene/tri-nucleotide combination declines with gene length. Ultra-short genes may be too constrained for our permutation approach and underestimate selective pressures. While 12% of genes in our study harbored fewer than 10 permutations per mutation, these genes contained only ~ 3% of all mutations, as these genes are exceptionally short. Exclusion of these genes did not appreciably alter observed *dN/dS* patterns (Fig. S3E).

Mutations can be permuted across every identical tri-nucleotide context within a particular gene or every identical tri-nucleotide context within a particular transcript. For differentially-spliced genes, transcript and gene annotations differ: transcripts are comprised of a subset of exons that define the whole gene. Hence, WES data directly sequences transcripts, which can be overlaid along the genome to infer genes. Because transcript annotations directly match WES data, which comprises 85% of available samples, we chose to constrain permutations at the transcript level (ENST) rather than the gene level (ENSG or Hugo Symbols)^66^. This choice does not appreciably affect dN/dS patterns (Fig. S25), however there is a slight universal shift towards a dN/dS rate of 1 (in both drivers and passengers) when permuting at the gene level. Presumably, this is because exons exclusive to rare splicing variants experience weaker selective pressures (and/or less transcription-coupled DNA repair.) The subtle differences between gene-level and transcript-level null models may explain the subtle difference in genome-wide dN/dS levels between our approach and the dNdScv model^1^ (Fig. S3C).

Lastly, we note that binning nonsynonymous and synonymous mutations at the genomewide level (e.g. drivers and passengers) provided the most robust estimates of *dN/dS* when bootstrapping observed tumor samples. Statistical power is insufficient when binning at the individual gene level. Bootstrapping also demonstrated that log transformation of *dN/dS* values increases statistical power, and thus was generally applied to *dN/dS* analyses in this study.

#### A Parametric Null Model of Mutagenesis

For comparison, we also calculated *dN/dS* using dNdScv^67^ – a previously-published parametric null model of mutagenesis in cancer^1^. To compare both methods, dNdScv was ran globally and separately on samples stratified by the total number of substitutions using the following parameters:

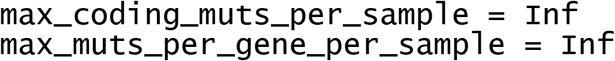

Global *dN/dS* values of all nonsynonymous mutations (*w*_all_, reported by dNdScv) were used. This model reproduced our nonparametric *dN/dS* trends (Fig. S3) and was used to infer patterns of selection in synonymous mutations (Fig. S19). We note that stratifying tumors in TCGA into 20 bins of equal sample-size (as was done in ^1^), rather than evenly-spaced bins, averages-out a significant proportion of the negative selection observed in passengers, since low mutation burden tumors reside within the tail-end of the distribution (Fig. S10).

#### Orthogonality of dN/dS with Mutational Burden and effects of excluding samples with no synonymous mutations

Mutational burden is generally calculated as the total number of substitutions within a sample (i.e. *d*_N_ + *d*_S_), however these tallies are also used in our measurement of *dN/dS.* Hence, any interdependence of mutational burden with *dN/dS* could bias our understanding of the relationship between selection and genome-wide mutation rate. We consider the interdependence of these two measures by assuming that both *d*_N_ and *d*_S_ are Poisson-distributed with rate parameters *λ*_N_ and *λ*_S_ The joint probability mass density of any combination of these two quantities is then:

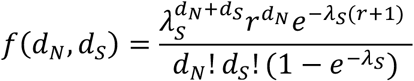

Here, *r* = *λ*_N_/*λ*_S_. The expectation value of *dN/dS,* for any degree of selection versus any combination of nonsynonymous and synonymous mutation tallies can then be calculated simply by exhaustively summing over all combinations that arise with probability above machine precision. In Figure S5, we compare the variation in *dN/dS* for a typical genome under neutral selection or equally-balanced positive and negative selection (r = 2.8) using the *d*_N_ + *d*_S_ and *d*_S_ mutational burden metrics. We observe less deviation from expectation using *d*_N_ + *d*_S_ primarily because *d*_S_ alone is a poor proxy for the mutation rate — i.e. there are far fewer synonymous mutations to use to estimate the mutation rate. *d*_N_ + *d*_S_ did exhibit slightly greater bias in observed *dN/dS* relative to expectation, however this bias was small compared to the variation in estimates (<5% for mutational burdens greater than 2) and biased observed estimates towards increased values of *dN/dS,* which will only understate the degree of negative selection. Lastly, we note that because the genome-wide *dN/dS* is approximately 1, deviations from these theoretical calculations should be minimal.

We also tested the effects of this non-orthogonality of our approach in three additional ways. First, we investigated the correlation of mutational burden metrics mutation rate in our simulated tumors (see below) and found that *d*_N_ + *d*_S_ correlated most strongly with mutation rate (Fig. S5C). Next, we randomly-partitioned all protein-coding mutations into two necessarily-orthogonal halves: a half that defined the mutational burden and a half that was used for calculating *dN/dS*. This partitioning found that selection patterns persisted (Fig. S5B). Finally using WGS data, we compared *dN/dS* to measures of mutational burden that excluded data from protein-coding regions (all intergenic and all intronic mutations), which once again represents a completely-orthogonal comparison of *dN/dS* with mutational burden (Fig. S3).

#### Identification of driver genes in cancer

For all analysis using SNVs, unless explicitly stated, a comprehensive list of 299 pan-cancer driver genes derived from 26 computational tools was used to catalog driver genes^16^. Other pan-cancer driver gene sets tested were derived from COSMIC’s Driver Gene Census^15^ (downloaded on October 2016) and IntOGen’s Cancer Drivers Database^17^ (v2014.12) which contained 602 and 459 number of driver genes, respectively.

Many driver genes are associated with only particular tumor subtypes. To compare patterns of selection across cancer subtypes without increasing or decreasing the size of the list for each subtype, we chose to use a single set of driver genes for most analyses. This may understate the degree of positive selection in driver genes as mutations in these genes may be passengers in some tumor subtypes. In Fig. S8, we investigate patterns of selection using the top 100 driver genes identified for each tumor type and observe decreased signatures of positive selection overall in driver genes. Nevertheless, the patterns of attenuated selection in drivers and passengers remains. While tissue-type specific driver genes certainly exist, our results suggest that our statistical power to detect drivers still remains too limited to justify subdividing analyses by tumor type in many cases.

For all CNA analysis, GISTIC 2.0^68^ was used to identify a set of genomic regions enriched for copy number gains and copy number losses using recommended settings with a confidence threshold of 0.9. CNAs used to identify these peaks were downloaded from the NIH Genomic Data Commons (GDC)^27^ in the TCGA cohort. For each amplification peak, the closest gene was annotated as a putative Oncogene, and similarly the closest gene to each deletion peak was annotated as a putative Tumor Suppressor. The top 100 amplification peaks (oncogenes) and deletion peaks (Tumor Suppressors) were classified as drivers for each of the 32 tumor types. 34% of identified driver genes appear in more than one tumor type, while 2.6% of identified driver genes appear in more than five tumor types.

For both SNV and CNA analysis, passengers were defined as mutations that did not reside within driver genes. The vast majority of mutations are passengers, and their relative totals for both SNVs and CNAs are depicted in Fig. S24.

#### Annotation of clonal and subclonal mutations

Since TCGA contains SNVs with high coverage and available purity estimates, only MC3 SNVs (exclusive to TCGA) were used in this analysis (WGS read-depth is generally lower than WES read-depth). Variant allele frequencies (VAFs) were calculated per site as the number of mutant read counts divided by the total number of read counts. VAFs were adjusted for purity using calls made by ABSOLUTE^27,69^, collected from GDC. A VAF threshold of 0.2 was used to define ‘subclonal’ (< 0.2) vs ‘clonal’ (> 0.2) SNVs. Different VAF thresholds were considered (Fig. S12) and the choice of ‘clonal’ thresholding did not impact the conclusions of this study.

#### Polyphen2 analysis

PolyPhen2 annotations in the MC3 SNP calls were used^18^. Only missense mutations that were categorized as either ‘benign’, ‘probably damaging’ or ‘possibly damaging’ were used. The fraction of pathogenic missense mutations was calculated as the number of pathogenic mutations categorized as either “probably damaging” or “possibly damaging” divided by the total number of categorized mutations.

#### Classification of genes by functional category

To test for patterns of selection in functionally related genes, we annotated all mutations by different functional categories and Gene Ontology (GO) terms^26^. Oncogenes and tumor suppressors were annotated from a curated set of 99 high confidence cancer genes^70^. Essential genes were collected from a genome-wide CRISPR screen that identified genes required for proliferation and survival in a human cancer cell line^71^. Housekeeping genes were defined as genes with an exon that is expressed in all tissues at any nonzero level, and exhibits a uniform expression level across tissues^72^. Interacting proteins were downloaded from the mentha database in April 2019^73^.

To identify highly expressed genes, median transcripts per million (TPM) in 54 tissue types (v7 release) were downloaded from the Genotype-Tissue Expression (GTEx) project^45^. Tissues that contained high expression in most genes, specifically testes, were removed. Only genes that had TPM counts above zero in any of the 53 remaining tissues were used. TPM counts were averaged across all tissues. Highly expressed genes were defined as the top 1000 genes expressed across all tissues.

To test for signals of negative selection in other functional groups, we annotated mutations by candidate GO terms according to Biological Processes: Transcription Regulation (GO Term ID: 0140110), Translation Regulation (GO Term ID: 0045182), and Chromosome Segregation (GO Term ID: 0007059).

#### Somatic Copy Number Alteration (CNAs)

All CNAs were downloaded from the COSMIC database on June 2015^15^. Mitochondrial CNAs were discarded from analysis, as copy number changes are difficult to infer. Gene annotations and the locations of telomeres and centromeres were downloaded from the UCSC Genome Browser (hg19). Telomeric and centromeric regions were masked from all measurements of *dE/dI.* Because the selection patterns of non-focal CNAs — alterations with at least one terminus in a telomere or centromeric region — were not noticeably different from long (>100kb) focal CNAs, these two alteration classes were aggregated for analysis. Notably, we observed positive selection for both amplifications and deletions within oncogenes, and for both deletions and amplifications within Tumor Suppressors. For this reason, we did not distinguish between gains and losses, nor oncogenes and Tumor Suppressors in published analyses: any CNA that overlapped an oncogene or tumor suppressor in any region (for any fraction of the CNA) was classified as a driver. Mutational burden was defined simply as the total number of CNAs within a sample. Pan-cancer CNAs from cBioPortal (August 2018) were also analyzed, however consistent purity and ploidy estimates could not be obtained by using either ABSOLUTE^69^ or TITAN^74^, so this data was not used for published analyses of CNAs.

#### Measurements of selection on CNAs

*dE/dI* was calculated using a ‘Breakpoint Frequency’ metric and a ‘Fractional Overlap’ metric. For both metrics, the *dE/dI* of a particular gene set *i* (e.g. driver or passenger genes) is defined by a genomic track *T_j.g_*, which is one for every annotated region *g* of the track and zero elsewhere. Only non-centromeric and non-telomeric regions are considered in the mappable human genome *G.* Each CNA *C*_g,m_ is defined by its position on the genome *g* and the mutational burden *m* of the tumor harboring the mutation. For ‘Breakpoint Frequency’ *C*_m,i_ is one at the position of both termini of the CNA and zero elsewhere. For ‘Fractional Overlap’ *C*_m,i_ is *1/L,* where *L* is the length of the CNA, for every region of the genome spanned by the CNA and zero elsewhere. For a particular range of mutational burdens *M*, *dE/dI* was defined as:

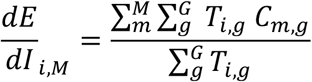

We note that calculation is accelerated by >100x by commuting *T*_i,g_ with the outer summation 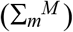. Lastly, we randomly permuted the start and stop positions of each CNA, while preserving its length, to derive a set of neutral CNAs not experiencing selection. This permutation analysis finds that *dE/dI* for both breakpoint frequency and fractional overlap is ~1 in the absence of selection (Fig. S23).

#### Tumor purity analysis in TCGA samples

Tumor purity estimates from the ABSOLUTE algorithm^69^ were downloaded from the GDC on May 2020. For all tumors and for tumors with < = 10 substitutions, correlation coefficients between the total number of substitutions and tumor purity were calculated. To evaluate the effects of tumor purity on patterns of selection, tumors below increasing thresholds of tumor purity were removed from the analysis, and dN/dS was calculated on tumors stratified by mutational burden bins (as described above.)

#### Expression analysis

Gene expression data was downloaded from the COSMIC database on September 2019. Genes used to identify different protein folding pathways were downloaded from ^75^, genes involved in protein degradation pathways were identified from ^76^. The median gene expression of all genes in each protein folding pathway was used. Patients were binned by the total number of substitutions (using MC3 SNP calls from TCGA) and CNAs, and the average gene expression of each bin was calculated.

#### Cancer subtype analysis

All tumor subtypes in TCGA and ICGC were grouped into 9 subcategories, based on broad, predominantly anatomical features. Anatomical features (i.e. organ and systems of organs), rather than histological features or inferred cell-of-origin, were used as groupings because we believe that the fitness effects of mutations should be predominantly defined by the environment of the tumor. Nevertheless, we observed attenuated selection in both drivers and passengers in many broad histologically defined classifications (e.g. adenocarcinomas & sarcomas). For all cancer grouping analysis (broad and subtype), tumors were stratified into bins by the total number of substitutions (*d*_N_ + *d*_S_) on a log scale. Since tumor subtypes vary in their range of mutational burdens, (e.g. KIRC cancer subtypes only have tumors with <100 substitutions), *dN/dS* values in the lowest and highest mutational burden bin for each cancer-subtype are shown.

Specific cancer subtype categories were taken directly from the NCI Genomic Data Commons (GDC)^27^. Because CNAs were downloaded from COSMIC, CNA datasets were not classified with this same ontology. Table S1 details how CNA classifications were mapped on GDC categories (and sometimes more broadly-defined groups). All subtypes with >200 samples were used in our CNA subtype analyses (Fig. S15).

#### An evolutionary model with Hill-Robertson Interference

Somatic cells in our populations are modeled as individual cells that can stochastically divide and die in a first-order (memoryless) Gillespie Algorithm. This model was developed and described previously^33^. During division, cells can acquire advantageous drivers with rate *μT*_drivers_ and deleterious passengers with rate *μT*_passengers_ - these values specify the mean of Poisson-distributed pseudo-random number (PRN) generators that prescribe the number of drivers and passengers conferred during division (e.g. the number of drivers per division *n_d_* = Poisson[*n*_d_ = *k*; λ = *μT*_drivers_] = *λ^k^e^−k^/k!*). The Distribution of Fitness Effects (DFE) conferred by each driver and each passenger are Exponentially-distributed PRNs with probability densities *P(s_i_* = *x; s*_drivers_) = Exp[-*x/S*_drivers_]/*s*_drivers_ and *P(S_i_* = *x*; *s*_passengers_) = *-* Exp[*-x/ S*_passengers_]/*s*_passengers_ respectively. Simulations with other exponential-family DFEs do not qualitatively differ from these exponential distributions^28^. The aggregate absolute cellular fitness is 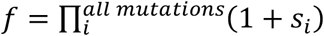 in our Multiplicative Epistasis model and Δ*f* = *S_i_*/(1 + *vf*) with *v* = 1 in our Diminishing-Returns Epistasis Model where *Δf* is the change in cellular fitness with each mutation^77^. The rate of cell birth is inversely proportional to cellular fitness, while the rate of cell death 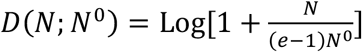 increases with the population size of the tumor *N*. With these birth and death processes, mean population size abides by a Gompertzian growth law in the absence of additional mutations, which is scaled by the mean cellular fitness E[*N*(< *f* >)] = Log[1 + < *f* > */ N*^0^] (derived from Master Equation^28^). While, programmatically, mutations exclusively affect the birth rate and the constraints on growth exclusively affect the death rate, we previously demonstrated that birth and death rates are generally nearly-balanced such that dynamics are not affected by this design choice.

Because somatic cells do not recombine during cell division, dominance coefficients were not explicitly modeled. Thus in diploid cancers, our selection coefficients estimate the mean heterozygous effect of drivers and passenger (i.e. *hs)* Similarly, Loss of Heterozygosity (LOH) events (gene losses, gene conversions, mitotic recombination, etc) are not explicitly modeled either; however, these events can be viewed as additional mutations that may be either adaptive drivers or deleterious passengers in the model. As sequencing data improves, we believe that it will be informative to explicitly model dominance coefficients, tumor ploidy, and LOH events.

Simulations progressed until tumor extinction (*N* = 0 cells), malignant transformation (*N* = 10^6^ cells), or until approximately 100 years had passed (18,500 generations). Only fixed mutations (present in the Most Recent Common Ancestor) within clinically-detectable growths were analyzed in our ABC pipeline. The behavior of this model has been described previously^28,33^ and the most relevant assumptions of this model and their effects on the conclusions of this study are described in Table S2.

Cells in our populations are fully described by their accrued mutations, and birth and death times. Birth and death events were modeled using an implementation of the Next Reaction^78^, a Gillespie Algorithm that orders events using a Heap Queue. Generation time in our model was defined as the inverse of the mean birth rate of the population: 1/ <B(d, p)>. While all mutation events occurred during cell division, if mutations were to occur per unit of time (rather than per generation), rapidly growing tumors would acquire drivers at a slightly slower rate as generation times decline over time. This effect, however, is negligible compared to the variation in waiting times conferred by the variation in mutation rates (division times merely double, while mutation rates vary by 100,000-fold).

This simple evolutionary model is defined by five parameters *μT*_drivers_, *μT*_passengers_, *s*_drivers_, *s*_passengers_, and *N*^0^. The target size of drivers is defined as the approximate number of nonsynonymous mutations in the Bailey Driver Screen *T*_drivers_ = (# of driver genes)•(mean driver length)•(fraction of SNVs that are nonsynonymous) = 300 genes • 1298 loci/gene • 0.737 nonsynonymous loci / loci = 286,886 nonsynonymous loci. The target size of passengers was simply the remaining loci in the protein coding genome, *T*_passengers_ = 20,451,136 nonsynonymous loci. The mutation rate was constant throughout each tumor simulation and randomly-sampled from a uniform distribution in log-space that ranged from 10^-12^ to 10^-7^ mutations•loci^-1^•generation^-1^. While tumors were initiated from this broad range, malignancies (N > 10^6^ cells) were almost always restricted to mutation rates between 10^-10^ and 10^-8^ (Fig. S20), as tumors with mutation rates drawn below this range almost never progressed to cancer within 100 years and tumors with mutation rates drawn above this range went extinct through natural selection.

The likelihood that tumors progress to cancer in the presence of deleterious passengers depends heavily on the initial population size *N^0^* of the tumor. This dependence was studied previously^33^, where it was demonstrated that reasonable evolutionary simulations (those that progress to cancer >10% of the time, but less than 90% of the time) are restricted to a fourdimensional manifold *N\** within the five-dimensional phase space of parameters. For this reason, *N*^0^ N^*^(*s*_drivers_, *s*_passengers_, *μT*_drivers_, *μT*_passengers_) was determined by the other four parameters. To first-order, this manifold is *T*_passengers_ *s*_paassengers_ *T* (*T*_drivers_ *s*_drivers_^2^), however a more precise estimate (Eq. S8 of ^33^) incorporating more precise estimates of Muller’s Ratchet and the effects of hitchhiking on both driver and passenger accumulation rates, which does not exist in closed form was used. Additionally, at very low values of *s*_drivers_, progression to cancer is limited by time, not by the accumulation of deleterious passengers. Hence, we assigned *N^0^* such that:

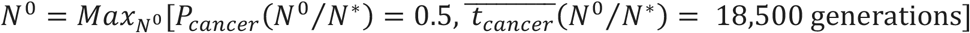

Here, *P*_cancer_ and *t*_cancer_ – the likelihood and waiting-time to cancer – are defined by equations S8 and S12 respectively in ^33^. *N^0^* was determined from these equations using Brent’s Method. Supplementary Figure 17 depicts the values of *N^0^,* which ranged from 1 to 100 for all simulations.

In tumors that progress to malignancy (*N* = 10^6^), only fixed nonsynonymous mutations (present in all simulated cells) were recorded. We also recorded (i) the fitness effect of these mutations, (ii) the mean population fitness, (iii) the number of generations until malignancy, and (iv) the mutation rate. These two values were used to generate the number of synonymous drivers and passengers, where P(*d*_s_ = *k*) = Poisson[*k*; *λ* = *μT*drivers/passengers */r t*_MRCA_] defines the number of synonymous drivers/passengers conferred, *t*_MRCA_ represents the number of division until the Most Recent Common Ancestor arose in the simulation, *r* = 2.795 represents the ratio of nonsynonymous to synonymous loci within the genome, weighted by the genome-wide trinucleotide somatic mutation rate, and the Poisson PRN generator was defined above. In simulations where synonymous drivers could arise, a fraction of the recorded nonsynonymous mutations (ranging from 0 – 20%) were simply re-labeled as synonymous drivers (as opposed to nonsynonymous drivers). This was done, again, by Poisson-sampling in proportion to the desired fraction for each cancer simulation.

20 × 20 combinations of *s*_drivers_ and *s*_passengers_ parameters were simulated (Fig. S16 & S17). Simulations were repeated until 10,000 cancers at each parameter combination were obtained or until 10 million tumor populations were simulated. While we attempted to initiate tumors at a population size where the probability of progression to cancer was 50%, some parameter combinations still did not yield 10,000 cancers after 10 million attempts (i.e. *P*_cancer_ < 0.1%). These combinations were predominately at low values of *s*_drivers_, which were far from the MLE estimate of *s*_drivers_ and represent unrealistic evolutionary scenarios: drivers cannot be weakly beneficial, relegated to only 300 genes, and still overcome deleterious passengers within 100 years. These simulations are annotated as “Progression Impossible.” Simulation parameter sweeps were performed for both the Multiplicative and Diminishing Returns Epistasis models. Twenty fractions of synonymous drivers were also generated (ranging from 0% to 20%). These fractions were generated by simply re-labeling the driver mutations which conferred fitness (generated during the simulation) as synonymous, instead of nonsynonymous.

#### Summary statistics of simulated and observed tumors

For both simulated and observed data, we summarized *dN/dS* rates versus mutational burden for drivers and for passengers by decade-sized bins: (0, 10], (10, 100], (100, 1,000]. Mutational burden for simulations was defined as the total number of substitutions (*d*_N_ + *d*_S_) – exactly as it was defined for observed data. For simulated data, *dN/dS* = *d*_N_/(*d*_S_ • r). Like observed data, *dN/dS* rates attenuated towards 1 for both drivers and passengers for all values of *s*_drivers_ and *S*_passengers_.

Mutational Burdens (MB) for simulated and observed data were summarized with the parameters of a Negative Binomial distribution, where 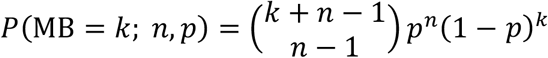. This distribution has been used previously to summarize the mutational burdens of human tumors ^79^ and exactly defines the expected number of mutations at transformation in a MultiStage Model of Tumorigenesis^30^ when *n* drivers are needed for transformation and the probability that any mutation be a driver is 1 -*p* ^80^. Both *n* and*p* were used to summarize MB. These quantities were determined by Maximum Likelihood optimization of the probability mass function above over the support of mutational burdens of [1, 1,000] substitutions. The Han-Powell quasi-Newton Least-squares method was used for optimization.

Age-dependent Cancer Incidence rates (CI) were summarized with the parameters of a Gamma distribution, where 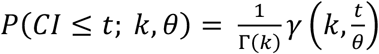. Here, 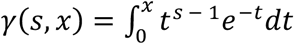 is the lower incomplete gamma function and Γ(*k*) = γ(*k*, ∞) is the regular gamma function. Similar to our summarization of mutational burdens, this distribution is a generalization of the exact waiting time to transformation expected from a Multi-Stage Model of Tumorigenesis when tumors arise at a uniform rate over time, require *k* drivers for transformation, and wait an average time of *θ* between drivers ^80^. This Cumulative Distribution Function was fit to observed incidence rates for all patients above 20 years of age using the least squares numerical optimization defined above (All cancer sites combined, both sexes, all races, 2012 - 2016 ^81^). Patients under 20 years of age were excluded because cancers in these patients generally arise from germline predispositions to cancer, which are (i) not directly modeled by our simulations, (ii) not detected as somatic mutations, and (iii) result in age-incidence curves that do not agree with a Gamma distribution^30^. Because all cancer simulations are initiated at *t = 0* (instead of uniformly in time, as is presumed in the Multi-Stage Model), the simulated data was fit using the probability density function of this distribution (instantaneous derivative) using Maximum Likelihood and the optimization algorithm described above. The cumulative distribution, then, represents the expected age-incidence cancer incidence rate when simulations begin at uniformly-distributed moments in time and, thus, was used to generate Figure 3D. Only the shape parameter *k* was used in ABC (and *θ* was ignored), as this parameter only specifies the dimensionality of time (simulation time was measured in cellular generations, not years) and all values of *θ* in our simulations are equivalent under a Gauge transformation. Additionally, we do not expect the exact times of incidence to be particularly informative as the time of transformation is generally somewhat earlier than the time of detection.

#### Use of Approximate Bayesian Criterion (ABC) for model selection and parameter inference

Like many Bayesian analyses, the main steps of an ABC analysis scheme are: (1) formulate a model, (2) fit the model to data (parameter estimation), and (3) improve the model by checking its fit (posterior-predictive checks) and (4) comparing this model to other models 82,83

The nine summary statistics described above were used to compare simulations to observed data. Agreement was summarized with a Log-Euclidian distance, as all summary statistics resided on the domain [0, ∞) and log-transformation of the summary statistics minimized heteroscedasticity of the simulated data relative to a square-root or no transformation. Variance of the summary statistics was not normalized. ABC was performed using the ‘abc’ R package^82^.

The rejection method (Feedforward Neural Net) and tolerance (0.5) were chosen based on their capacity to minimize prediction error of the simulated data using Leave-one-out Cross Validation (CV, Fig. S18A). 10,000 instances of the neural network, which was restricted to a single layer, were initiated and the median prediction of these networks were used. These parameters were used for both model comparison and parameter inference. The posterior model probability (postpr) was used to compare the two epistatic models (Diminishing Returns versus Multiplicative). The likelihood of the data under the Diminishing Returns model (14%) was less than the likelihood under the Multiplicative Epistasis Model (86%). For parameter inferencing, the *s*_drivers_ and *s*_passengers_ prior values were log-transformed.

For the synonymous driver model, the base model (without synonymous drivers) was simply the lowest quantity of synonymous drivers (0%) in the parameter sweep of synonymous driver quantities (Fig. S18B). The posterior probability mass of this value 0.043 was used as the one-sided *p*-value for the null hypothesis that these two models are equally predictive. Although the synonymous driver model agreed with the observed data slightly-better, *s*_drivers_ and *S*_passengers_ parameters could not be inferred from the data because the potential for synonymous drivers destroys the utility of a *dN/dS* statistics, which is predicated on the notion that synonymous mutations are neutral. Virtually any value of *dN/dS* is attainable when the right combinations of selective pressures on nonsynonymous and synonymous are paired (Fig. S18C).

### Supplementary Note

Here, we consider alternative explanations for the attenuation of *dN/dS* rates in driver genes with increasing Tumor Mutational Burden (TMB). In the main text, we argue that the attenuation in both the driver and the passenger *dN/dS* curves observed within cancers are most consistent with evolutionary models where passengers are deleterious and, thus, interfere with natural selection. Certainly, the attenuation of *dN/dS* in passengers is the most direct evidence for the deleteriousness of these mutations; however, the attenuation of *dN/dS* of drivers due to Hill-Robertson Interference is also strong evidence of for the deleteriousness of passengers.

We cannot reasonably entertain evolutionary models where Hill-Robertson Interference does not exist because it is an inextricable consequence of asexual evolution. Instead in this note, we consider other factors that might also impart an attenuating *dN/dS* in drivers in an asexually-evolving tumor and discuss how these models are inconsistent with the observed genomic patterns within drivers.

#### Evolutionary models with a fixed quantity of driver mutations (multi-hit model)

Suppose progression to malignancy is contingent upon acquisition of a fixed number of nonsynonymous driver mutations (we refer to this as a ‘multi-hit’ model), while the acquisition of passenger mutations (including synonymous mutations within driver genes) is entirely random. In such a scenario, TMB will abide by a negative binomial distribution with the ratio of drivers to passengers necessarily increasing as TMB decreases - driver quantities are fixed, while passengers are randomly dispersed. Because synonymous driver mutations are also passengers, the quantity of these mutations will also decrease with TMB, thereby increasing the driver *dN/dS* ratio as TMB decreases.

While driver dN/dS will decrease with TMB in this model, this model also predicts that tumors will exhibit a constant quantity of nonsynonymous drivers and a paucity of synonymous driver mutations at low TMB. Quite the contrary, low TMB exhibit slightly *more* synonymous driver mutations than expected (Fig. S19A). Furthermore, the quantity of nonsynonymous drivers increases with TMB suggesting that higher TMB tumors must acquire more drivers to overcome their deleterious passenger load.

#### Evolutionary models where a fraction of nonsynonymous mutations within driver genes are neutral (causing selection bias for drivers at low TMB)

The multi-hit tumorigenesis model might still be capable of explaining observed *dN/dS* patterns in drivers, if we assume that a fraction of nonsynonymous mutations within driver genes are not drivers, but instead neutral mutations. This model can recapitulate the increase in nonsynonymous drivers with TMB because low TMB tumors are conditionally-required to harbor fewer neutral mutations overall (including fewer mutations in passenger genes, fewer synonymous mutations, and fewer neutral nonsynonymous mutation within driver genes).

Unfortunately, the quantitative shape of this model is still inconsistent with the observed *dN/dS* driver curve.

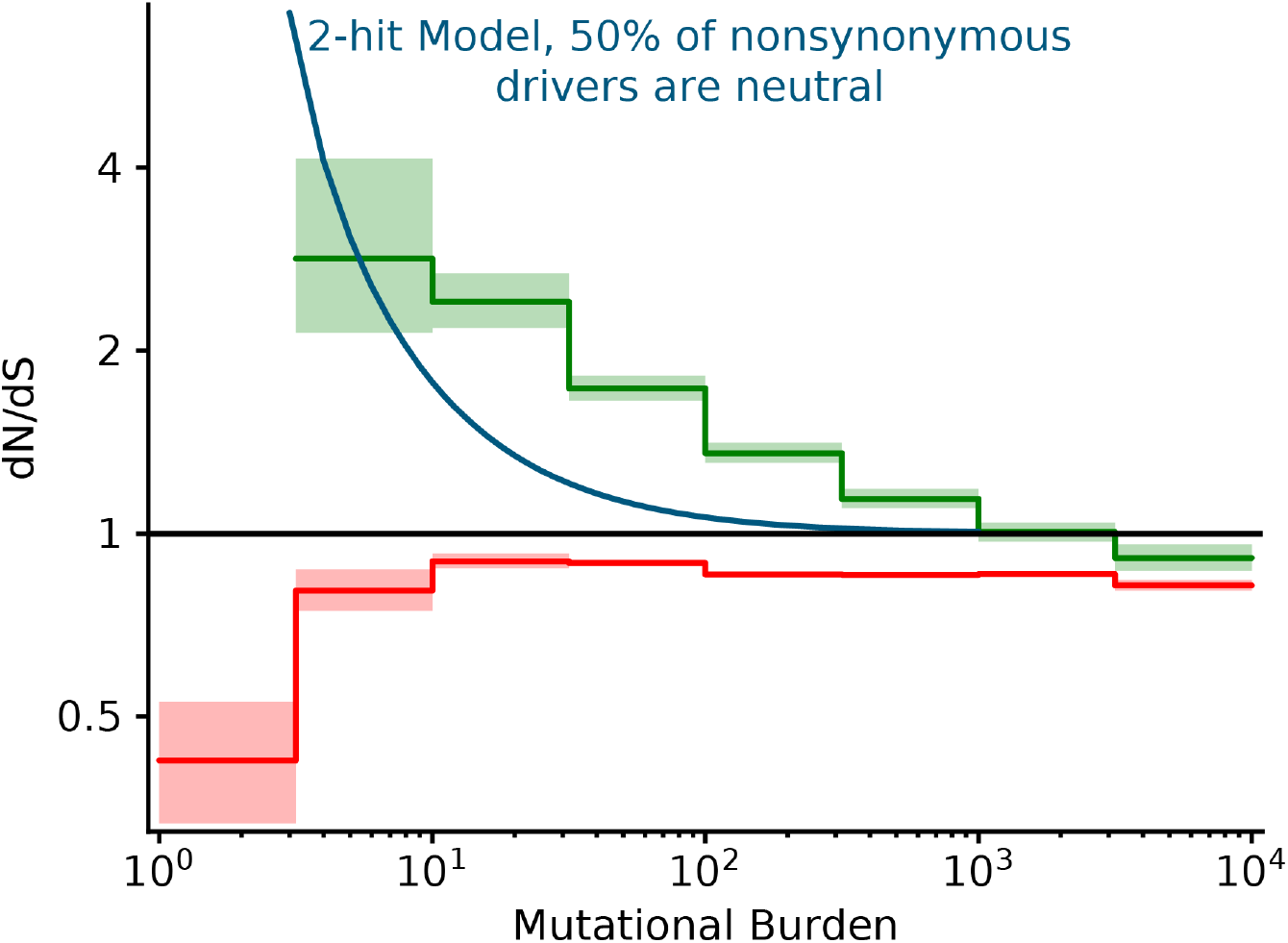
A multi-hit tumor model without deleterious passengers cannot explain dN/dS patterns in drivers. We considered a model of tumorigenesis where a fixed quantity of drivers *n* is required for tumorigenesis. Passengers, synonymous driver mutations, and neutral nonsynonymous mutations within driver genes (50% of nonsynonymous mutations within these genes) were all negative-binomially distributed random variates with shape parameter *n*. Expected dN/dS values for this model were calculated exactly. The resulting dN/dS curve in drivers (blue) declines hyperbolically with TMB and cannot explain observed dN/dS patterns in drivers (green).

Specifically, the effects of selection bias for nonsynonymous drivers decline faster than the observed attenuation in drivers. This model predicts a hyperbolic decline, whereas we found that the observed dN/dS in drivers is better-described by a logistic curve after log-transforming TMB.

#### Evolutionary models with diminishing returns to driver mutations

Additionally, we considered an extension of our model of tumor evolution with adaptive drivers and deleterious passengers where driver mutations impart a weaker fitness advantage as the background fitness of a cell increases (diminishing-returns epistasis). Specifically, the change in fitness *Δf* imparted by a driver *i* with untransformed fitness benefit *st* is *Δf = s_i_*/(1 + *f).* This model might also explain the decline in *dN/dS* in driver genes without invoking the need for a substantial deleterious passenger load. Using our ABC procedure and the same prior probability distributions (10^-2^ – 10^0^ for mean driver fitness benefit and 10^-4^ – 10^-2^ for mean passenger fitness cost) we find that this model cannot explain observed dN/dS patterns in drivers and passengers better than our base model with multiplicative epistasis between all mutations. The likelihood of the data under the Diminishing Returns model (14%) was less than the likelihood under the Multiplicative Epistasis Model (86%).

### Supplementary Figures

**Supplemental Figure 1.**
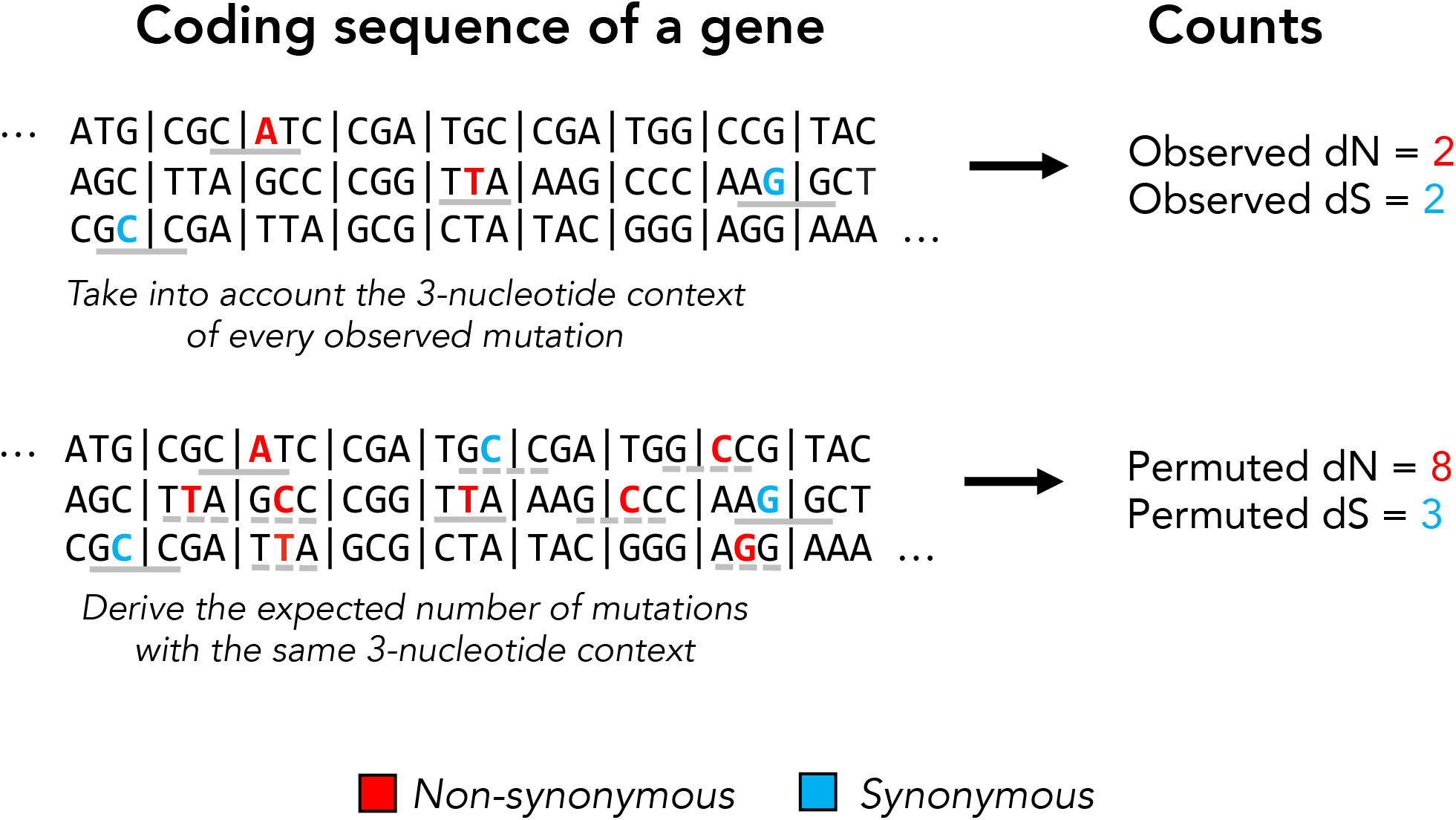
Schematic of our permuted dN and dS calculation. Permuted synonymous and nonsynonymous counts are used to account for mutational biases in dN/dS calculations. Observed mutations and their 3-nucleotide context is shown in a solid gray bar. Permuted mutations with the same 3-nucleotide context are shown in dashed gray lines. Note that permutations do not preserve the codon position of a mutation and can alter protein coding effect (nonsynonymous vs. synonymous).

**Supplemental Figure 2.**
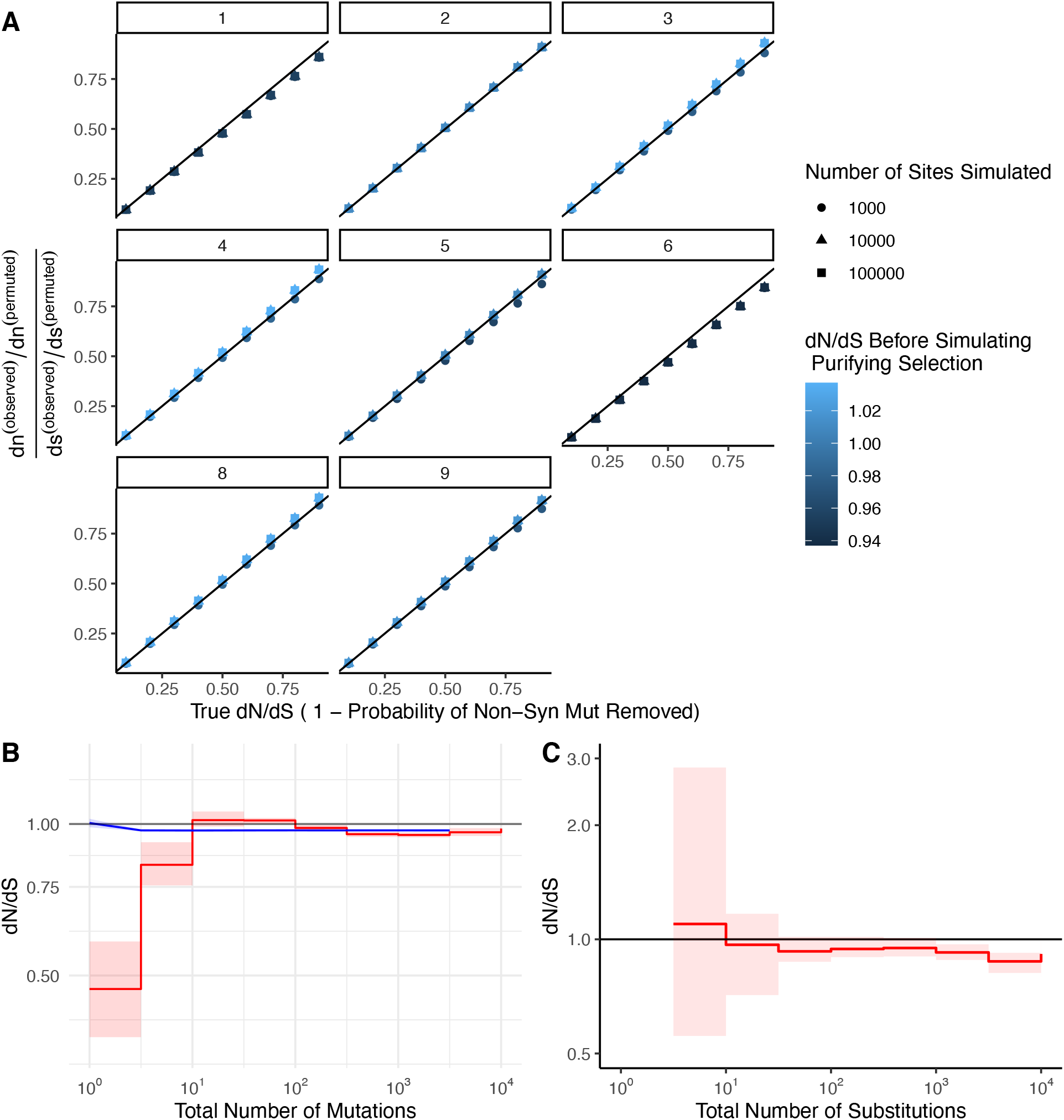
Permutation-based null model of mutagenesis corrects for mutational biases in dN/dS calculations. **A.** Simulations (*N* = 100) of negative selection under extreme mutational bias scenarios where all mutations are generated from a single Mutational Signature (e.g. APOBEC or smoking, COSMIC Signatures 1-9, grey titles). Bias-corrected dN/dS values calculated from these simulations are compared to simulated levels of negative selection. Colors denote bias-corrected dN/dS before negative selection was simulated, which is expected to be neutral (~1). Negative selection is simulated as the probability of randomly removing nonsynonymous mutations, (e.g. a simulated ‘true’ dN/dS of 0.1 defines simulations where each nonsynonymous mutation had a 90% probability of removal). Shapes correspond to different numbers of sites simulated. Black line identifies perfect correspondence between bias-correct dN/dS and simulated (true) dN/dS. **B.** 95% confidence intervals of dN/dS in passenger mutations randomly sampled in blue (N=1000) from high mutational burden tumors (> 10 substitutions) in the same proportion of sites as binned in Fig. 2A. Red line denotes observed dN/dS of passengers in ICGC and TCGA as depicted in Fig. 2A. **C.** dN/dS of weakly expressed genes (defined as having < 1 TPM across all samples in GTEx) in tumors stratified by the total number of substitutions within ICGC and TCGA. Solid black shows dN/dS values of 1, expected under neutrality. Error bars are 95% confidence intervals determined by bootstrap sampling.

**Supplemental Figure 3.**
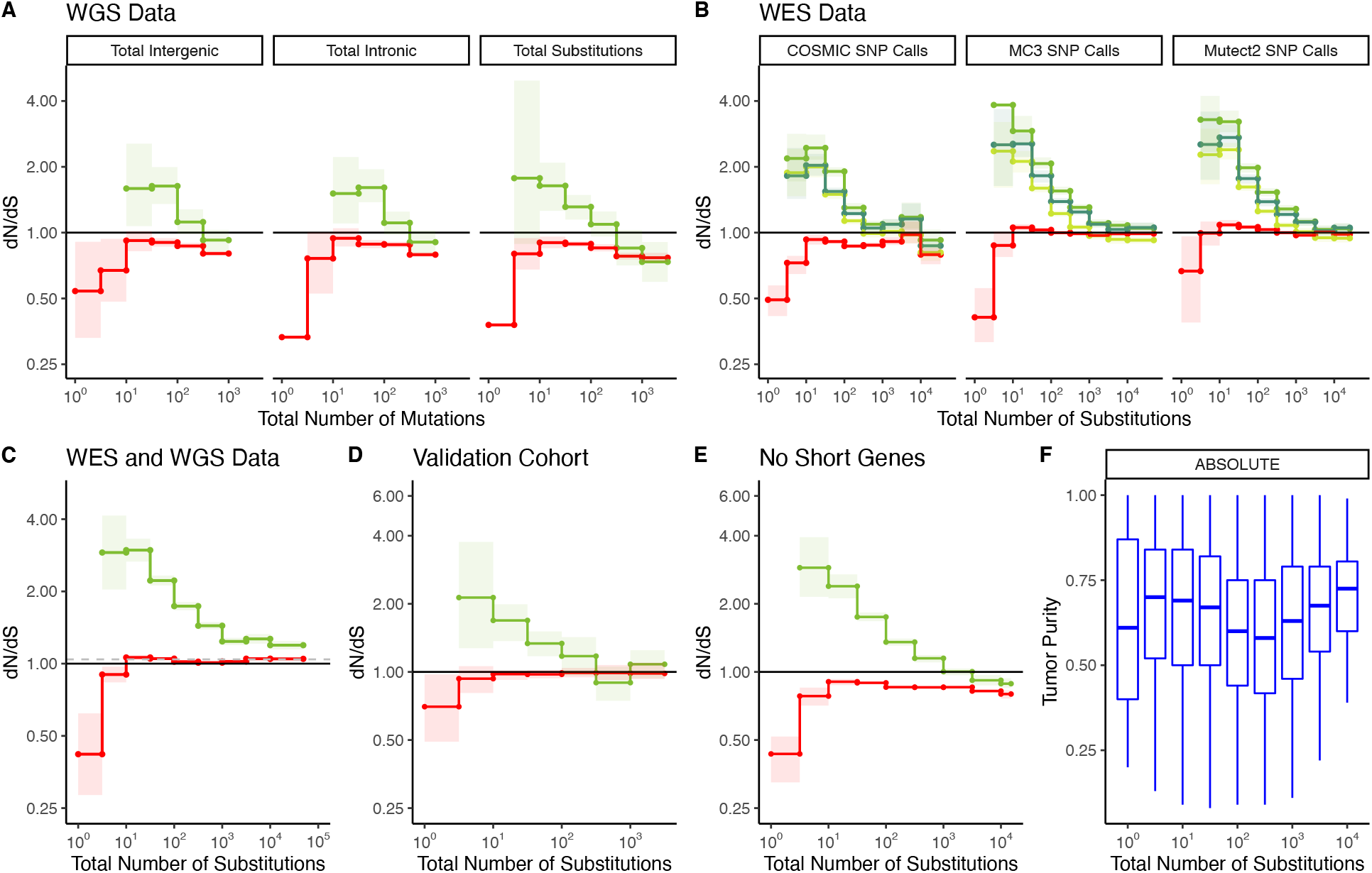
Patterns of attenuated selection persist across mutation burden metrics, sequencing platforms, mutation calling algorithms, data repositories, and choice of driver gene set. **(A-C)** dN/dS calculations within passenger and driver gene sets for various burden metrics, sequencing platforms, mutation calling algorithm, choice of driver gene set, and data repository. The solid black line (dN/dS = 1) annotates expected dN/dS under neutrality in all panels. Error bars are 95% confidence intervals determined by bootstrap sampling. **(A)** Tumors in ICGC stratified by either the total number of intergenic mutations, intronic mutations or substitutions. **(B)** dN/dS calculations for various pan-cancer driver gene sets stratified by the total number of substitutions. Shown are tumors within TCGA called by different mutation callers (Mutect2 vs consensus, MC3 SNP calls), and SNV calls from COSMIC. **(C)** dN/dS calculations within passenger and driver gene sets within tumors in ICGC and TCGA stratified by the total number of substitutions. Instead of using our nonparametric null model, we calculate dN/dS using dNdScv^1^ as a null model of mutagenesis (with default parameters and unrestricted quantities of coding mutations per gene). Grey dashed line represents global dN/dS values of all tumors without stratifying by mutational burden. **(D)** Validation of dN/dS calculations within passenger and driver gene sets in primary untreated tumors, distinct from ICGC and TCGA, stratified by the total number of substitutions**. (E)** dN/dS of tumors in TCGA and ICGC stratified by the total number of substitutions after removing short genes, i.e. genes with fewer than 10 permutations. **(F)**. ABSOLUTE purity estimates in TGCA from the GDC in tumors stratified by the total number of substitutions.

**Supplemental Figure 4.**
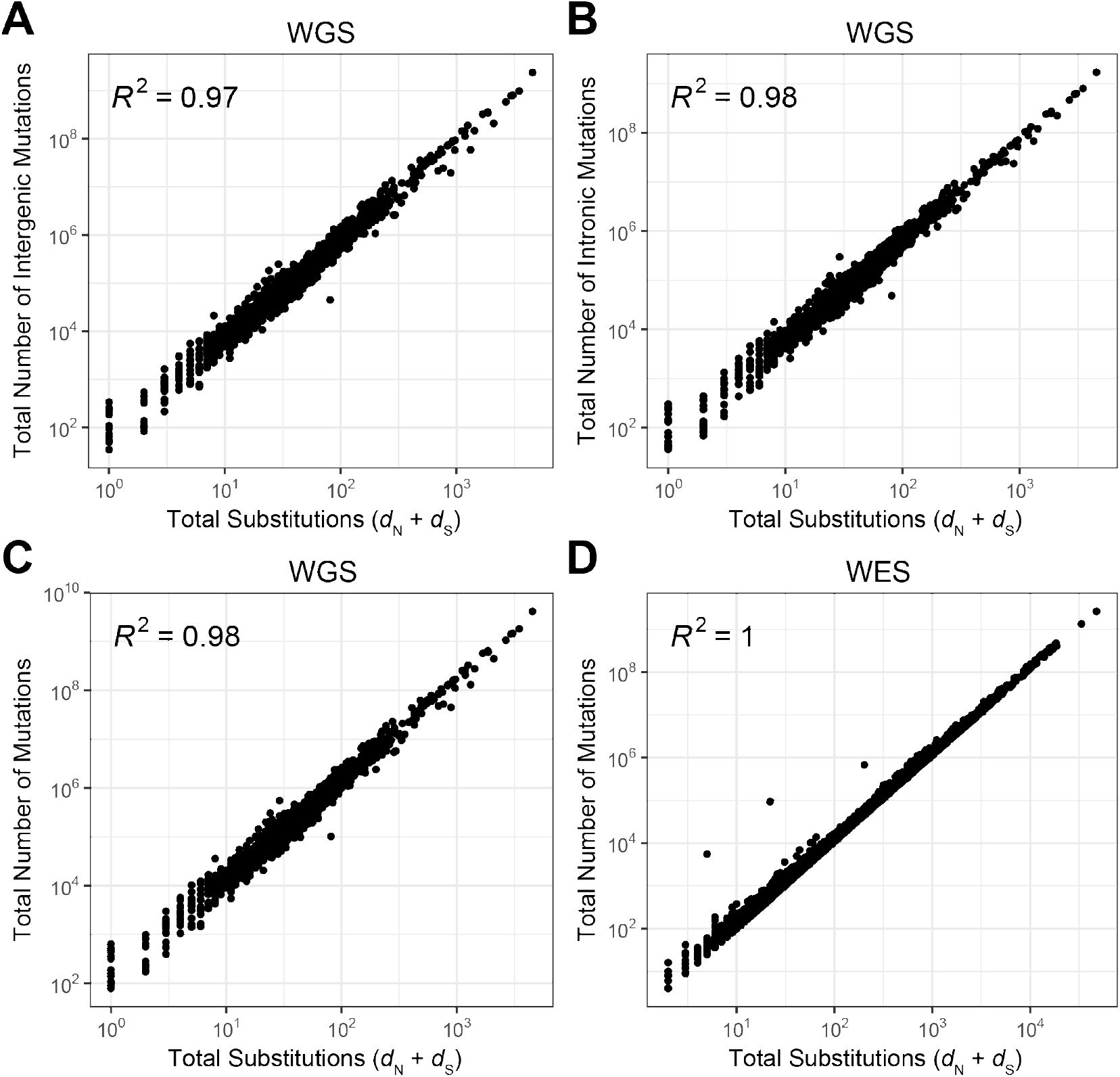
Mutation burden metrics, used as a proxy for the tumor mutation rate, are correlated across datasets. **(A)** Correlation between the total number of substitutions and the total number of intergenic or **(B)** intronic mutations within tumors in TCGA (WES). **(C)** Correlation between the total number of mutations (TMB) and total number of substitutions for tumors in ICGC (WGS) and **(D)** and TCGA (WES). Because all mutational burden metric are highly correlated, general patterns of selection are unaffected by choice of mutational burden metric.

**Supplemental Figure 5.**
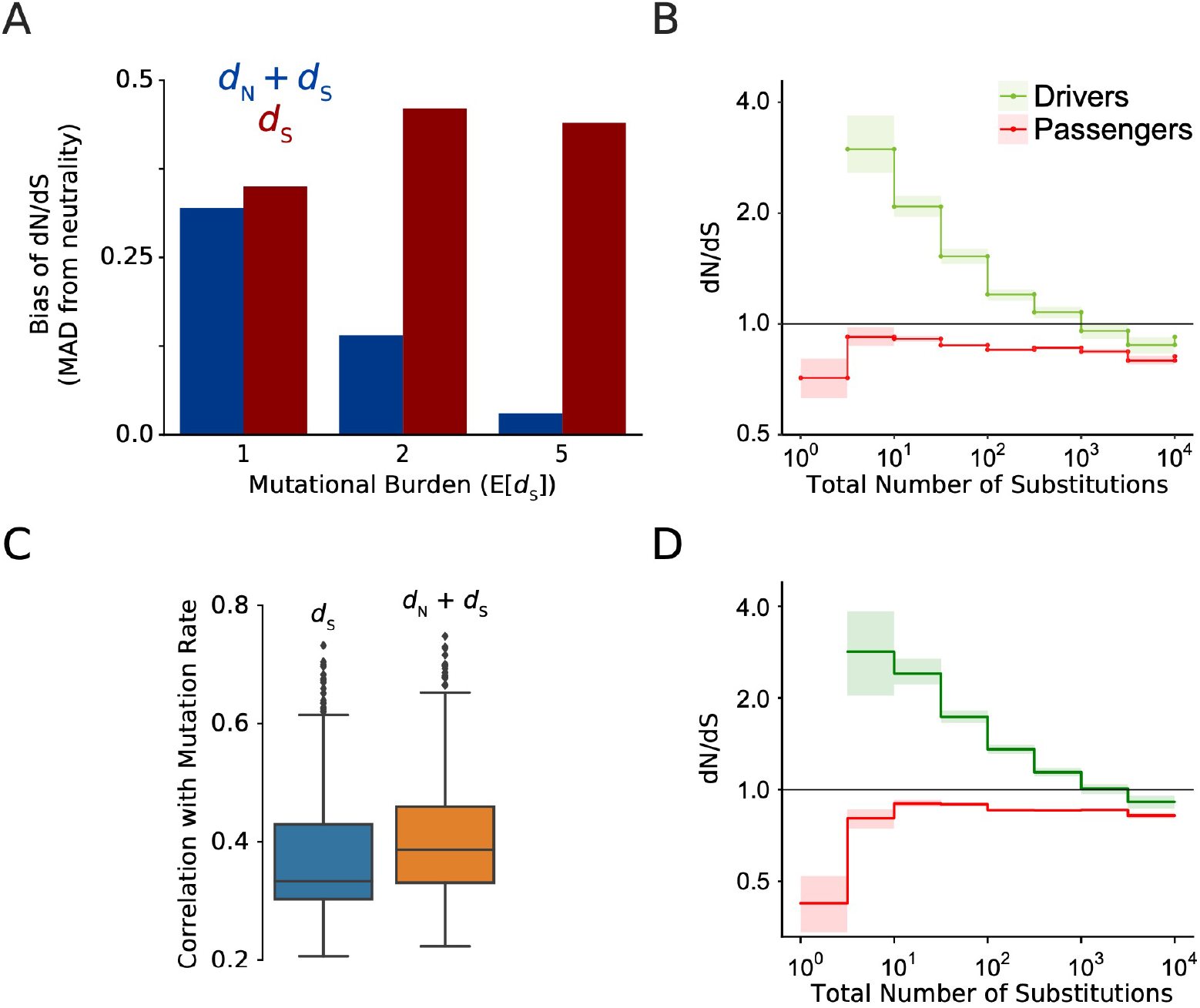
Stratification of dN/dS by mutational burden (defined as *d_N_* + *d*_S_) does not bias dN/dS values and correlates well with mutation rate in simulations. **(A)** Theoretical bias of dN/dS (Mean Absolute Deviation from neutrality) of mutational burden metrics that contribute to dN/dS calculations. *d*_N_ + *d*_S_ (i.e. Total Substitutions) imparts less bias than *d*_S_ (i.e. Total Synonymous Substitutions). Bias determined by analytical model of dN/dS with ratios of Poisson-sampled mutation tallies (Methods). Bias rapidly decreases with mutational burden for *d*_N_ + *d*_S_. Total Substitutions (*d*_N_ + *d*_S_) exhibit less bias than Total Synonymous Substitutions (*d*_S_). **(B)** Patterns of selection persist when independent mutation counts (completely orthogonal) were used for estimating selection (dN/dS) and mutational burden (*d*_N_ + *d*_S_). Independent accounts were achieved by randomly partitioning mutations into two halves and using one half to calculate dN/dS and the half to calculate Total Number of Substitutions separately. Tumors were from TCGA. dN/dS and Error Bars (95% Confidence Interval) are same as in Figure 2. Solid black line of 1 denotes dN/dS expected under neutrality. **(C)** Pearson correlation of both mutational burden measures with mutation rate in computational model of tumor evolution (Methods). The mutational burdens of ~4 million simulated cancers were compared to their programmed mutation rate. *d*_N_ + *d*_S_ correlated well with mutation rates across a range of evolutionary parameters and was more highly correlated with mutation rate than *d*_S_ alone. **(D)** Same as in Figure 2A of the main text, except tumors with no synonymous mutations and tumors with no nonsynonymous mutations are included. dN/dS values at low Mutational Burdens are not appreciably altered by these filters.

**Supplementary Figure 6.**
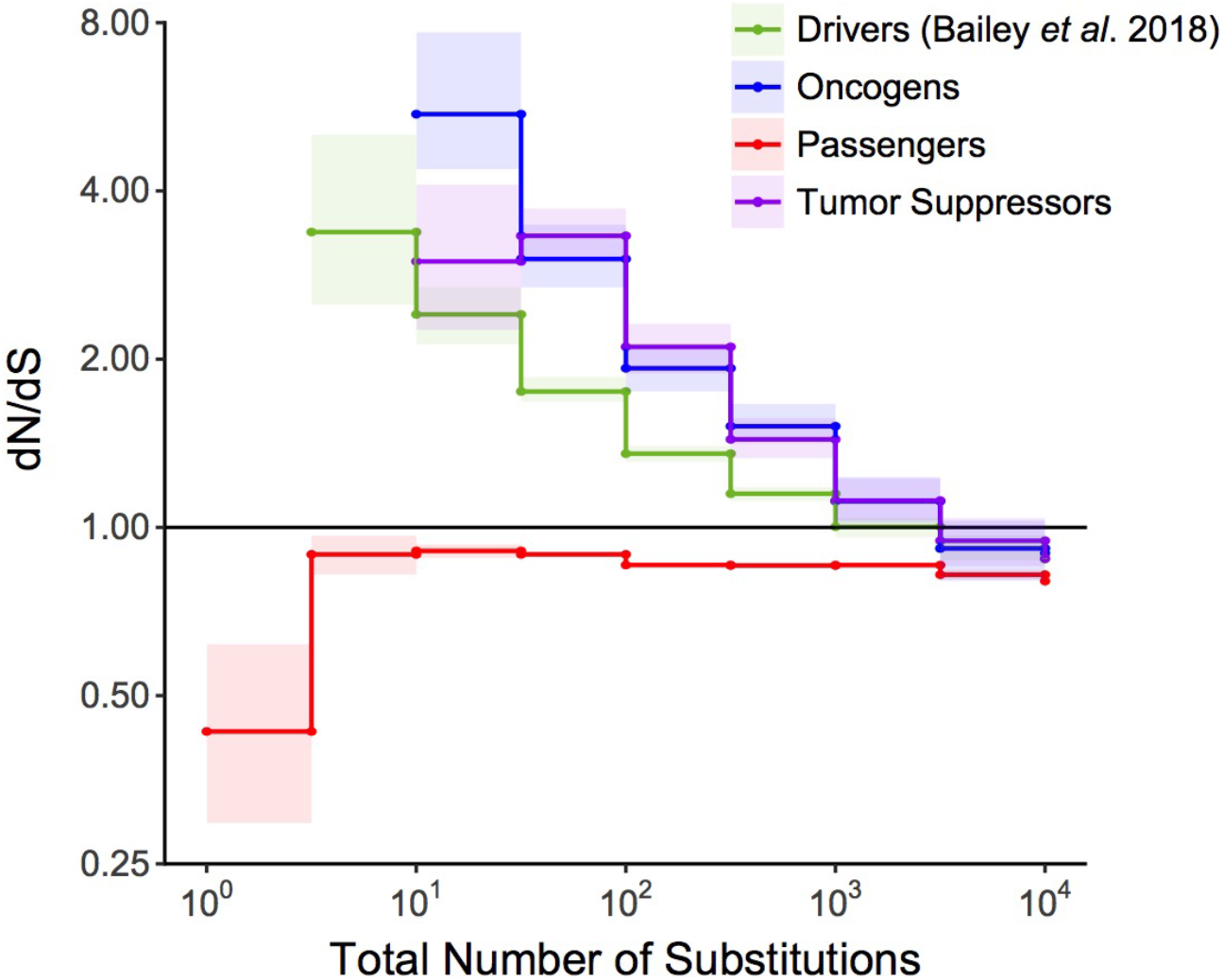
Attenuation of selection with increasing mutational burden in both Oncogenes and Tumor Suppressors. dN/dS of passenger and driver gene sets^16^ within tumors in TCGA stratified by the total number of substitutions present in the tumor (*d*_N_ + *d*_S_). Tumor suppressors (purple), oncogenes (blue) and pan-cancer driver (green) gene sets are shown. Solid black shows dN/dS values of 1, expected under neutrality. Error bars are 95% confidence intervals determined by bootstrap sampling.

**Supplemental Figure 7.**
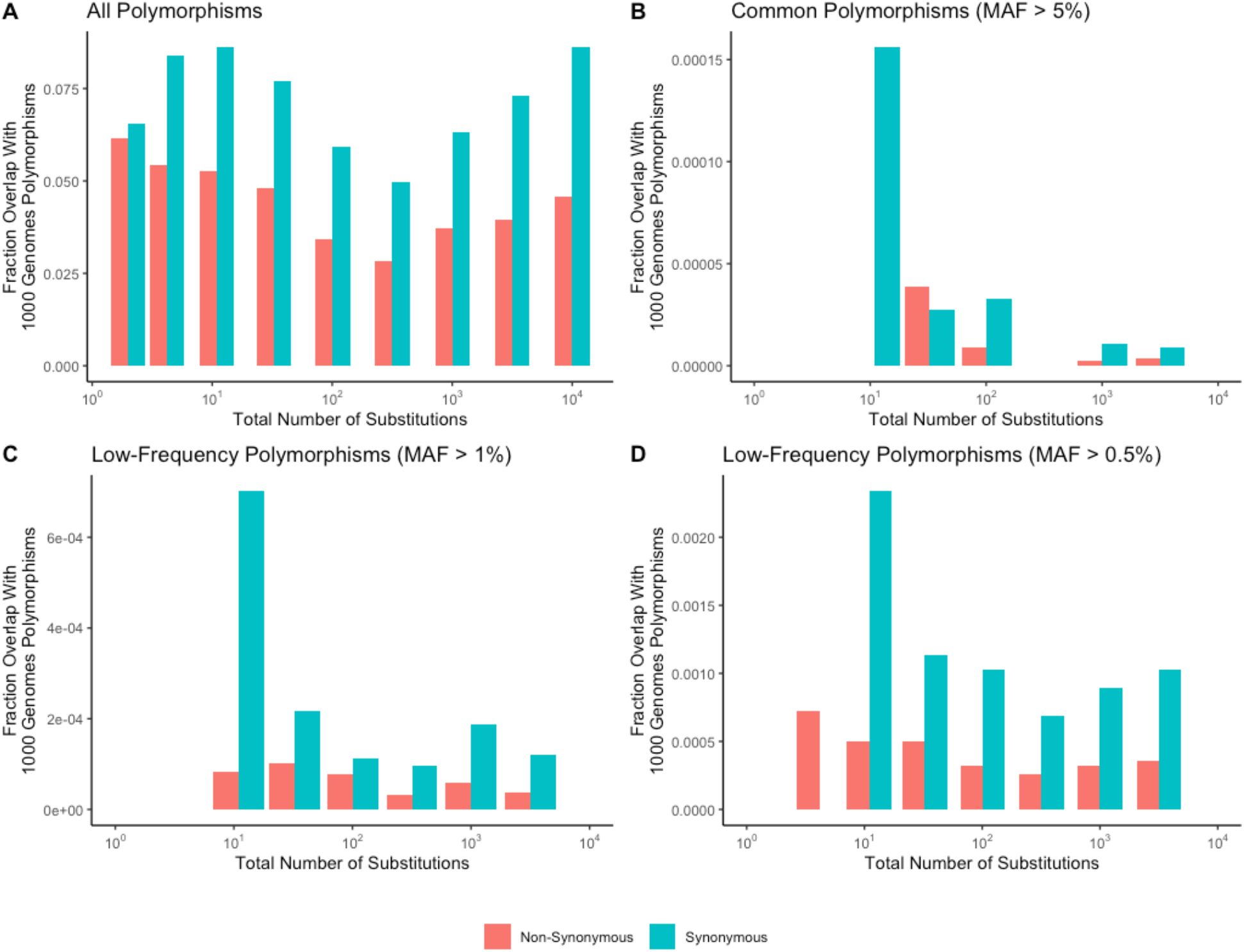
No common germline polymorphisms observed in low mutation rate cancers. **(A)** Fraction of mutations that overlap all germline polymorphisms in the 1000 Genomes Project within tumors stratified by the total number of substitutions. **(B-D)** Fraction of mutations that overlap only common (MAF > 0.05, 0.01 or 0.005) polymorphisms in the 1000 Genomes Project within tumors stratified by the total number of substitutions. WGS and WES datasets are shown. Colors denote mutations that are synonymous (blue) or nonsynonymous (red). Strong negative germline selection is expected only within common polymorphisms. No mutations within low mutational burden cancers (≤10 substitutions) overlap common polymorphic sites (when MAF > 0.1). Note that there are no synonymous mutations at MAF > 0.05 within low mutational burden cancers that could lower dN/dS rates through germline contamination.

**Supplemental Figure 8.**
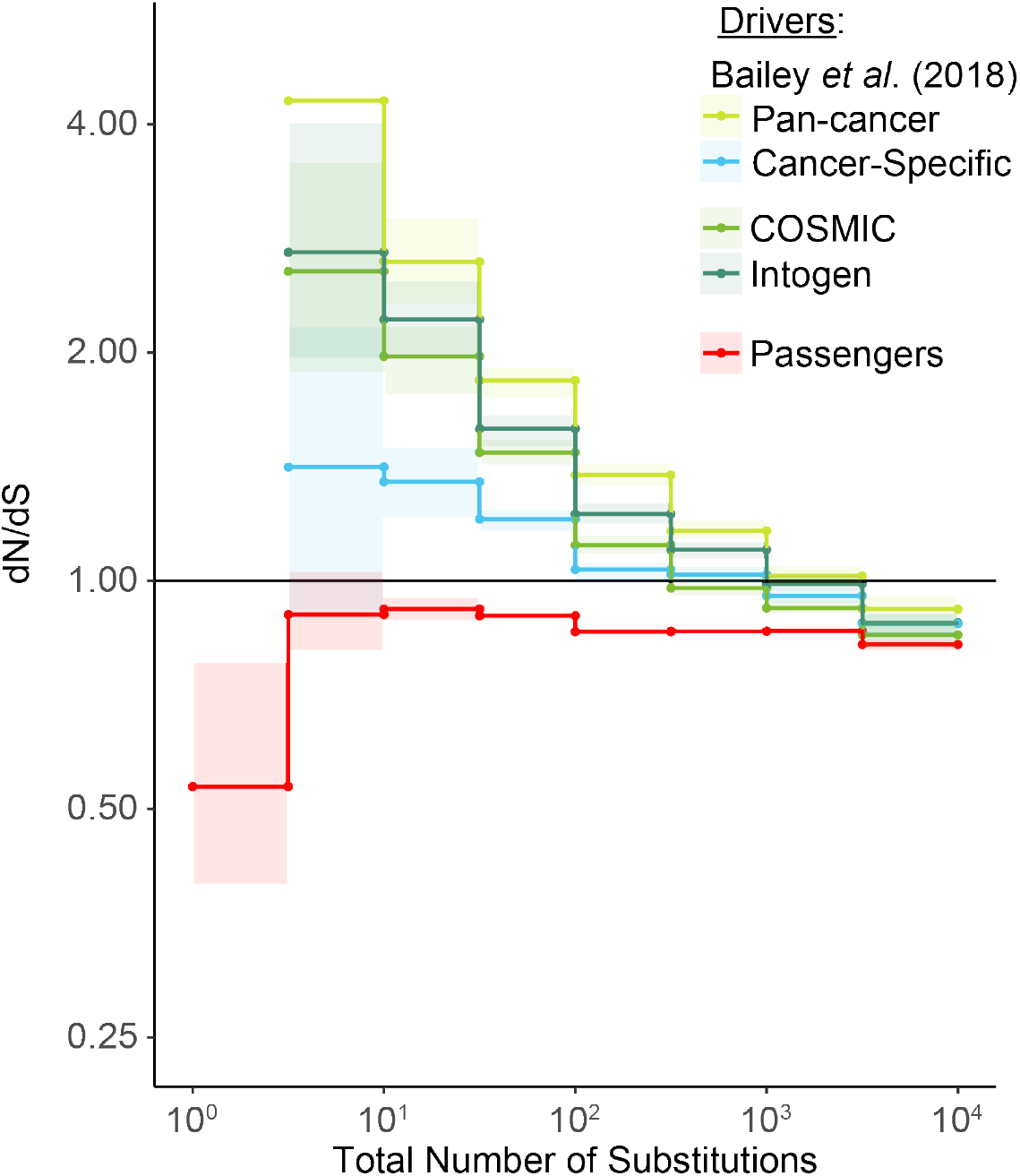
Weaker signals of positive selection within cancerspecific drivers. dN/dS values of passenger and different driver gene sets within tumors in TCGA stratified by the total number of substitutions present in the tumor (*d*_N_ + *d*_S_). Pan-cancer driver (lime) and cancer-specific (blue) driver gene sets identified by Bailey *et al.* 2018^16^ are shown. Pan-cancer driver genes identified in this study also exhibited stronger signatures of positive selection than driver genes identified by COSMIC^84^ (light green) and Intogen^17^ (forest green). Hence, pan-cancer drivers from Bailey *et al.* 2018 were used throughout this study. Cancer-specific gene sets are defined as the top 100 recurrently mutated genes within the particular cancer type, and used separately for each of the 33 cancer types in TCGA. Solid black shows dN/dS values of 1, expected under neutrality. Error bars are 95% confidence intervals determined by bootstrap sampling.

**Supplemental Figure 9.**
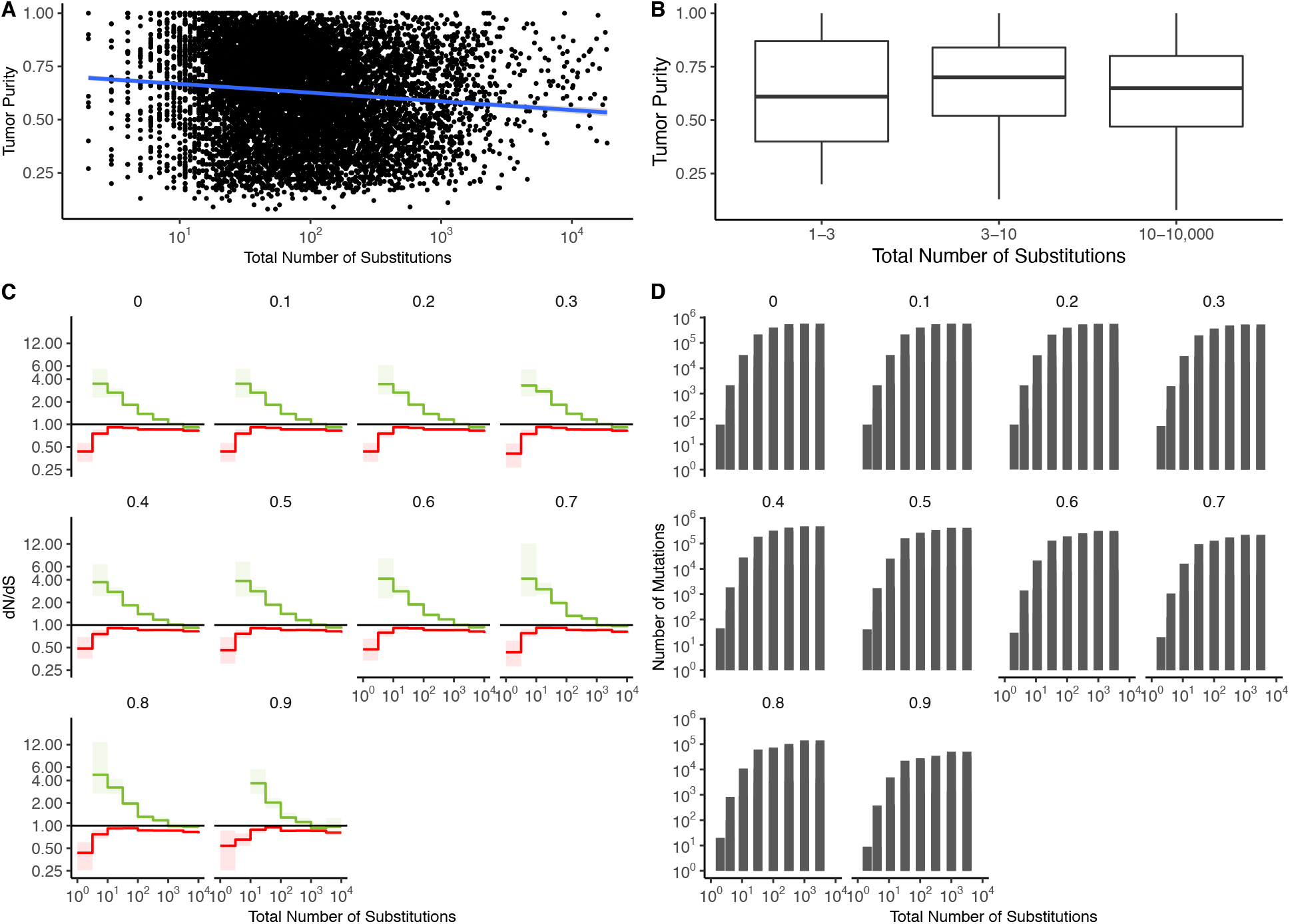
Patterns of attenuated selection persist across tumor purity thresholds. **(A).** Correlation between tumor purity (calculated by GDC using the ABSOLUTE^69^ algorithm, Methods) and the total number of substitutions in all TCGA samples (*r* = −0.0008, *R^2^* = 7 x 10^-7^). Blue line denotes a linear regression fit and grey colors denote the 95% confidence intervals for the fit of this linear model. **(B).** Boxplot of tumor purity in TCGA samples stratified into low mutation rate bins (1-3 and 3-10 substitutions) and high mutation rate bins (10-10,000 substitutions).(**C).** dN/dS in driver (green) and passenger (red) gene sets of tumors in TCGA stratified by the total number of substitutions after removing tumors below various purity thresholds. Values at the top denote the threshold of tumors removed from the analysis. (e.g. 0.3 shows dN/dS of tumors with a purity >= 0.3.) **(D)** Number of mutations in each bin within **(C)** after removing tumors at increasing purity thresholds. Error bars are 95% confidence intervals determined by bootstrap sampling.

**Supplemental Figure 10.**
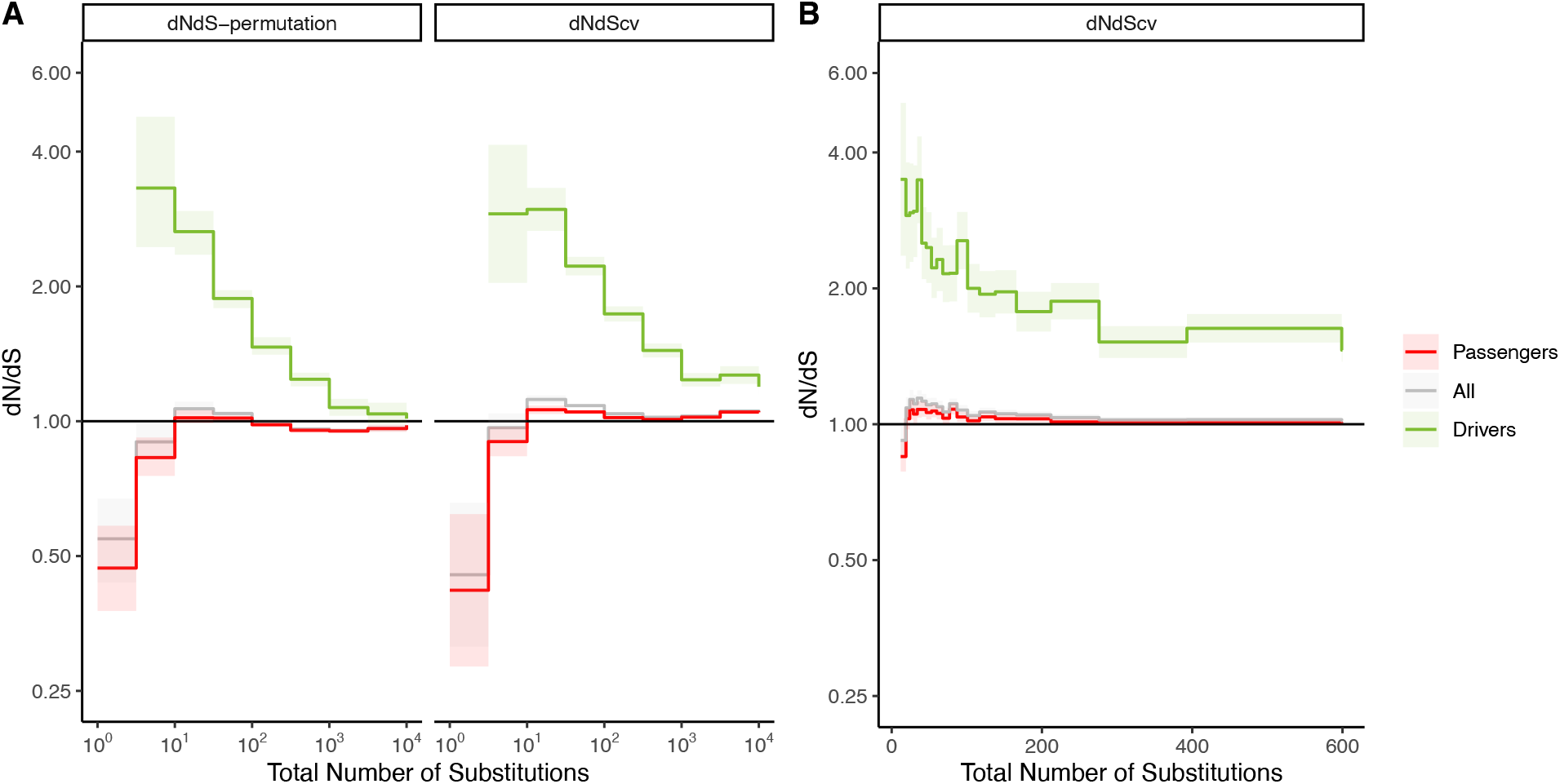
Comparison of dN/dS to results in Martincorena *et al.* (2017) for tumors stratified by mutational burden. **(A)**. dN/dS in driver (green), passenger (red) and all gene sets (grey) of tumors in TCGA stratified by the total number of substitutions using 9 bins of equal width (log-scale TMB), as depicted in Figure 2. Left panel uses our non-parametric null model of mutagenesis to calculate dN/dS, while the right panel uses dNdScv (from Martincorena *et al.* 2017) as a null model of mutagenesis. Error bars are 95% confidence intervals determined by bootstrap sampling. **(B).** dN/dS of driver (green), passenger (red) and all gene sets (grey) of tumors in TCGA stratified by the total number of substitutions using 20 bins of equal sample sizes, as was done in Figure 5 of Martincorena *et.al.* 2017. The binning scheme and linear axes compress results at low TMB. To replicate Martincorena *et.al.* 2017, three tumor types were also excluded in this analysis: UVM, CHOL, and DLBC. DNdScv was used as a null model of mutagenesis. dN/dS for driver and passenger genes sets was not calculated in Figure 5 of Martincorena *et al* (2017). Error bars are 95% confidence intervals derived from dNdScv.

**Supplemental Figure 11.**
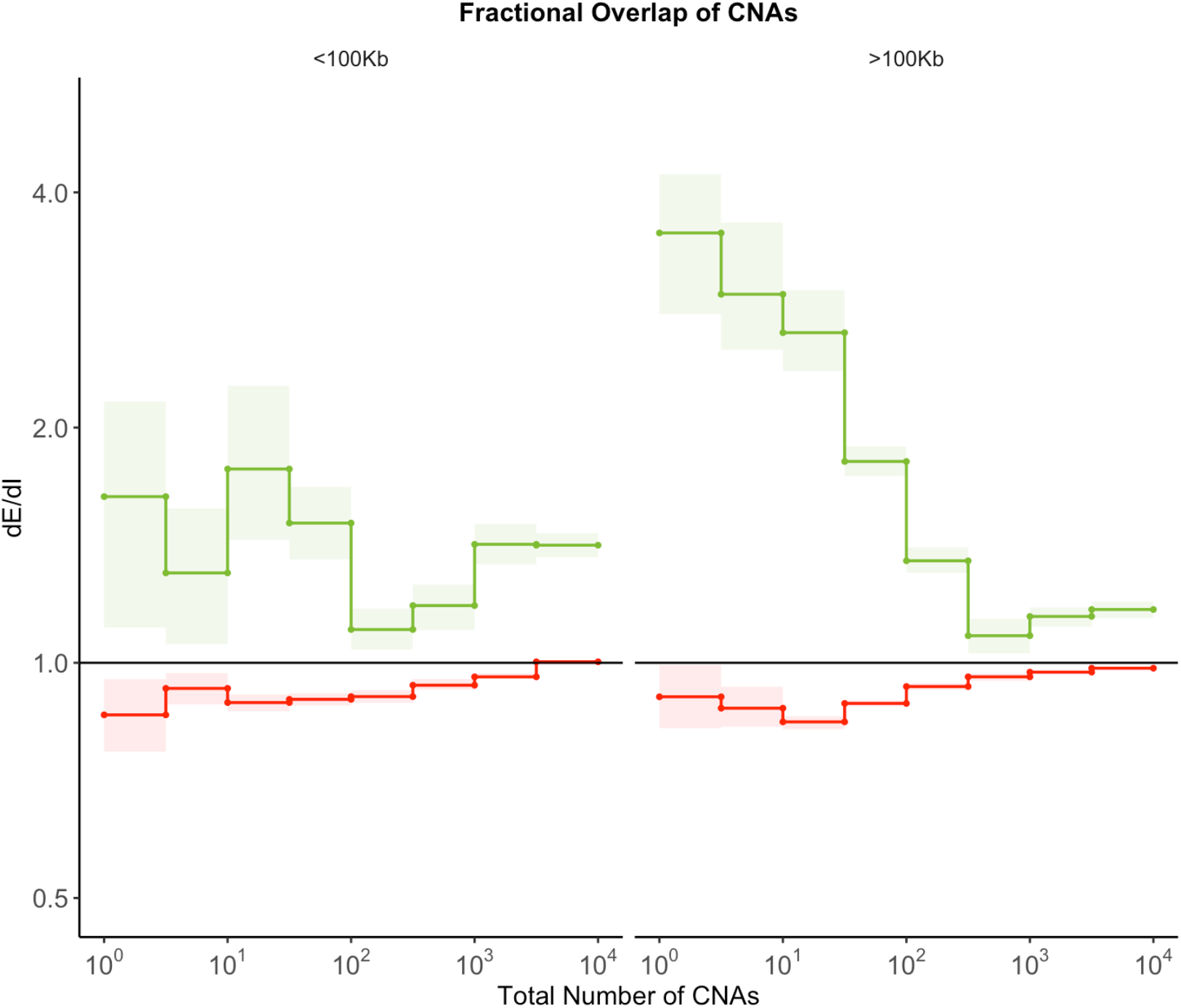
Fractional overlap of CNAs within exomic regions (dE) relative to intergenic regions (dI) exhibits similar patterns of selection as Fractional Overlap. Calculations of fractional overlap^20^ of exomic regions (dE) to intergenic (dI) regions within passenger and GISTIC^68^ driver gene sets in tumors stratified by the total number of CNAs present. dE/dI is shown separately for CNAs greater than 100Kb in length (right) and smaller than 100Kb in length (left). Solid black line of 1 denotes values expected under neutrality. Error bars are 95% confidence intervals determined by bootstrap sampling.

**Supplemental Figure 12.**
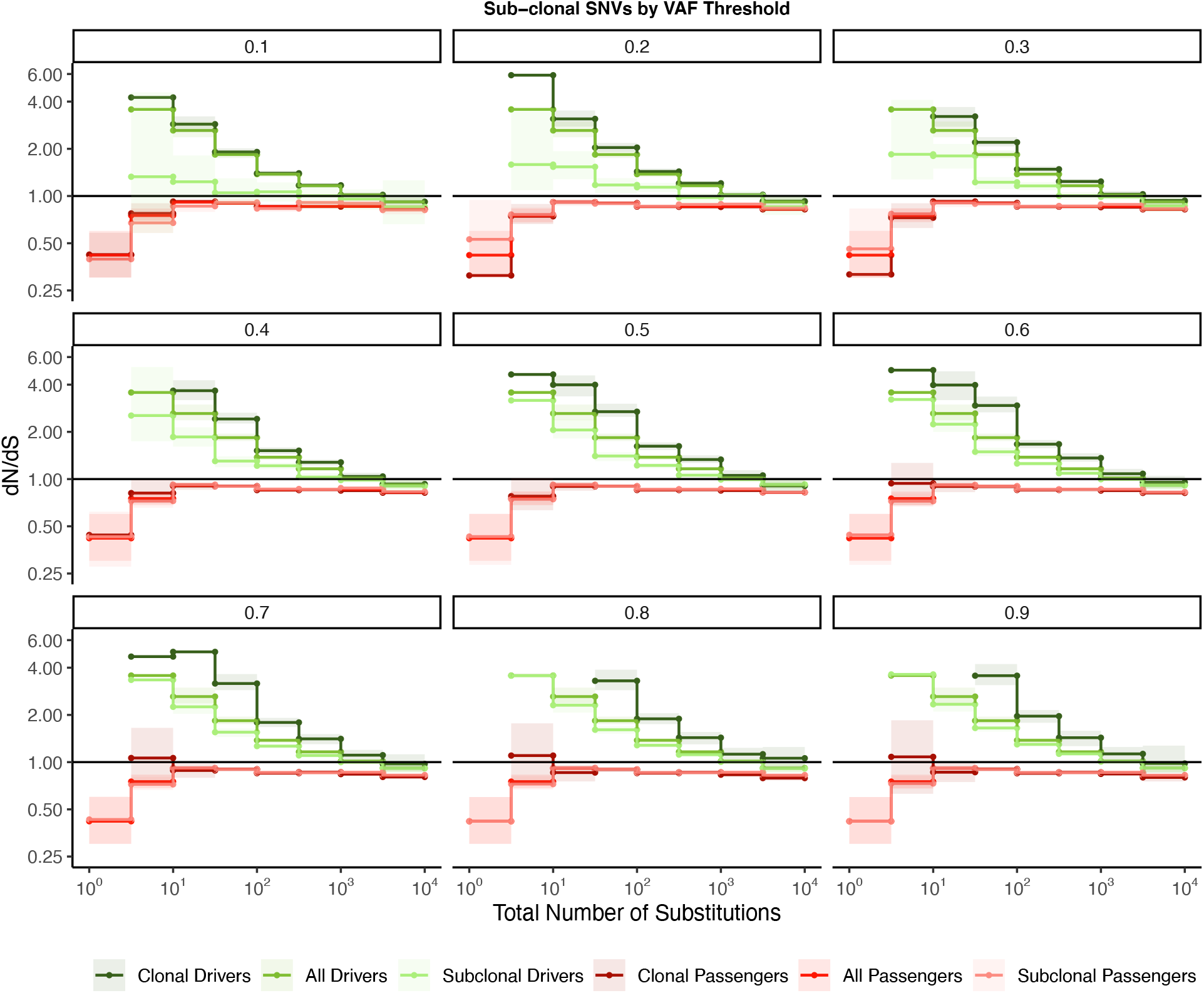
Signal of negative selection in subclonal mutations are robust to VAF threshold. dN/dS calculations within clonal and subclonal passenger and driver gene sets within tumors in TCGA stratified by the total number of substitutions. Title of each graph corresponds to increasing VAF threshold value used to define ‘subclonal’ (e.g. mutations with a VAF > 0.2 are clonal; mutations with a VAF < 0.2 are subclonal). Darker colors denote clonal passengers and drivers, while lighter colors denote subclonal passengers and drivers. Solid line of 1 is shown of dN/dS values expected under neutrality. Error bars are 95% confidence intervals determined by bootstrap sampling.

**Supplemental Figure 13.**
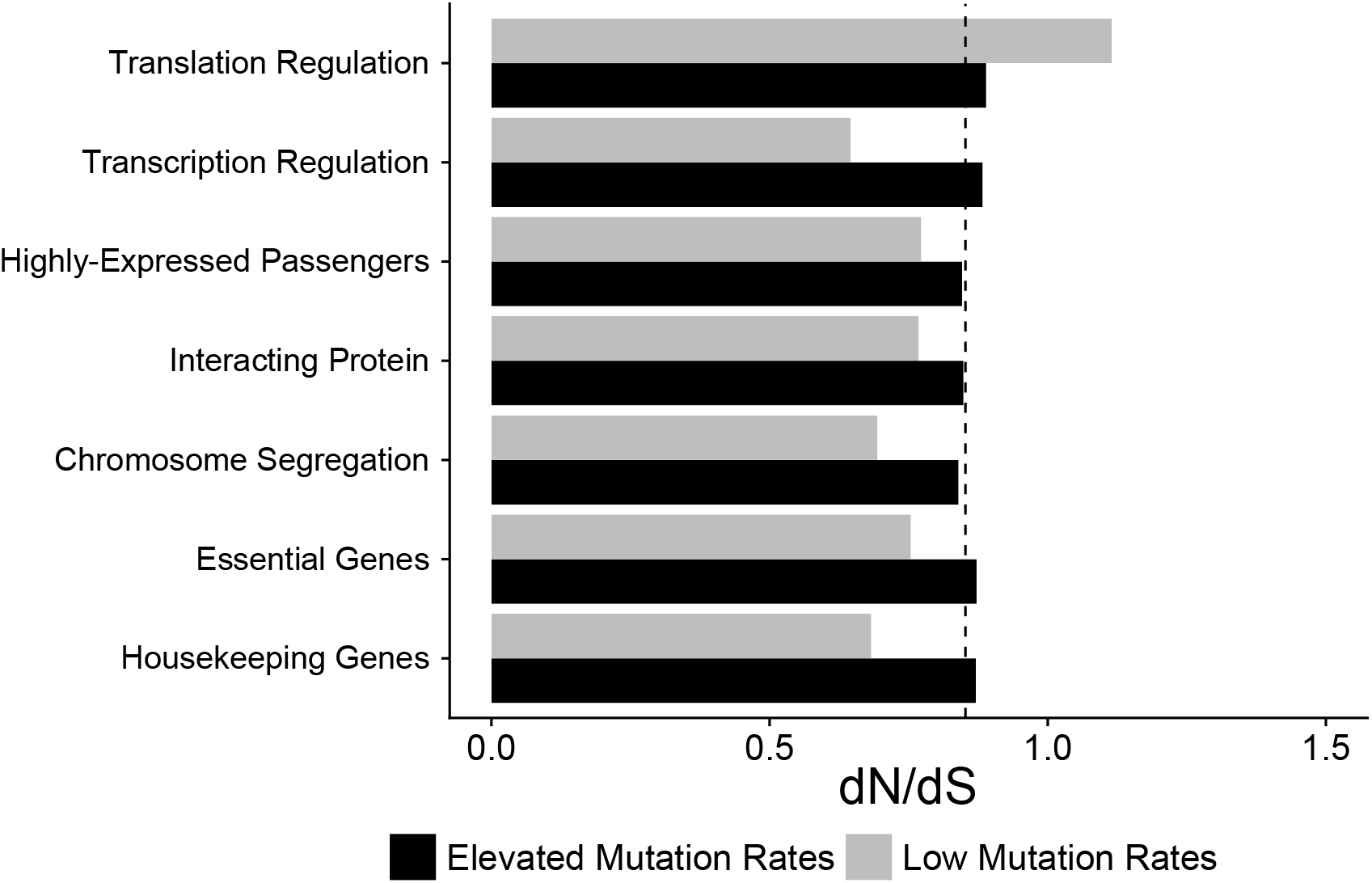
Attenuation of negative selection within different functional gene sets. dN/dS of passengers within different functional gene sets in low and high mutational burden tumors (*d*_N_ + *d*_S_ < 10 for low, grey; dn + ds > 10 for high, black). Both TCGA and ICGC genomic data were used. Dotted line denotes genomewide dN/dS of passengers for all mutation rates. Error bars are 95% confidence intervals determined by bootstrap sampling. Patterns of negative selection are not specific to any particular functional category (e.g. Essential or Housekeeping genes).

**Supplemental Figure 14.**
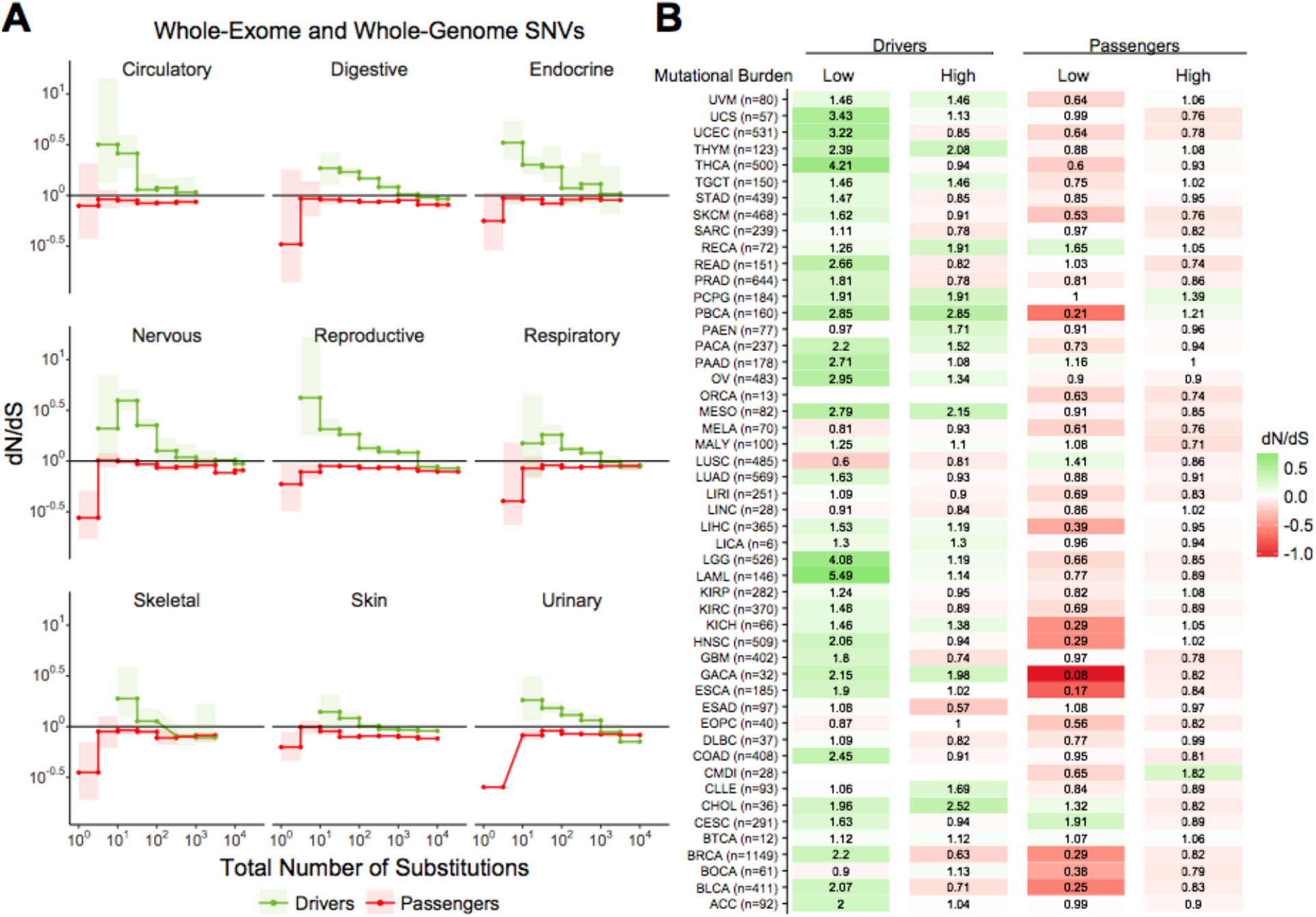
Attenuation of selection in SNVs persists across cancer subtypes and broad cancer group categories. **(A)** dN/dS in passenger and driver gene sets within tumors stratified by the total number of substitutions in broad tumor subcategories. Error bars are 95% confidence intervals determined by bootstrap sampling. **(B)** Log-scale heatmap of dN/dS values in passenger and driver gene sets of tumors stratified by the total number of substitutions within all 50 cancer subtypes in ICGC and TCGA. dN/dS of the lowest and highest mutational burden bin for each cancer subtype are shown.

**Supplemental Figure 15.**
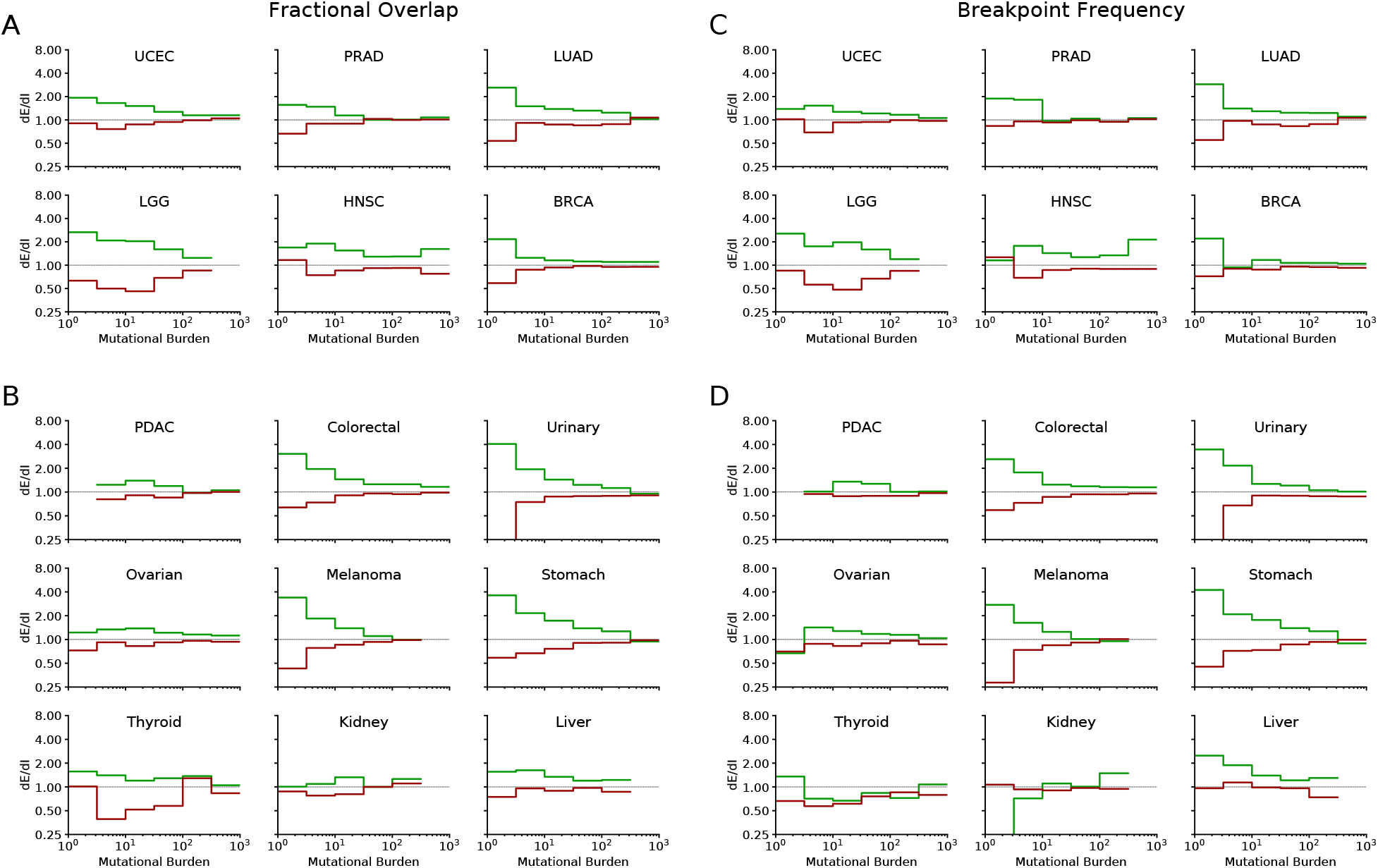
Attenuation of selection in CNAs is robust to cancer subtypes and broad cancer group categories. **(A)** Normalized fractional overlap (dE/dI) of driver (green) and passenger (red) Copy Number Alterations (CNAs) with the human exome for the six most commonly sequenced cancer subtypes (presented in Fig. 2). dE/dI > 1 suggests positive selection, while dE/dI < 1 suggests negative selection. Tumors are stratified by Mutational Burden (Total CNAs). **(B)** Same as in **(A)** for cancer subtypes with >200 genotyped samples that were not presented above (nine subtypes). **(C-D)** dE/dI of normalized breakpoint frequency stratified by Mutational Burden and segregated by cancer subtype. Subtype groupings are same as **(A-B)**. In general, both dE/dI measures exhibit positive selection on drivers that attenuates with mutational burden as well as negative selection on passengers that also attenuates with mutational burden across tumor subtypes. However, several exceptions are evident - especially for less-sequenced subtypes (bottom row of B & D).

**Supplemental Figure 16.**
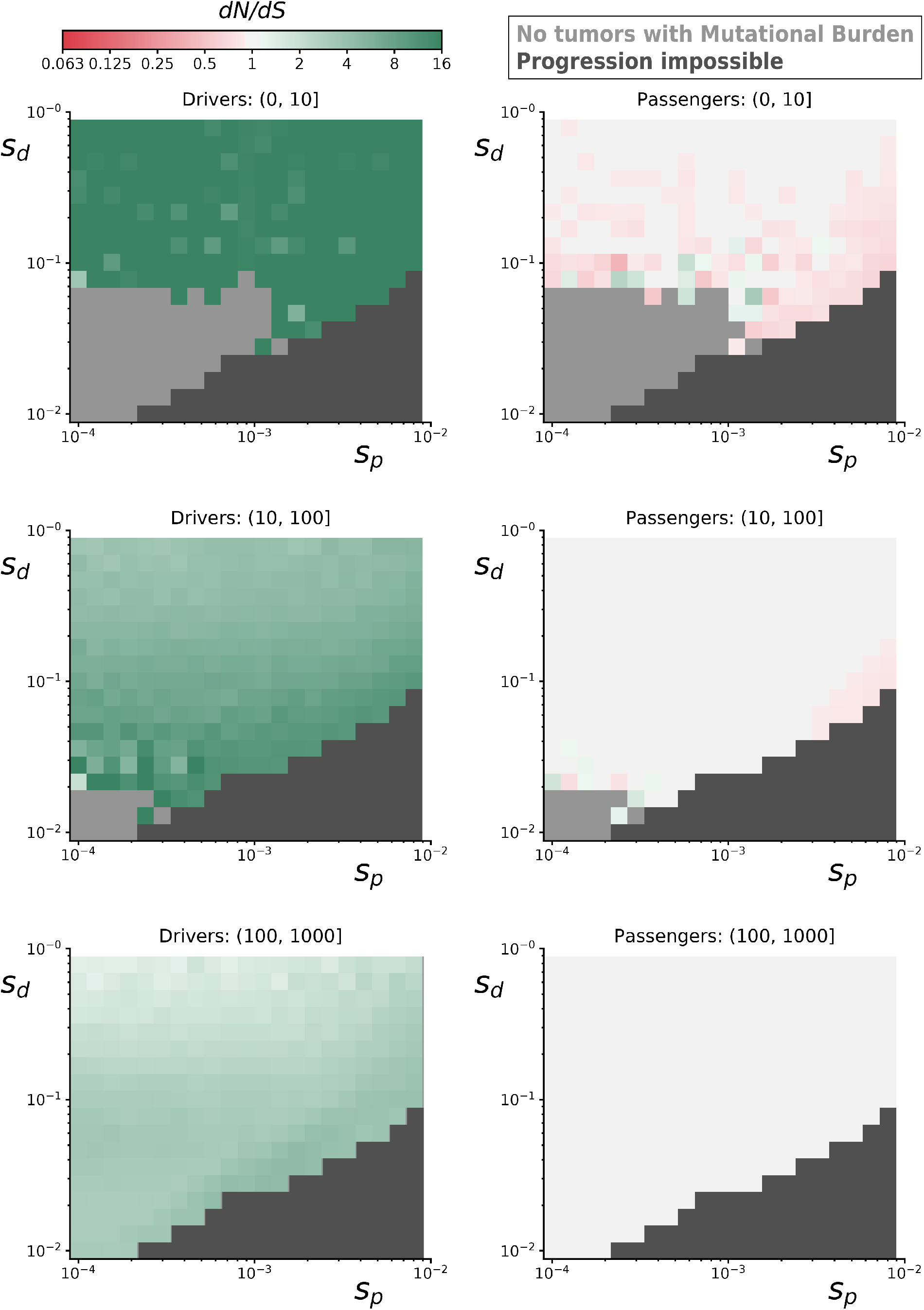
dN/dS rates of drivers and passengers in simulated cancers with various fitness coefficients. 10,000 simulated tumors were generated for various combinations of mean driver fitness benefits (*s*_drivers_) and mean passenger fitness costs (*s*_passengers_, Methods). For some parameter combinations, the combined fitness cost of passengers overwhelmed the fitness benefit of drivers and prevented cancer progression within 100 years (dark grey). dN/dS values of simulated mutations were calculated for drivers (left) and passengers (right) at various mutational burden (Total number of nonysnonymous and synonymous mutations). Top row is a mutational burden of 1 - 10; middle row is 11 - 100, and bottom row is 100 - 1,000. Some parameter combinations did not produce any tumors with low mutational burdens (light grey). Across all parameters, positive selction on drivers and negative selection against passengers attenuates with mutational burden. Passengers exhibit minimial negative selection in general, despite a collective burden that often prevented tumor progression, because of strong Hill-Roberston interference in asexual populations.

**Supplementary Figure 17.**
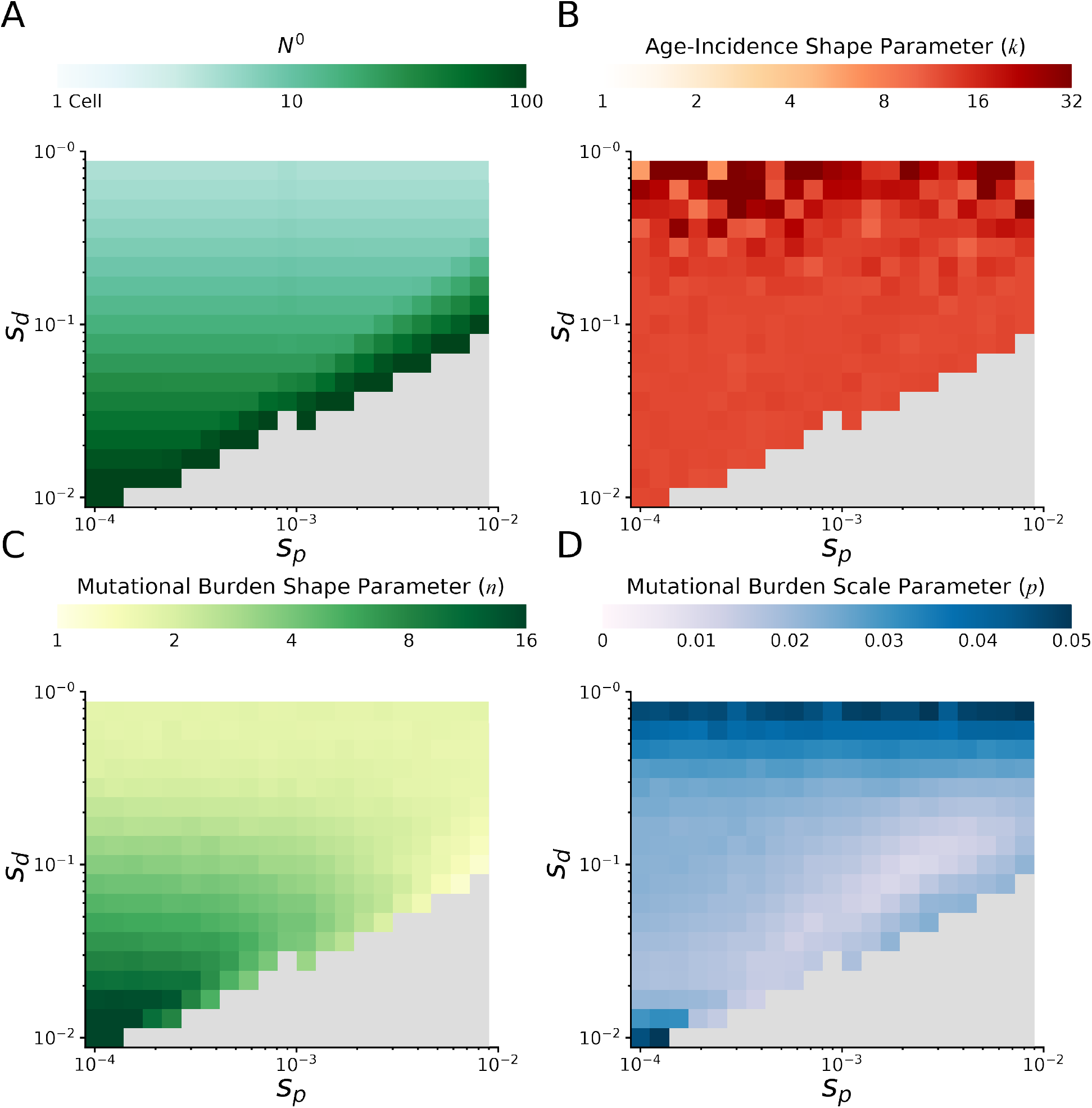
Probability of cancer by age and mutational burdens in simulated cancers at various fitness coefficients. Clinical summary statistics of simulated tumors at various combinations of mean driver fitness benefits (*s*_drivers_) and mean passenger fitness costs (*S*_p_, Methods). **(A)** Initial population size *N^0^* of simulated tumors. Initial population size approximates the equilibrium population size of a tumor following an initiating driver. Large population sizes are necessary for tumor progression when passenger deleteriousness is large compared to driver advantageousness - otherwise natural selection cannot drive carcinogenesis. Eventually, tumor progression is not possible for any reasonable initial population size (grey area). **(B)** MLE of Gamma distribution shape parameters describing the cancer age-inicidence rates of simulated tumors. A Gamma distribution of age-incidence is expected from the Armitage-Doll multistage model of tumorigenesis and describes human age-incidence rates well (Methods)^30^. Larger values correspond to a steeper increase in rate with age; human patient rates are ~5 pan-cancer. Scale parameter of the parametric fit is not informative because of a Gauge freedom in the model. **(C)** MLE of shape and **(D)** scale parameters of Negative Binomial distributions describing the mutational burdens of simulated tumors. Smaller values of shape parameter correspond to broader distributions of mutational burden; human tumors exhibit a value of ~2 pan-cancer. Smaller values of scale parameter correspond to a larger mean mutational burden; human tumors exhibit a value of ~1/50 (i.e. 50 passengers per rate-limiting driver).

**Supplementary Figure 18.**
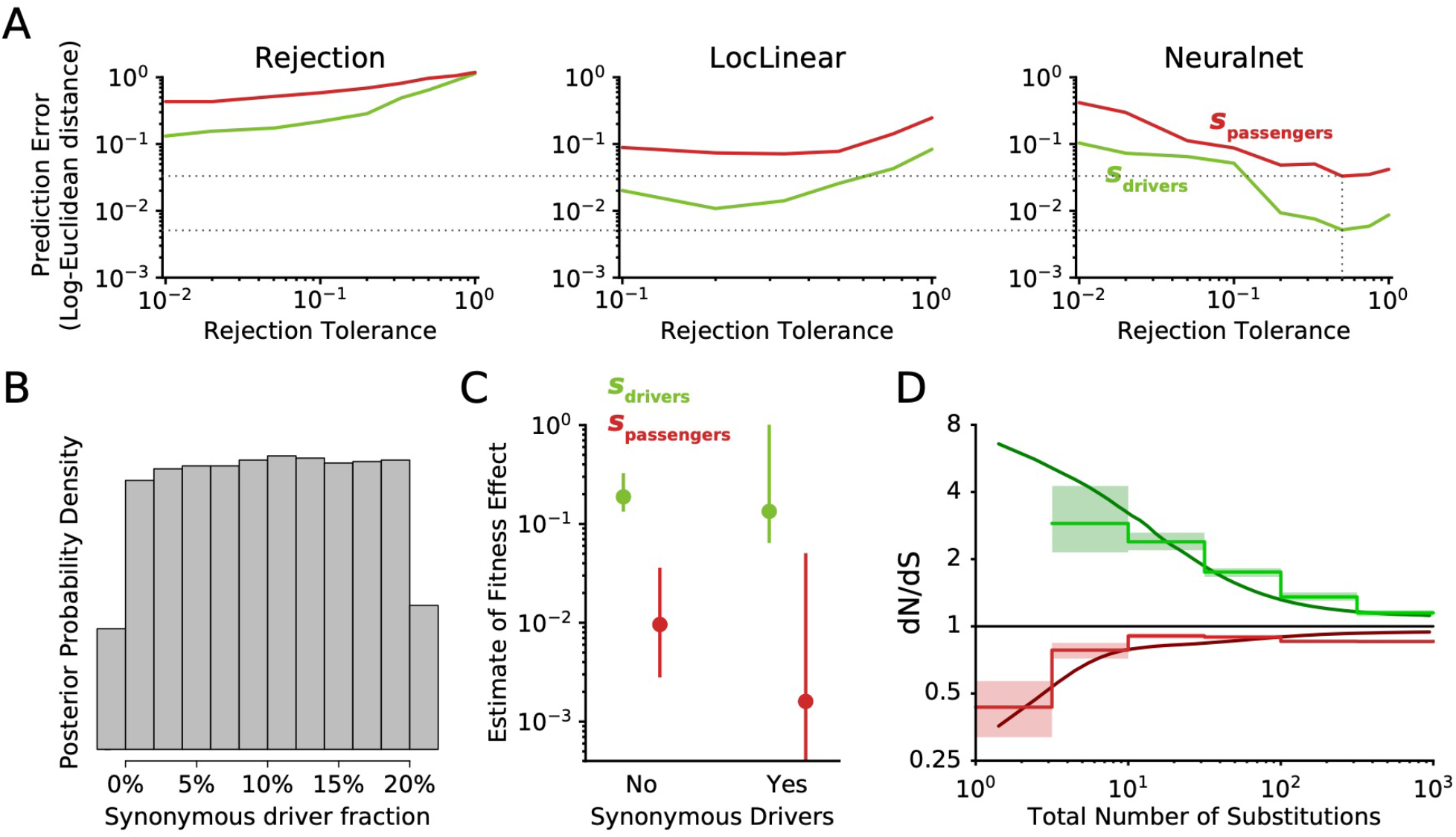
Implementation and use of ABC for model selection and parameter estimation. **(A)** Leave-one-out Cross Validation (CV) on the simulated data was used to select an optimal Rejection Tolerance and optimal rejection method. Observed data can be compared to simulated data using model rejection alone (left), or by comparing observed data to a (middle) local-linear regression or (right) Feed-Forward Neural Network single-layer model trained on the simulated data. In general, unsupervised training of a neural network on simulated data will often improve prediction accuracy by denoising stochasticity in the simulations (via kernel prediction.) A neural network with a rejection tolerance of 0.5 minimized prediction error of both driver and passenger fitness effects (illustrated by dotted lines) and was used to infer selection coefficients. This Cross Validation optimization procedure for ABC is advised^82^. **(B)** Posterior probability of models of tumor evolution incorporating synonymous drivers. The prior distribution of synonymous driver fractions (uniform from 0% to 20%) is nearly-identical to this posterior distribution. This suggests that nearly all models incorporating synonymous drivers can explained observed dN/dS patterns with the right combination of fitness parameters. **(C)** Posterior distribution of fitness effect of driver fitness benefits (*s*_drivers_) and passenger fitness costs (*s*_passengers_) after synonymous drivers are incorporated. MLE (circles) and 95% Confidence Intervals (lines) are reported. Similar to (B), incorporation of synonymous drivers undermines the ability of ABC to accurately infer fitness coefficients. **(D)** Comparison of dN/dS rates from one million simulated tumors (using ML estimates of *s*_drivers_ and *S*_passengers_, dark smooth lines) to observed dN/dS patterns (light, stepped lines). Both observed and simulated dN/dS rates of passengers rapidly approach 1 as mutation burden increases. This is presumably because, for populations near mutation-selection balance, the size of the fittest class of cells declines exponentially with the mutation rate (discussed in ^33^).

**Supplementary Figure 19.**
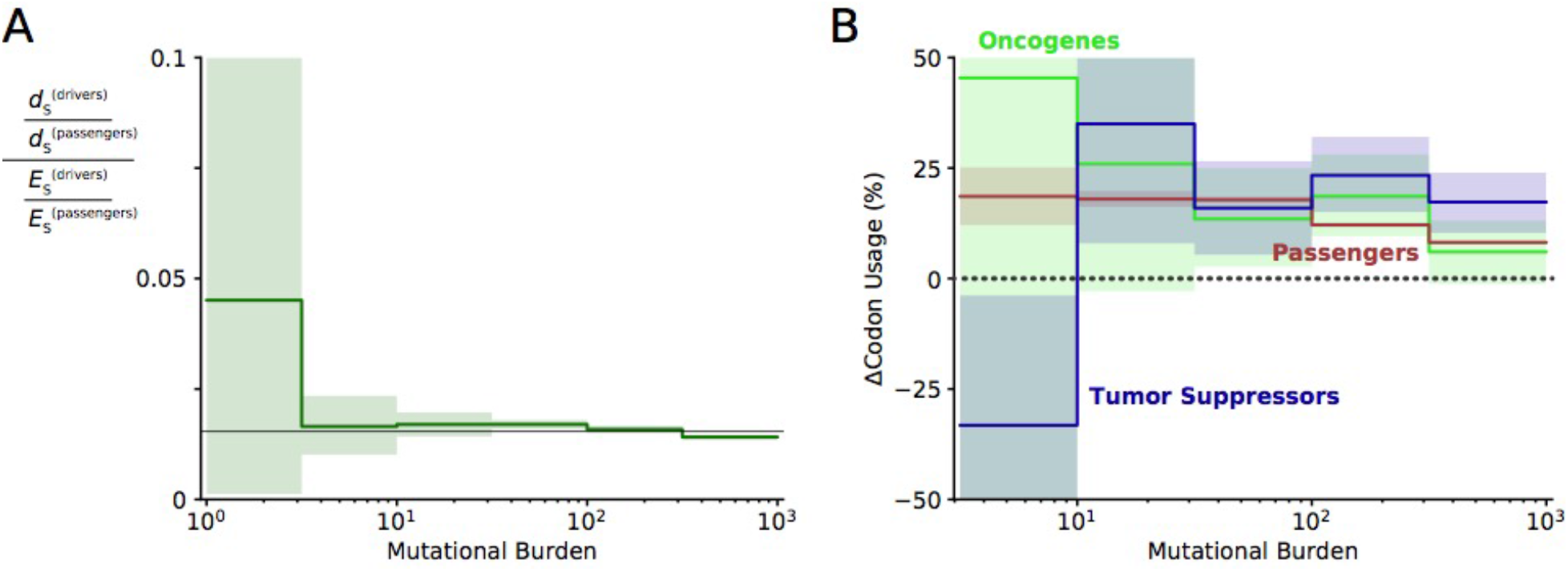
Evidence of positive selection on synonymous mutations within driver genes at low mutational burdens. (A) The quantity of synonymous mutations within driver genes was compared to the quantity of synonymous mutations within passenger genes and both were normalized by their expected frequencies using dNdScv. Black line denotes the genome-wide ratio of synonymous drivers to synonymous passengers (~2%, i.e. driver genes are ~2% of the human coding genome). At low mutational burdens, a non-significant increase in the quantity of synonymous drivers is observed, suggestive of positive selection for these mutations. **(B)** The change in codon usage imparted by all synonymous mutations was calculated for oncogenes, tumor suppressors, and passenger genes. Bias in codon usage suggests a functional effect of synonymous mutations. Increase in codon usage is expected to increase translational efficiency and increase protein abundance. Oncogenes are expected to exhibit positive selection for increased codon usage and exhibit a non-significant increase as mutational burden declines - consistent with positive selection for synonymous mutations within oncogenic drivers that is attenuated by Hill-Robertson interference. Similarly, tumor suppressors are expected to exhibit a decrease in codon usage at low mutational burdens, which is indeed significant (*p* = 0.03) presumably because there are more annotated tumor suppressor genes.

**Supplementary Figure 20.**
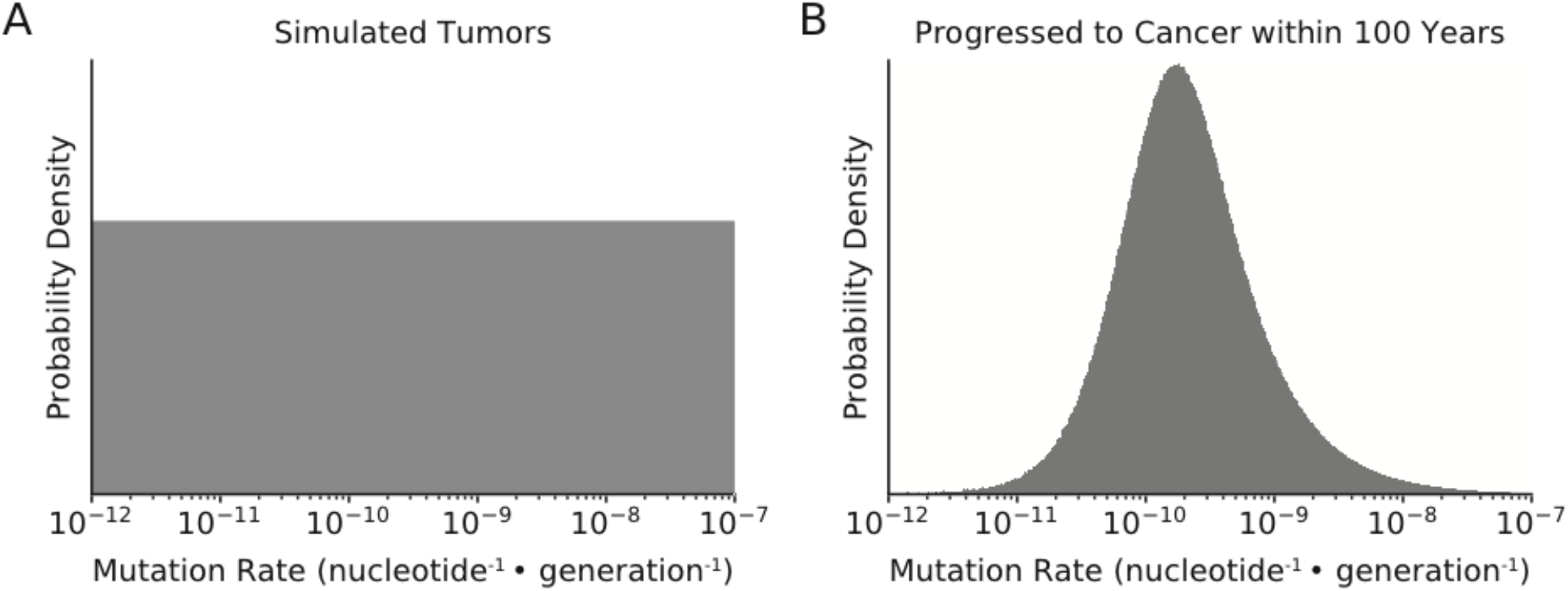
Distribution of Mutation Rates of simulated tumors. **(A)** Mutation rates of all simulated tumors were randomly-sampled from a uniform distribution (in log-space) from 10^−12^ to 10^−7^ nucleotide^−1^ • generation^-1^. **(B)** In simulations that best agreed with observed data (MLE of *s*_drivers_ = 18.8%, *s*_passengers_ = 0.96%), only tumors with intermediate mutation rates progressed to cancer within 100 years. Tumors with lower mutation rates do not progress to cancer within the 100-year time constraint of simulations, while tumors with exceptionally high mutation rates collapse via mutational meltdown.

**Supplemental Figure 21.**
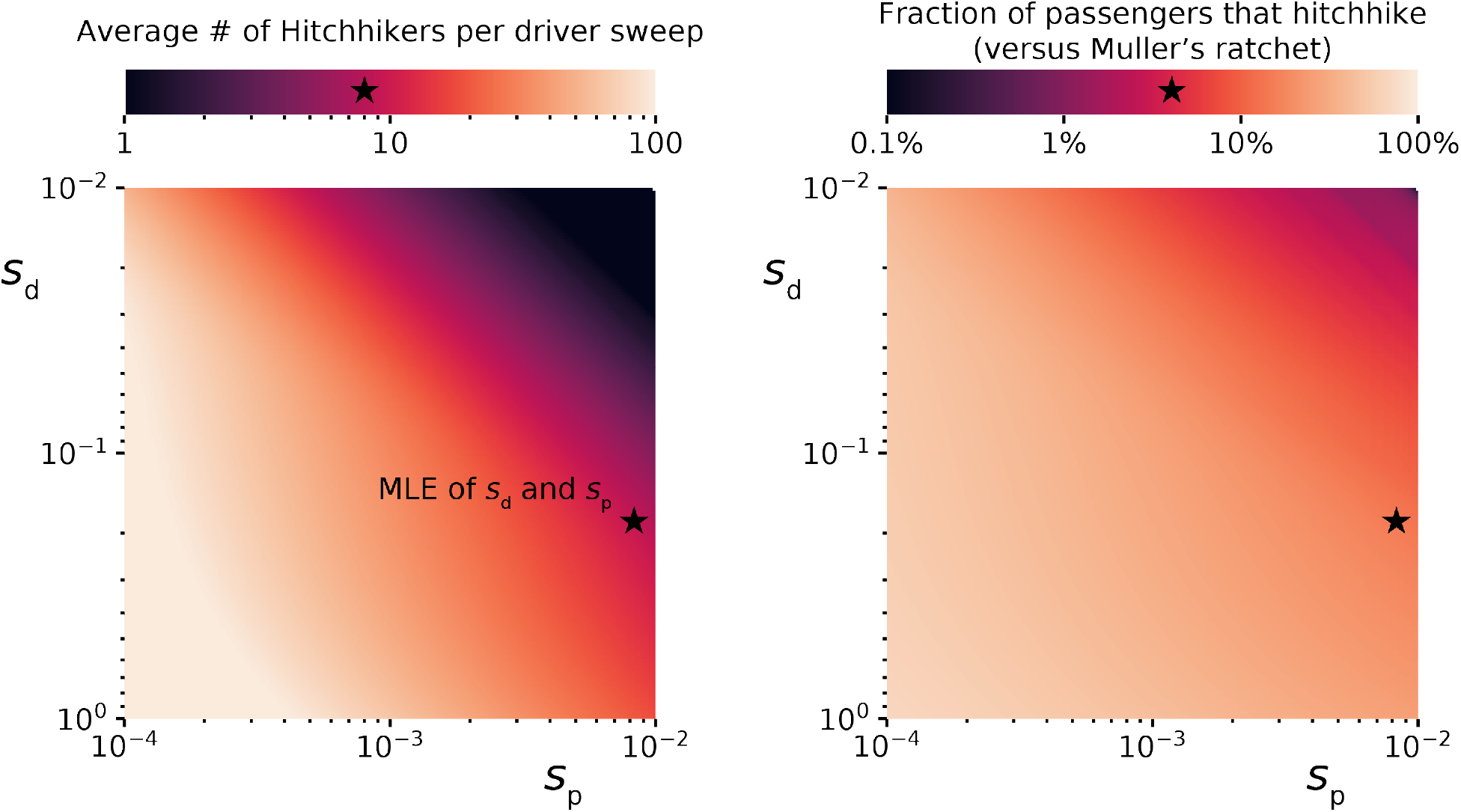
Relative contribution of Genetic Hitchhiking and Muller’s Ratchet to fix deleterious passengers. Using analytical theory developed in ^7,33,85^, we can estimate the relative rates of genetic hitchhiking and Muller’s Ratchet in our pancancer model of tumor evolution. As the relative strength of driver alterations increase (*s*_drivers_) relative to the selective cost of passengers (*s*_passengers_), more passengers hitchhike with each driver sweep (left). This increases the relative contribution of observed passengers that accumulate via hitchhiking (right). Using the Maximum Likelihood Estimates (MLE) of selection for drivers and against passengers, we estimate that an average of 8 passengers hitchhike with each driver, which account for 5.0% of accumulated passengers (the majority, and remainder, accumulate via Muller’s Ratchet).

**Supplemental Figure 22.**
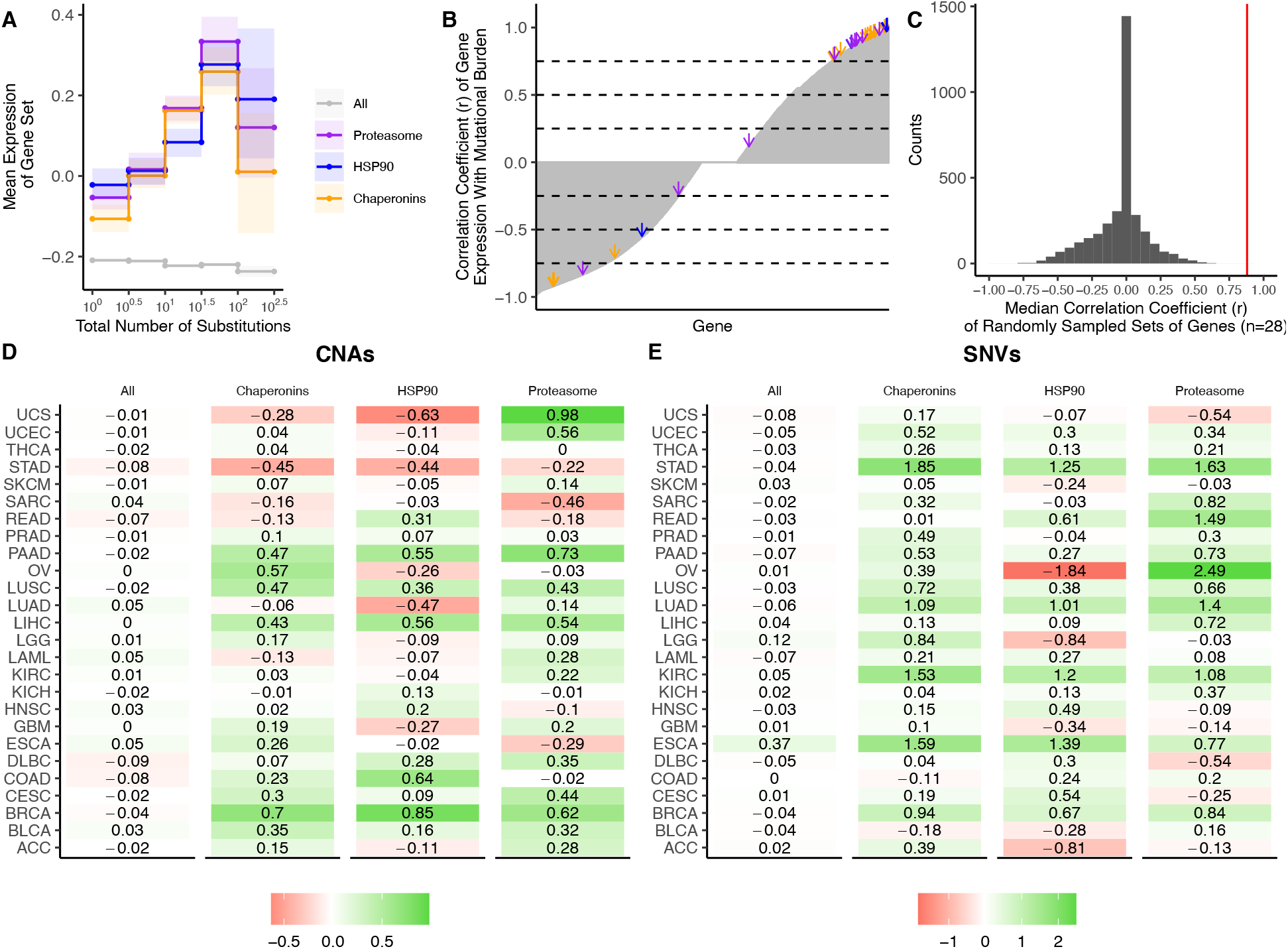
Upregulation of heat-shock protein pathways in tumors with elevated mutational burdens. **(A)** Z-scores of median gene expression of (i) all genes, (ii) HSP90, (iii) Chaperonins, and (iv) the Proteasome averaged across tumors stratified by the total number of CNAs. Expression of HSP90, Chaperonins, and Proteasome gene sets increases with the mutational burden of tumors (weighted *R*^2^ of 0.78, 0.87 and 0.84, respectively). Error bars are 95% confidence intervals determined by bootstrap sampling. **(B)** Correlation coefficients (*r*) of the expression of each gene in the genome (grey) in tumors stratified by the total number of substitutions. Shown in arrows are the correlation coefficients for HSP90 (blue), Chaperonins (orange), and the Proteasome (purple). Dashed lines in intervals of 0.25 are for viewing purposes only. **(C)** Median correlation coefficients of 10 million randomly sampled gene sets of the same size as HSP90, Chaperonins and the Proteasome (n=28) in grey. Red line denotes the median correlation coefficients of HSP90, Chaperonins, and the Proteasome (0.88). None of the randomly sampled gene sets have a higher median correlation coefficient than the observed value (0.88.) **(D-E)** Log-scale heatmap of changes in the Z-scores of median gene expression values of gene sets in for tumors stratified by the total number of substitutions **(D)** or CNAs **(E)** for cancer subtypes in TCGA. Changes in the mean gene expression of all genes, HSP90, Chaperonins, and Proteasome gene sets in the lowest and highest mutational burden bin for each cancer subtype are shown. Colors denote whether changes in gene expression from low mutational burden bins to high mutational burden bins are positive (green) or negative (red). Expression of HSP90, Chaperonins, and Proteasome gene sets increases with the mutational burden of tumors across cancer types stratified by the number of SNVs (*p* > 0.05, *p* < 6 × 10^−4^, *p* < 3 × 10^−3^ respectively; Wilcoxon signed-rank test) and CNAs (*p* > 0.05, *p* < 2 × 10^−2^, *p* < 1.5 × 10^−2^ respectively; Wilcoxon signed-rank test).

**Supplemental Figure 23.**
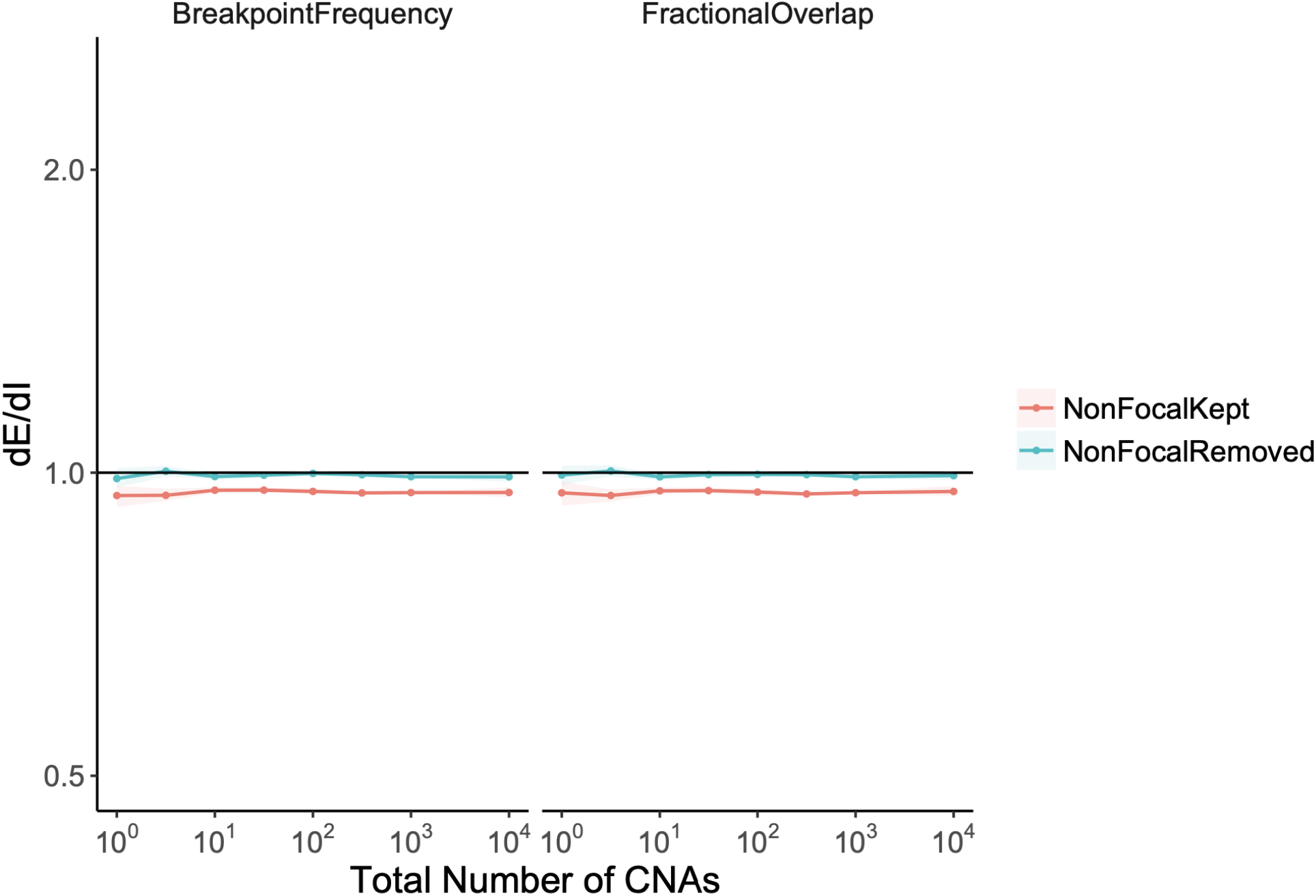
Random permutations of the positions of observed CNAs exhibit neutral values of dE/dI. The stop and start location of each observed CNA was randomly permuted, while preserving its length. dE/dI was calculated for CNAs (with and without non-focal amplifications) using both metrics: breakpoint frequency and fractional overlap. dE/dI values of random permutations are approximately 1, as expected for CNAs not experiencing selection.

**Supplementary Figure 24.**
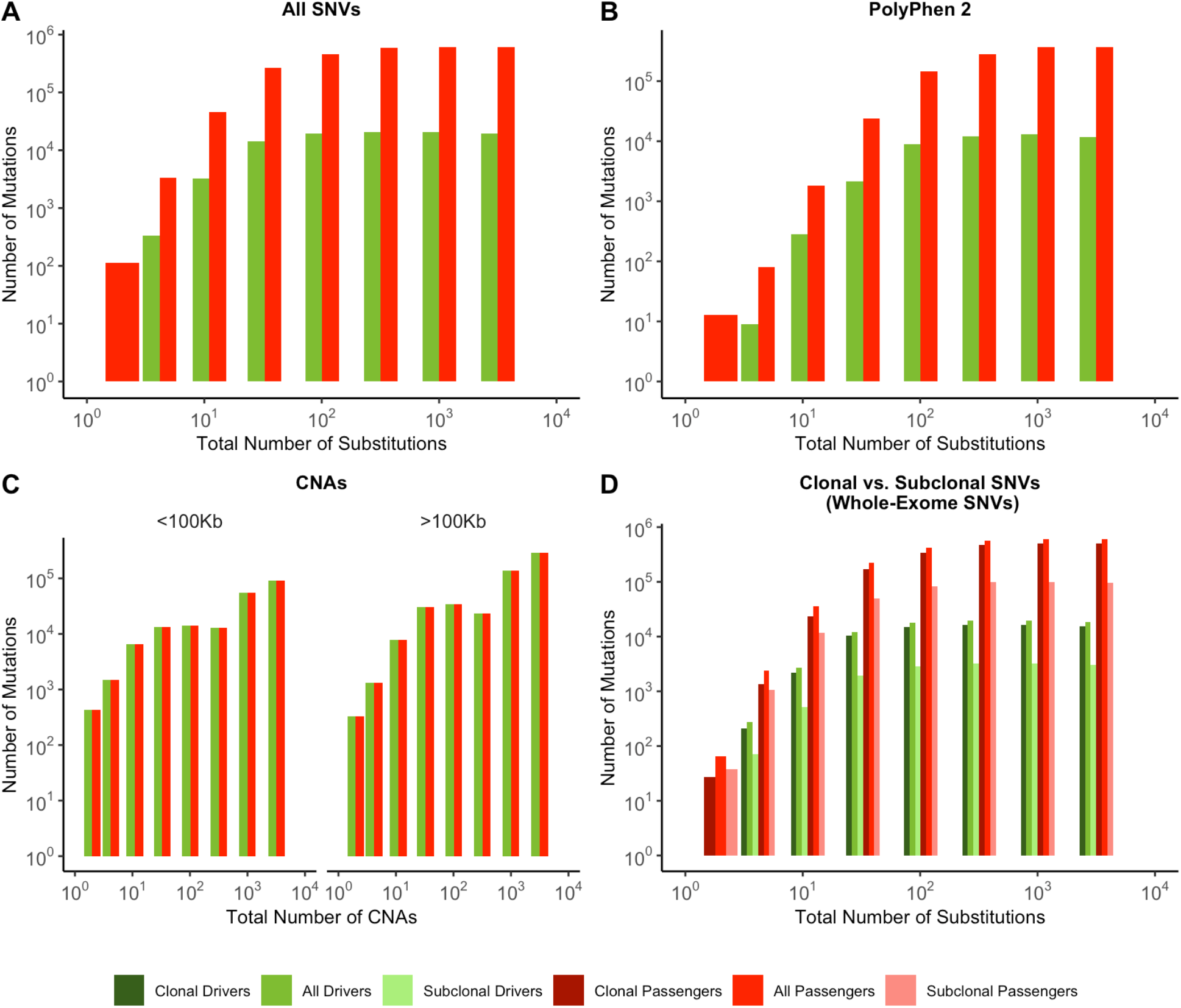
Quantity of mutations within each mutational burden bin for data depicted in Figure 2. **(A-D)** all report the total number of samples used in their respective figure pane within Figure 2. **(A)** Counts of mutations in passenger (red) and driver (green) gene sets within tumors stratified by the total number of substitutions in ICGC and TCGA. **(B)** Counts of the fraction of pathogenic missense mutations, annotated by PolyPhen2, in the same driver and passenger gene sets also stratified by total number of substitutions. **(C)** Counts of CNAs that reside within putative driver and passenger gene sets (identified by GISTIC 2.0, Methods) in tumors stratified by the total number of CNAs and separated by CNA length. **(D)** Counts of clonal (VAF > 0.2; darker colors) and subclonal (VAF < 0.2; lighter colors) passenger and driver gene sets in tumors stratified by the total number of substitutions.

**Supplementary Figure 25.**
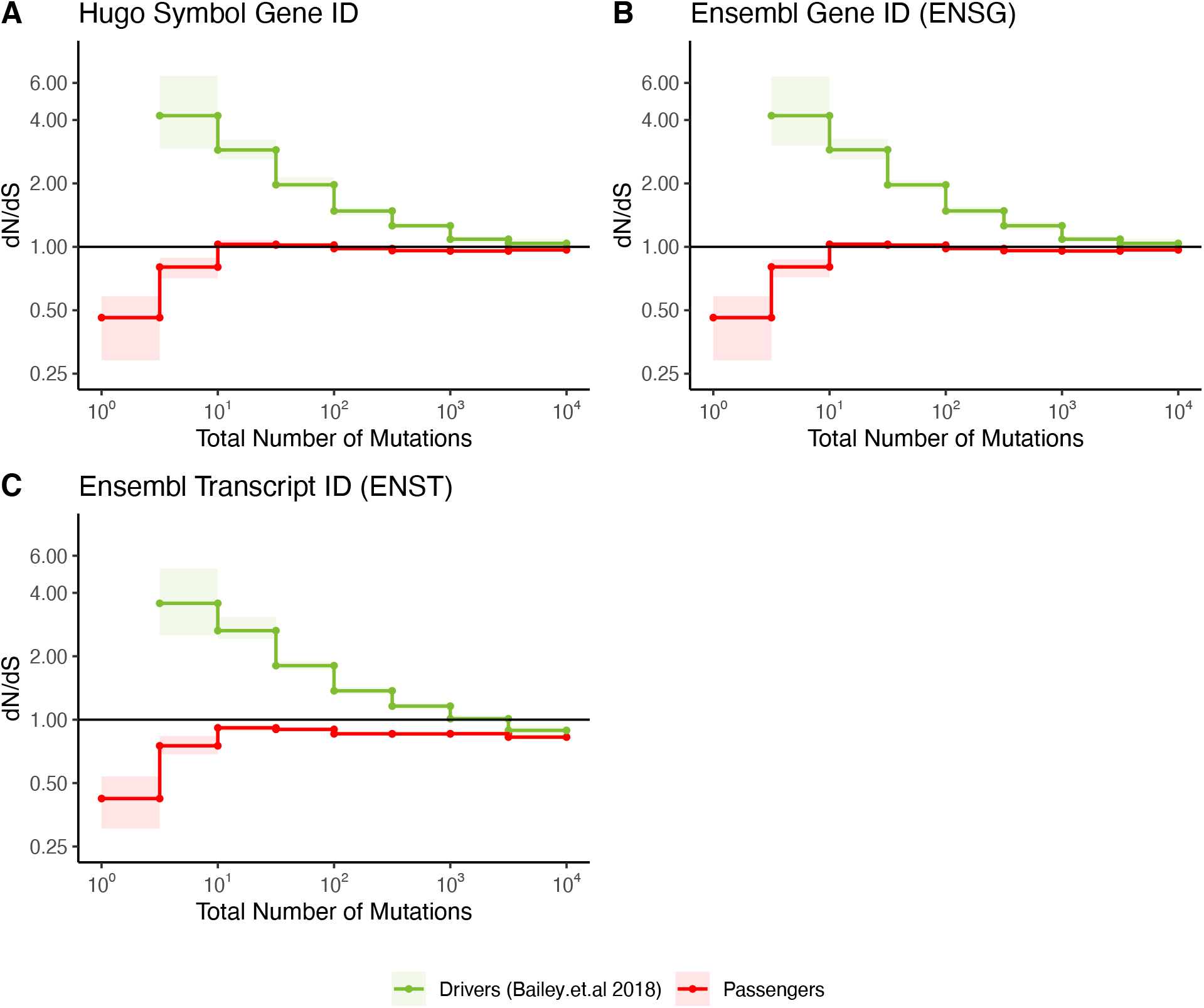
Patterns of selection when permuting gene sequences at the transcript or gene level. All panels show dN/dS of passenger and driver genes in tumors stratified by mutational burden within ICGC and TCGA datasets. **(A-B)** Gene-level sequences, annotated by Hugo symbols or Ensembl gene IDs, are used to permute the tri-nucleotide context of a mutation under the null model of mutagenesis. **(C)** Transcript level gene sequences, annotated by Ensembl, are used to permute the tri-nucleotide context of a mutation under our null model of mutagenesis. The solid line of 1 denotes dN/dS values expected under neutrality. Error bars (shaded area) represent 95% confidence intervals determined by bootstrap sampling.

### Supplementary Tables

**Table S1.** Broad (meta-categories) of cancer subtypes.

| **BROAD CATEGORY (N)** | **GDC TUMOR SUBTYPES IN GROUP** |
| --- | --- |
| Circulatory (371) | LAML, DLBC, CLLE, CMDI, MALY |
| Endocrine (925) | ACC, THYM, THCA, PAEN, PCPG |
| Urinary (1199) | BLCA, KICH, KIRC, RECA |
| Nervous (1059) | LGG, GBM, PBCA |
| Reproductive (3328) | BRCA, CESC, EOPC, OV, PRAD, UCEC, TGCT, UCS |
| Respiratory (1557) | LUSC, LUAD, HNSC |
| Skeletal (378) | SARC, BOCA, MESO |
| Digestive (2181) | ORCA, LIRI, PAAD, STAD, READ, CHOL, COAD, ESCA, GACA, LINC, ESAD, BTCA, LIHC |
| Skin (614) | UVM, SKCM, MELA |

**Table S2.**
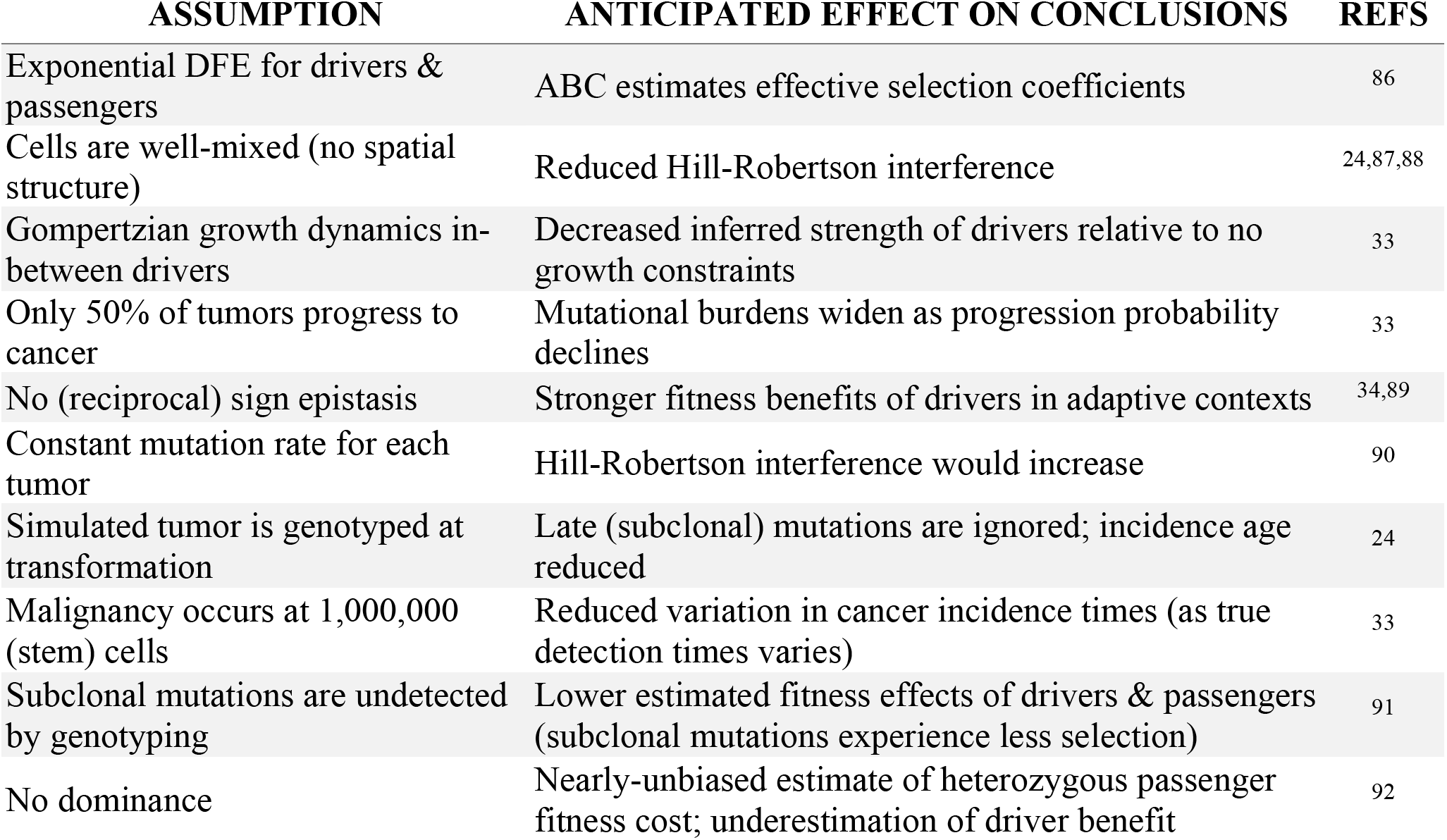
Assumptions of model of tumor evolution and anticipated effects

## References

1. Martincorena, I. et al. Universal Patterns of Selection in Cancer and Somatic Tissues. Cell 171, 1029–1041.e21 (2017).

2. Weghorn, D. & Sunyaev, S. Bayesian inference of negative and positive selection in human cancers. Nat. Genet. 49, 1785–1788 (2017).

3. Hill, W. G. & Robertson, A. The effect of linkage on limits to artificial selection. Genet. Res. 8, 269–94 (1966).

4. Drummond, D. A. & Wilke, C. O. Mistranslation-induced protein misfolding as a dominant constraint on coding-sequence evolution. Cell 134, 341–52 (2008).

5. Lobkovsky, A. E., Wolf, Y. I. & Koonin, E. V. Universal distribution of protein evolution rates as a consequence of protein folding physics. Proc. Natl. Acad. Sci. 107, 2983–2988 (2010).

6. Johnson, T. Beneficial mutations, hitchhiking and the evolution of mutation rates in sexual populations. Genetics (1999).

7. Neher, R. a & Shraiman, B. I. Fluctuations of fitness distributions and the rate of Muller’s ratchet. Genetics 191, 1283–1293 (2012).

8. Zapata, L. et al. Negative selection in tumor genome evolution acts on essential cellular functions and the immunopeptidome. Genome Biol. 19, 67 (2018).

9. Alexandrov, L. B. & Stratton, M. R. Mutational signatures: the patterns of somatic mutations hidden in cancer genomes. Curr. Opin. Genet. Dev. 24, 52–60 (2014).

10. Haradhvala, N. J. et al. Mutational Strand Asymmetries in Cancer Genomes Reveal Mechanisms of DNA Damage and Repair. Cell 164, 538–49 (2016).

11. Ostrow, S. L., Barshir, R., DeGregori, J., Yeger-Lotem, E. & Hershberg, R. Cancer Evolution Is Associated with Pervasive Positive Selection on Globally Expressed Genes. PLoS Genet. 10, 16–20 (2014).

12. Ellrott, K. et al. Scalable Open Science Approach for Mutation Calling of Tumor Exomes Using Multiple Genomic Pipelines. Cell Syst. 6, 271–281.e7 (2018).

13. Cancer Genome Atlas Research Network et al. The Cancer Genome Atlas Pan-Cancer analysis project. Nat. Genet. 45, 1113–20 (2013).

14. Hudson, T. J. et al. International network of cancer genome projects. Nature 464, 993–998 (2010).

15. Forbes, S. a et al. The Catalogue of Somatic Mutations in Cancer (COSMIC). Curr Protoc Hum Genet Chapter 10, Unit 10.11 (2008).

16. Bailey, M. H. et al. Comprehensive Characterization of Cancer Driver Genes and Mutations. Cell 173, 371–385.e18 (2018).

17. Gonzalez-Perez, A. et al. IntOGen-mutations identifies cancer drivers across tumor types. Nat. Methods 10, 1081–1082 (2013).

18. Adzhubei, I. a et al. A method and server for predicting damaging missense mutations. Nat. Methods 7, 248–9 (2010).

19. Korbel, J. O. et al. Systematic prediction and validation of breakpoints associated with copy-number variants in the human genome. Proc. Natl. Acad. Sci. 104, 10110–10115 (2007).

20. Zack, T. I. et al. Pan-cancer patterns of somatic copy number alteration. Nat Genet 45, 1134–1140 (2013).

21. McGrail, D. J. et al. Proteome Instability Is a Therapeutic Vulnerability in Mismatch Repair-Deficient Cancer. Cancer Cell 37, 371–386.e12 (2020).

22. Santagata, S. et al. High levels of nuclear heat-shock factor 1 (HSF1) are associated with poor prognosis in breast cancer. Proc. Natl. Acad. Sci. 108, 18378–18383 (2011).

23. Messer, P. W. Measuring the Rates of Spontaneous Mutation From Deep and Large-Scale Polymorphism Data. Genetics 182, 1219–1232 (2009).

24. Sottoriva, A. et al. A Big Bang model of human colorectal tumor growth. Nat. Genet. 47, 209–216 (2015).

25. López, S. et al. Interplay between whole-genome doubling and the accumulation of deleterious alterations in cancer evolution. Nat. Genet. 52, 283–293 (2020).

26. Gene Ontology Consortium. The Gene Ontology (GO) database and informatics resource. Nucleic Acids Res. 32, 258D–261 (2004).

27. Grossman, R. L. et al. Toward a Shared Vision for Cancer Genomic Data. N. Engl. J. Med. 375, 1109–1112 (2016).

28. McFarland, C. D., Korolev, K. S., Kryukov, G. V, Sunyaev, S. R. & Mirny, L. a. Impact of deleterious passenger mutations on cancer progression. Proc. Natl. Acad. Sci. 110, 2910–2915 (2013).

29. National Cancer Institute, S. S. B. Cancer Incidence – Surveillance, Epidemiology, and End Results (SEER) Registries Research Data. Surveillance, Epidemiology, and End Results (SEER) Program (http://www.seer.cancer.gov). (2007).

30. Frank, S. A. Dynamics of cancer: Incidence, Inheritance, and Evolution. (2007).

31. Supek, F., Miñana, B., Valcárcel, J., Gabaldón, T. & Lehner, B. Synonymous Mutations Frequently Act as Driver Mutations in Human Cancers. Cell 156, 1324–1335 (2014).

32. Bozic, I. et al. Accumulation of driver and passenger mutations during tumor progression. Proc Natl Acad Sci USA 107, 18545–18550 (2010).

33. McFarland, C. D., Mirny, L. a & Korolev, K. S. Tug-of-war between driver and passenger mutations in cancer and other adaptive processes. Proc. Natl. Acad. Sci. 111, 15138–15143 (2014).

34. Rogers, Z. N. et al. Mapping the in vivo fitness landscape of lung adenocarcinoma tumor suppression in mice. Nat. Genet. 50, 483–486 (2018).

35. Sánchez-Rivera, F. J. et al. Rapid modelling of cooperating genetic events in cancer through somatic genome editing. Nature 516, 428–431 (2014).

36. Vermeulen, L. et al. Defining stem cell dynamics in models of intestinal tumor initiation. Science (80-.). 342, 995–998 (2013).

37. Hanahan, D. & Weinberg, R. A. The Hallmarks of Cancer. Cell 100, 57–70 (2000).

38. Camps, M., Herman, A., Loh, E. & Loeb, L. a. Genetic constraints on protein evolution. Crit Rev Biochem Mol Biol 42, 313–326 (2007).

39. Williams, B. R. et al. Aneuploidy Affects Proliferation and Spontaneous Immortalization in Mammalian Cells. Science (80-.). 322, 703–709 (2008).

40. Cassa, C. A. et al. Estimating the selective effects of heterozygous protein-truncating variants from human exome data. Nat. Genet. 49, 806–810 (2017).

41. Glaire, M. A. & Church, D. N. Hypermutated Colorectal Cancer and Neoantigen Load. Immunother. Gastrointest. Cancer 187–215 (2017).

42. Gorgoulis, V. G., Pefani, D. E., Pateras, I. S. & Trougakos, I. P. Integrating the DNA damage and protein stress responses during cancer development and treatment. J. Pathol. 246, 12–40 (2018).

43. Dai, C., Whitesell, L., Rogers, A. B. & Lindquist, S. Heat shock factor 1 is a powerful multifaceted modifier of carcinogenesis. Cell 130, 1005–1018 (2007).

44. Zhang, J. et al. International Cancer Genome Consortium Data Portal--a one-stop shop for cancer genomics data. Database 2011, bar026–bar026 (2011).

45. Carithers, L. J. & Moore, H. M. The Genotype-Tissue Expression (GTEx) Project. Biopreserv. Biobank. 13, 307–308 (2015).

46. Faltas, B. M. et al. Clonal evolution of chemotherapy-resistant urothelial carcinoma. Nat. Genet. 48, 1490–1499 (2016).

47. Jiao, Y. et al. Exome sequencing identifies frequent inactivating mutations in BAP1, ARID1A and PBRM1 in intrahepatic cholangiocarcinomas. Nat. Genet. 45, 1470–1473 (2013).

48. Puente, X. S. et al. Non-coding recurrent mutations in chronic lymphocytic leukaemia. Nature 526, 519–524 (2015).

49. Chen, K. et al. Mutational landscape of gastric adenocarcinoma in Chinese: Implications for prognosis and therapy. Proc. Natl. Acad. Sci. 112, 1107–1112 (2015).

50. Tirode, F. et al. Genomic Landscape of Ewing Sarcoma Defines an Aggressive Subtype with Co-Association of STAG2 and TP53 Mutations. Cancer Discov. 4, 1342–1353 (2014).

51. Li, M. et al. Whole-exome and targeted gene sequencing of gallbladder carcinoma identifies recurrent mutations in the ErbB pathway. Nat. Genet. 46, 872–876 (2014).

52. Pickering, C. R. et al. Integrative genomic characterization of oral squamous cell carcinoma identifies frequent somatic drivers. Cancer Discov. 3, 770–81 (2013).

53. Johnson, B. E. et al. Mutational Analysis Reveals the Origin and Therapy-Driven Evolution of Recurrent Glioma. Science (80-.). 343, 189–193 (2014).

54. Pilati, C. et al. Genomic Profiling of Hepatocellular Adenomas Reveals Recurrent FRK-Activating Mutations and the Mechanisms of Malignant Transformation. Cancer Cell 25, 428–441 (2014).

55. Oberg, J. A. et al. Implementation of next generation sequencing into pediatric hematology-oncology practice: moving beyond actionable alterations. Genome Med. 8, 133 (2016).

56. Chun, H.-J. E. et al. Genome-Wide Profiles of Extra-cranial Malignant Rhabdoid Tumors Reveal Heterogeneity and Dysregulated Developmental Pathways. Cancer Cell 29, 394–406 (2016).

57. Guo, G. et al. Whole-Exome Sequencing Reveals Frequent Genetic Alterations in BAP1, NF2, CDKN2A, and CUL1 in Malignant Pleural Mesothelioma. Cancer Res. 75, 264–269 (2015).

58. Ren, S. et al. Whole-genome and Transcriptome Sequencing of Prostate Cancer Identify New Genetic Alterations Driving Disease Progression. Eur. Urol. 73, 322–339 (2018).

59. Armenia, J. et al. The long tail of oncogenic drivers in prostate cancer. Nat. Genet. 50, 645–651 (2018).

60. Shern, J. F. et al. Comprehensive Genomic Analysis of Rhabdomyosarcoma Reveals a Landscape of Alterations Affecting a Common Genetic Axis in Fusion-Positive and Fusion-Negative Tumors. Cancer Discov. 4, 216–231 (2014).

61. George, J. et al. Comprehensive genomic profiles of small cell lung cancer. Nature 524, 47–53 (2015).

62. Petrini, I. et al. A specific missense mutation in GTF2I occurs at high frequency in thymic epithelial tumors. Nat. Genet. 46, 844–849 (2014).

63. Auton, A. et al. A global reference for human genetic variation. Nature 526, 68–74 (2015).

64. Wang, K., Li, M. & Hakonarson, H. ANNOVAR: functional annotation of genetic variants from high-throughput sequencing data. Nucleic Acids Res. 38, e164–e164 (2010).

65. Cibulskis, K. et al. Sensitive detection of somatic point mutations in impure and heterogeneous cancer samples. Nat. Biotechnol. 31, 213–219 (2013).

66. O’Brien, K. P., Remm, M. & Sonnhammer, E. L. L. Inparanoid: a comprehensive database of eukaryotic orthologs. Nucleic Acids Res. 33, D476–80 (2005).

67. Campbell, P. & Martincorena, I. dNdScv. Welcome Sanger Institute https://www.sanger.ac.uk/science/tools/dndscv (2017).

68. Mermel, C. H. et al. GISTIC2.0 facilitates sensitive and confident localization of the targets of focal somatic copy-number alteration in human cancers. Genome Biol 12, R41 (2011).

69. Carter, S. L. et al. Absolute quantification of somatic DNA alterations in human cancer. Nat. Biotechnol. 30, 413–421 (2012).

70. Kumar, R. D., Searleman, A. C., Swamidass, S. J., Griffith, O. L. & Bose, R. Statistically identifying tumor suppressors and oncogenes from pan-cancer genome-sequencing data. Bioinformatics btv430 (2015) doi:10.1093/bioinformatics/btv430.

71. Wang, T. et al. Identification and characterization of essential genes in the human genome. Science (80-.). 350, 1096–1101 (2015).

72. Eisenberg, E. & Levanon, E. Y. Human housekeeping genes, revisited. Trends Genet. 29, 569–74 (2013).

73. Calderone, A. & Cesareni, G. mentha: the interactome browser. EMBnet.journal 18, 128 (2012).

74. Ha, G. et al. TITAN: inference of copy number architectures in clonal cell populations from tumor whole-genome sequence data. Genome Res. 24, 1881–1893 (2014).

75. Kampinga, H. H. et al. Guidelines for the nomenclature of the human heat shock proteins. Cell Stress Chaperones 14, 105–111 (2009).

76. Tanaka, K. The proteasome: Overview of structure and functions. Proc. Japan Acad. Ser. B 85, 12–36 (2009).

77. Arjan G., J. Diminishing Returns from Mutation Supply Rate in Asexual Populations. Science (80-.). 283, 404–406 (1999).

78. Gibson, M. a. & Bruck, J. Efficient Exact Stochastic Simulation of Chemical Systems with Many Species and Many Channels. J Phys Chem A 104, 1876–1889 (2000).

79. Turajlic, S. et al. Whole genome sequencing of matched primary and metastatic acral melanomas. Genome Res. 22, 196–207 (2012).

80. Michor, F., Iwasa, Y., Lengauer, C. & Nowak, M. a. Dynamics of colorectal cancer. Semin. Cancer Biol. 15, 484–493 (2005).

81. Howlader, N. et al. SEER Cancer Stastistics Review 1975-2010. SEER Cancer Statistics Review, 1975–2010, National Cancer Institute. (2013).

82. Csilléry, K., François, O. & Blum, M. G. B. abc: an R package for approximate Bayesian computation (ABC). Methods Ecol. Evol. 3, 475–479 (2012).

83. Gelman, A. Parameterization and Bayesian Modeling. J. Am. Stat. Assoc. 99, 537–545 (2004).

84. Futreal, P. A. et al. A census of human cancer genes. Nat Rev Cancer 4, 177–183 (2004).

85. Bachtrog, D. & Gordo, I. Adaptive evolution of asexual populations under Muller’s ratchet. Evolution (N. Y). 58, 1403–1413 (2004).

86. Good, B. H., Rouzine, I. M., Balick, D. J., Hallatschek, O. & Desai, M. M. Distribution of fixed beneficial mutations and the rate of adaptation in asexual populations. Proc Natl AcadSci USA 109, 4950–4955 (2012).

87. Korolev, K. S. et al. Selective sweeps in growing microbial colonies. Phys Biol 9, 26008 (2012).

88. Erik A. Martens, Rumen Kostadinov, Carlo C. Maley, and O. H. et al. Spatial structure increases the waiting time for cancer. New J Phys 13, 1–25 (2012).

89. Krug, J. Adaptation in tunably rugged fitness landscapes: The Rough Mount Fuji Model. 1–33 (2014).

90. Goyal, S. et al. Dynamic mutation-selection balance as an evolutionary attractor. Genetics 191, 1309–1319 (2012).

91. McVean, G. A. & Charlesworth, B. The effects of Hill-Robertson interference between weakly selected mutations on patterns of molecular evolution and variation. Genetics 155, 929–44 (2000).

92. Whitlock, M. C. Fixation probability and time in subdivided populations. Genetics 164, 767–79 (2003).

